# Predicting the effects of SNPs on transcription factor binding affinity

**DOI:** 10.1101/581306

**Authors:** Sierra S Nishizaki, Natalie Ng, Shengcheng Dong, Cody Morterud, Colten Williams, Alan P Boyle

## Abstract

GWAS have revealed that 88% of disease associated SNPs reside in noncoding regions. However, noncoding SNPs remain understudied, partly because they are challenging to prioritize for experimental validation. To address this deficiency, we developed the SNP effect matrix pipeline (SEMpl). SEMpl estimates transcription factor binding affinity by observing differences in ChIP-seq signal intensity for SNPs within functional transcription factor binding sites genome-wide. By cataloging the effects of every possible mutation within the transcription factor binding site motif, SEMpl can predict the consequences of SNPs to transcription factor binding. This knowledge can be used to identify potential disease-causing regulatory loci.

## Background

To date, genome wide association studies (GWAS) have identified over 100,000 loci associated with over 200 human diseases and phenotypic traits [1, 2]. Though 95% of known single nucleotide polymorphisms (SNPs) and 88% of GWAS SNPs fall into noncoding regions of the genome, most genetics studies focus on mutations within coding regions [3, 4]. This large disparity in knowledge gained from big data initiatives is likely due to the more direct interpretability of genic variation even though noncoding variation is also strongly linked to human disease [5, 6]. Identifying noncoding mutations leading to gene misregulation is critical to fully understanding GWAS results and their impact on complex and polygenic disorders.

As noncoding GWAS variants are overwhelmingly abundant compared to coding variants, many methods have been developed to prioritize potentially disease-associated mutations in noncoding regions for further study [7]. Generally, these tools focus on known regulatory regions of the genome, relying on variant overlap with experimental annotations such as regions of open chromatin and transcription factor binding [8-10]. To date these computational prioritization tools have assisted in identifying a handful of causal disease mutations from GWAS [11, 12]. However, these tools have only shown up to a 50% concordance rate between predictions, highlighting the need for additional prioritization metrics [7]. One way to improve these predictions is to investigate additional regulatory features to better understand a variant’s mechanism of action.

Transcription factor binding sites (TFBSs) are a regulatory feature of particular interest as they make up 31% of GWAS SNPs, yet only comprise 8% of the genome [13]. Mutations in TFBSs influence transcription factor binding affinity, alter gene expression, and have been associated with multiple human diseases including cancer, type 2 diabetes, as well as with increased total cholesterol [14-19]. However, altering different bases within a TFBS have been found to confer different effects on transcription factor binding [20, 21]. This finding has been reflected in cases of human disease, where certain bases in a sequence motif are more correlated with an associated disease than others [22]. Currently, the effect of mutations in a TFBS is estimated using a position weight matrix (PWM), which denotes a transcription factor’s binding motif using *in silico* analyses to determine its predominant binding sequence using a competitive binding assay (Figure 1a) [23]. PWMs predict where a transcription factor may bind in the genome by acting as its most frequent binding sequence; however they may not recapitulate known binding activity and are not sufficient to predict which mutations within a motif may alter binding affinity [24]. Additionally, using PWMs to predict how a SNP may affect transcription factor binding can be challenging, as PWMs do not contain information on the potential direction of effect of a mutation.

**Figure 1.**
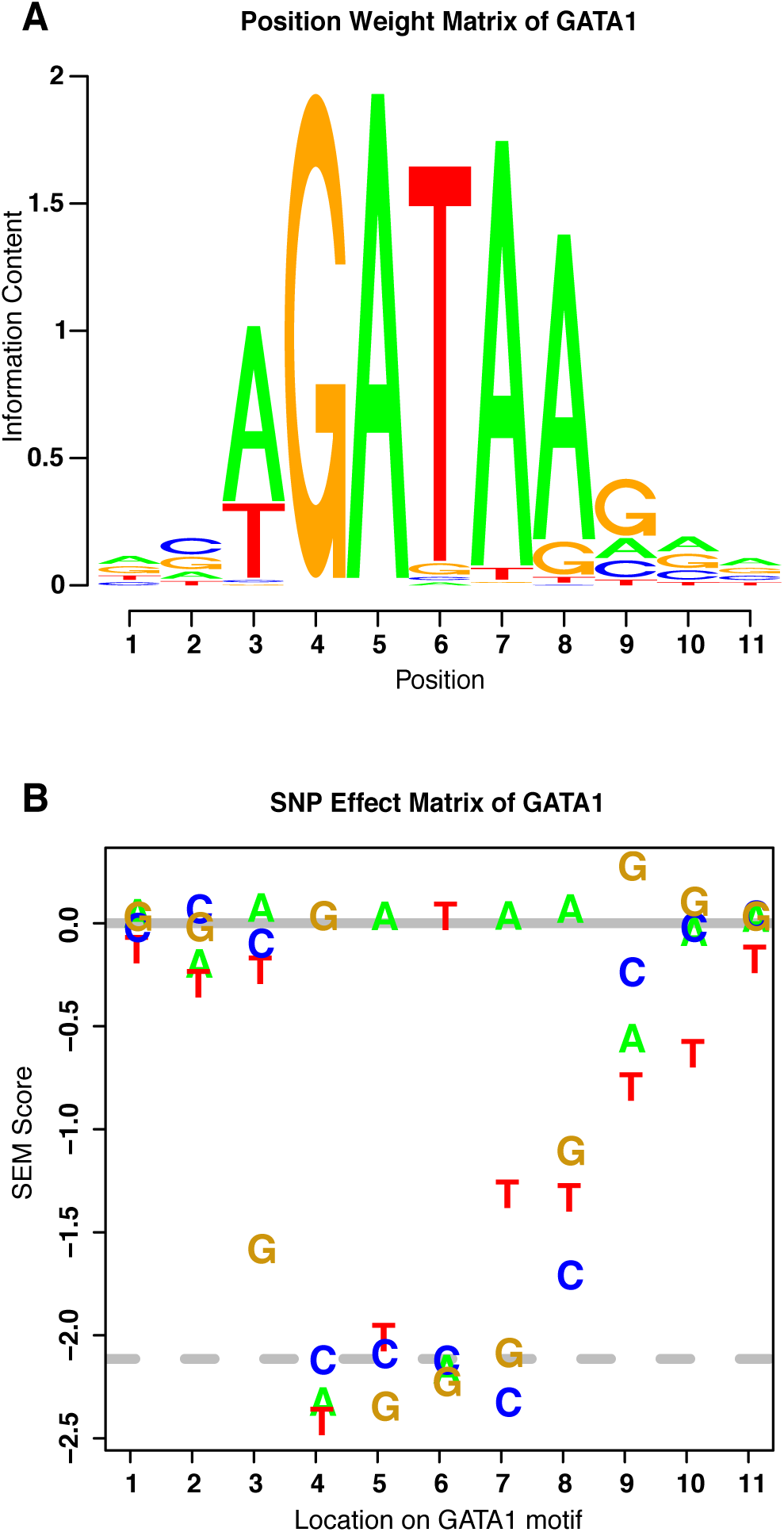
PWM v. SEM of transcription factor GATA1. A. The PWM can be read as likely nucleotides along a transcriptions factor’s motif. B. Similarly the SEM can be read as nucleotides along a motif, but with additional information about the effect any given SNP may have on transcription factor binding affinity. The solid grey line represents endogenous binding, the dashed grey line represents a scrambled background. We define anything above the solid grey line as predicted to increase binding on average, anything between the two lines as decreasing average binding, and anything falling below the dashed grey line as ablating binding on average.

While multiple tools have been developed to predict which mutations may lead to changes in binding affinity, all of these methods rely solely on information from PWMs and are thus subject to similar limitations [25-30]. More recently the Intragenomic Replicates (IGR) method was developed as a way to investigate FOXA1 involvement in breast cancer using GWAS data [31]. This method compares TFBSs containing putatively deleterious mutations to their wild-type counterparts using genome-wide chromatin immunoprecipitation followed by deep sequencing (ChIP-seq) data to estimate predicted changes to transcription factor binding affinity. The predictions generated by IGR were found to be highly correlated with ChIP-qPCR results and were successfully used to identify a risk allele associated with a fivefold change in gene expression in breast cancer. IGR represents a marked improvement over other methods due to its specific calibration of variants to ChIP-seq data, an endogenous source of transcription factor binding affinity information. Currently, IGR exists only as a method designed to probe individual mutations and must be reconstructed for each new mutation and transcription factor. However, the premise of using ChIP-seq data to predict transcription factor binding could be expanded to more quickly and accurately predict TBFS mutations.

In order to improve current methods to be applicable to a wide range of transcription factors and to better predict which mutations within TFBSs may lead to changes in binding affinity, we have developed a new method: the SNP effect matrix pipeline (SEMpl). Our method uses endogenous ChIP-seq data and existing variants genome-wide similar to the IGR method, however SEMpl also includes a catalogue of kmers separated by a single base change from a TFBS motif, allowing it to provide an estimate of the consequence of every possible mutation in a TFBS. We call these SNP effect matrices (SEMs, Figure 1). Here, we demonstrate that SEMs recapitulate known motifs, are robust to input data and cell type, and are better at predicting changes to transcription factor binding affinity than the current standard, PWMs. By developing SEM scores, we aim to improve the prioritization of noncoding GWAS variants for further experimental validation, expand the understanding of noncoding genomic variation, and further technology toward developing tools for personalized medicine.

## Results

### SNP Effect Matrix pipeline

SEMpl utilizes three types of experimental evidence to make its predictions: ChIP-seq data, which provides a transcription factor’s endogenous binding in the genome; DNase I hypersensitive sites sequencing (DNase-seq) data, which represents regions of open chromatin where transcription factors are known to function; and position weight matrices (PWMs), which denote previous knowledge of the binding pattern of transcription factors (Figure 1). We obtained ChIP-seq and DNase-seq data from the ENCODE project and PWMs from the JASPAR, Transfac, UniPROBE and Jolma databases [13, 29, 32, 33, 34].

SEMpl first enumerates a PWM of interest into a list of kmers using a permissive cutoff p-value threshold of 4^−5^ using the software TFM-Pvalue (Figure 2A)[35]. This first list of kmers, refered to as the endogenous kmer list, represents sequences where the transcription factor of interest has an increased likelihood of binding. To observe additional sequences which may show distinct binding preferences, SEMpl next takes the endogenous kmer list and simulates all possible SNPs *in silico* to create lists of mutated kmers (Figure 2B). For example, by changing all bases in position 6 to a G nucleotide in every kmer in the endogenous kmer list SEMpl creates a mutated kmer list for G in position 6. These lists of mutated kmers are then aligned to the hg19 human genome in regions of open chromatin, as determined by DNase-seq. The ChIP-seq score is then calculated as the highest signal value over the region 50bp before and after the aligned site (Figure 2C). Next the SEM score for each position is computed as the log2 of the average ChIP-seq signal to endogenous signal ratio for the mapped kmers for each mutated kmer list. Taken together, the SEM scores for each base form a matrix for each nucleotide at every position along the motif.

**Figure 2.**
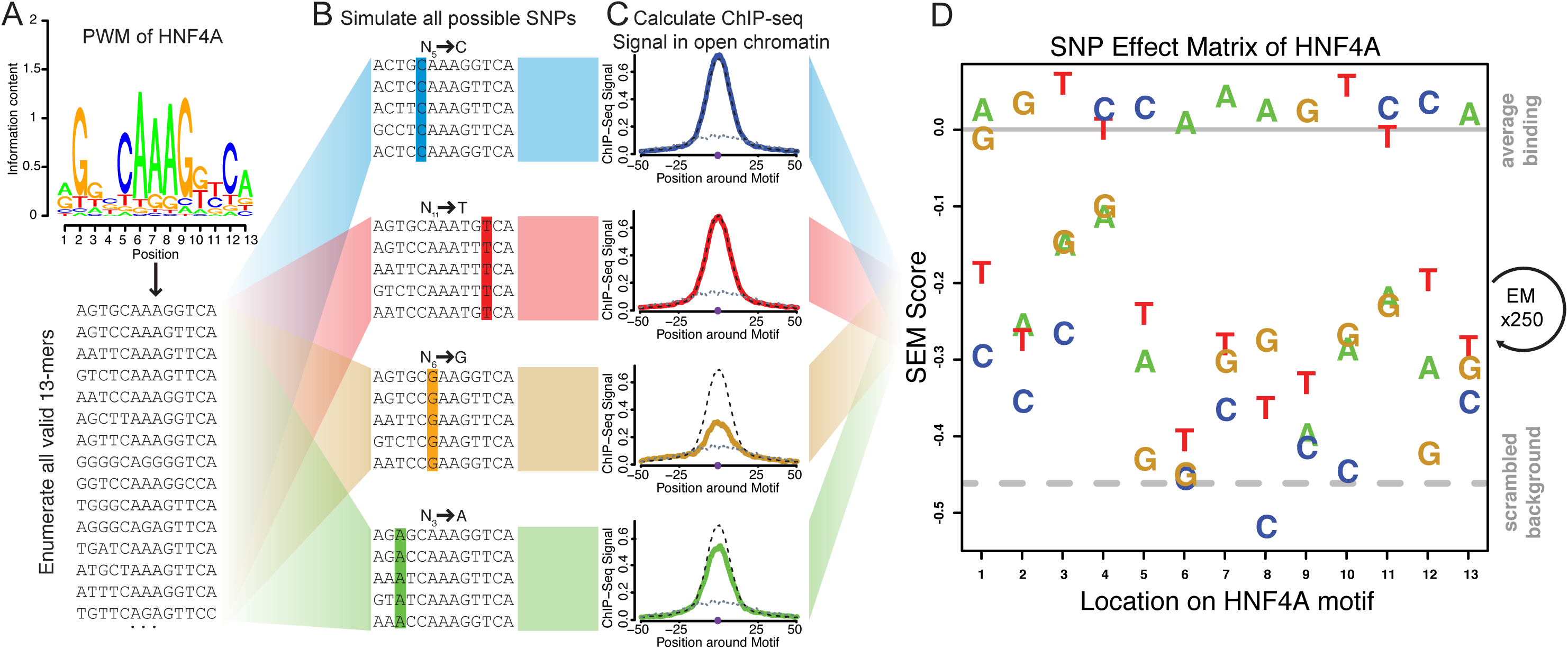
SEM Methods pipeline. A. All kmers with a PWM score below the TFM p-value are generated for a single transcription factor. B. All possible SNPs are introduced *in silico* for each kmer. C. All enumerated kmers are then aligned to the genome, and filtered for regions of open chromatin by DNase-seq. The average ChIP-seq scores are then calculated for each alignment (dash line represents endogenous binding, dotted line scrambled background). D. Final SEM scores are log2 transformed and normalized to the average binding score of the original kmers (solid grey line). A scrambled baseline, representing the binding score of randomly scrambled kmers of the same length is also added (dashed grey line). Once a SEM score is calculated, the output can be used to generate a new PWM. This iterative process can correct for disparities introduced by the use of different starting PWMs. The HepG2 cell line data was used for the ChIP-seq and DNase data for HNF4a.

The above process is repeated until convergence using an estimation maximization-like method (EM) (Figure 2D). To control for poor quality data and to identify background levels of binding, a final kmer list of randomly scrambled endogenous kmers is included to represent a random baseline where transcription factor binding would not be expected to occur (displayed as a dashed grey line on an SEM plot). We evaluate the final SEM by taking the average of 100 T-tests comparing a random sample of 1,000 endogenous binding sites with matched background sites. Finally, we define scores above 0 as predicted to increase binding on average, scores between 0 and the scrambled background as decreasing average binding on average, and scores falling below the scrambled background as ablating binding on average.

### SEM scores better recapitulate endogenous binding than PWMs

SEM scores are expected to be more representative of endogenous binding patterns than PWMs as these predictions are generated using an endogenous measure of genome-wide binding affinity. We demonstrate this by correlating SEM and PWM scores across full length kmers for transcription factor FOXA1 to their average ChIP-seq signals at corresponding sequences genome wide (Figure 3). When comparing predictions with experimentally generated binding affinity data above standard cutoffs (see methods), SEMs had a stronger correlation than PWMs (SEM: R^2^=0.66, PWM: R^2^=0.24), demonstrating our predictions represent a more robust measure of endogenous binding affinity. This pattern holds true when allowing a very lenient PWM cutoff of 11 (R^2^=0.28) as well as for the entire datasets (SEM: R^2^=0.19; PWM: R^2^=0.03).

**Figure 3.**
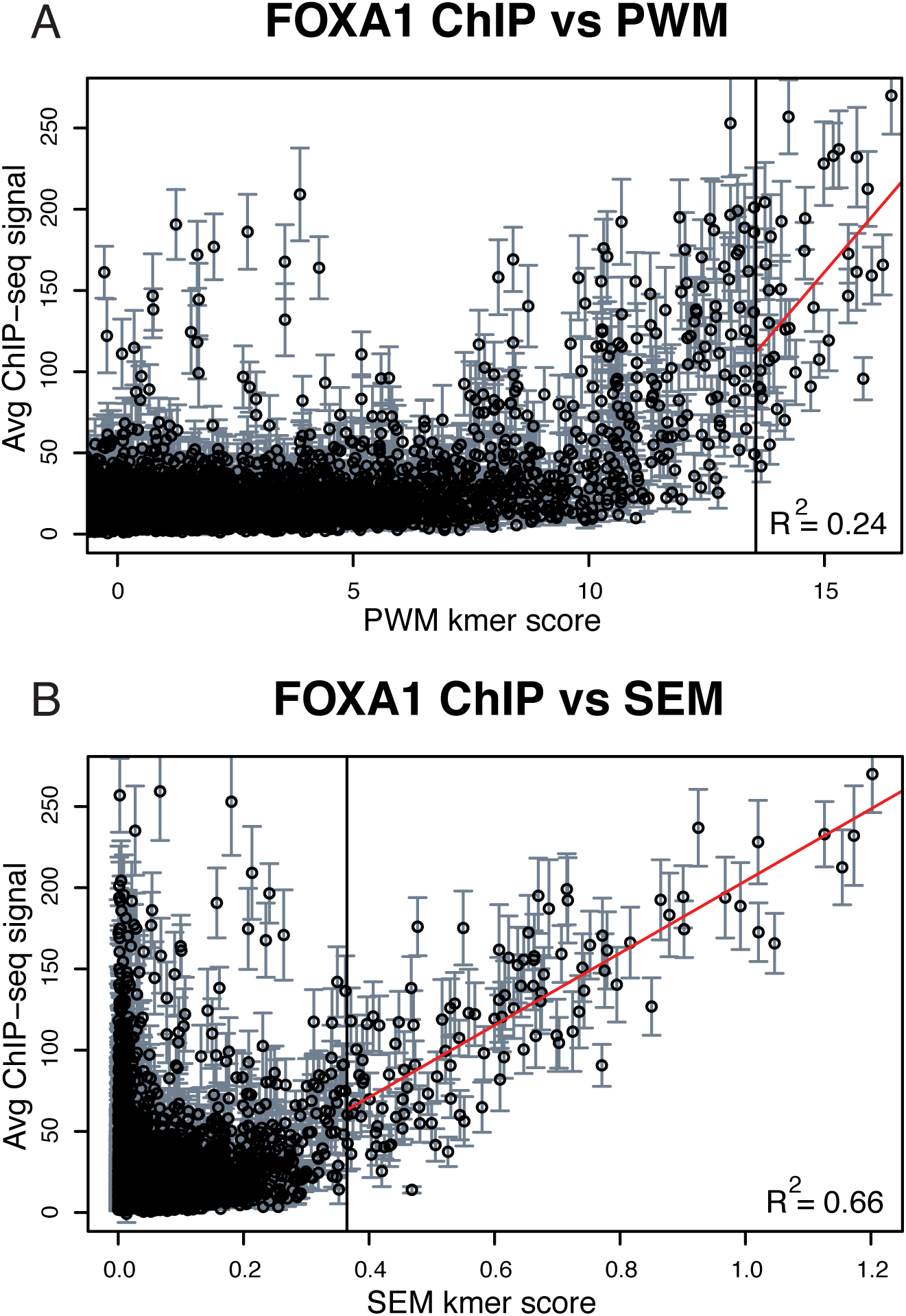
SEMs show a better correlation with whole kmer ChIP-seq signal (B, R^2^=0.66) than PWMs (A, R^2^=0.24). The line dividing the plot represents a standard cut off for PWM visualization (p-value = 4^−8^). Coefficient of determinations (R^2^) were calculated to the right of the vertical lines, representing the TFM p-value cutoff for PWMs and the average scrambled background cutoff for SEMs (0.36 for FOXA1). SEM values are displayed as 2^n^ for visualization purposes. PWM values only shown >0, a full plot can be found in Supplementary Figure 1.

These findings indicate that SEM plots better recapitulate known patterns of transcription factor binding beyond the information detailed in a PWM. Of note, there are cases where the PWM shows approximately equal information content for distinct bases sharing a position, yet the SEM plot reveals a wide margin of binding differences between the two bases fueled by differences in predicted direction of effect on binding affinity (i.e. position 3 or 10 of HNF4a in Figure 2).

### Ubiquitous Transcription Factors show cell type and dataset independence

To determine if SEM results show a dataset specific dependence, we evaluated the transcription factor FOXA1 using ChIP-seq data from two different ENCODE datasets gathered in the same HepG2 cell line (ENCFF658RGX; ENCFF898FCL) (Figure 3). We found nearly identical SEMpl outputs (p-value = 4.14e-56) using least squares regression analysis.

We next expanded this to investigate if SEM results were dependent on the cell line used and so included three additional ChIP-seq datasets (ENCFF699KBP; ENCFF845PAS; ENCFF723DLM) from distinct cell types. It is important to note that while some of the regions tested in the cell lines are at the same locations, there are large differences in the open chromatin regions (and thus site accessibility) across these cell types, often with >50% unique sites between cell types (bottom half of Figure 4). We saw high levels of correlation using these additional cell types, with R^2^ values over 0.97 for HepG2, A549, and T47D (p-values < 1e-32). We also saw this trend between SEMs run on different cell lines for additional transcription factors including MYC, NKFB1, and FOS, suggesting that for ubiquitous transcription factors we expect there to be no appreciable difference between SEMpl outputs. (Supplementary Figures 2-4)

**Figure 4.**
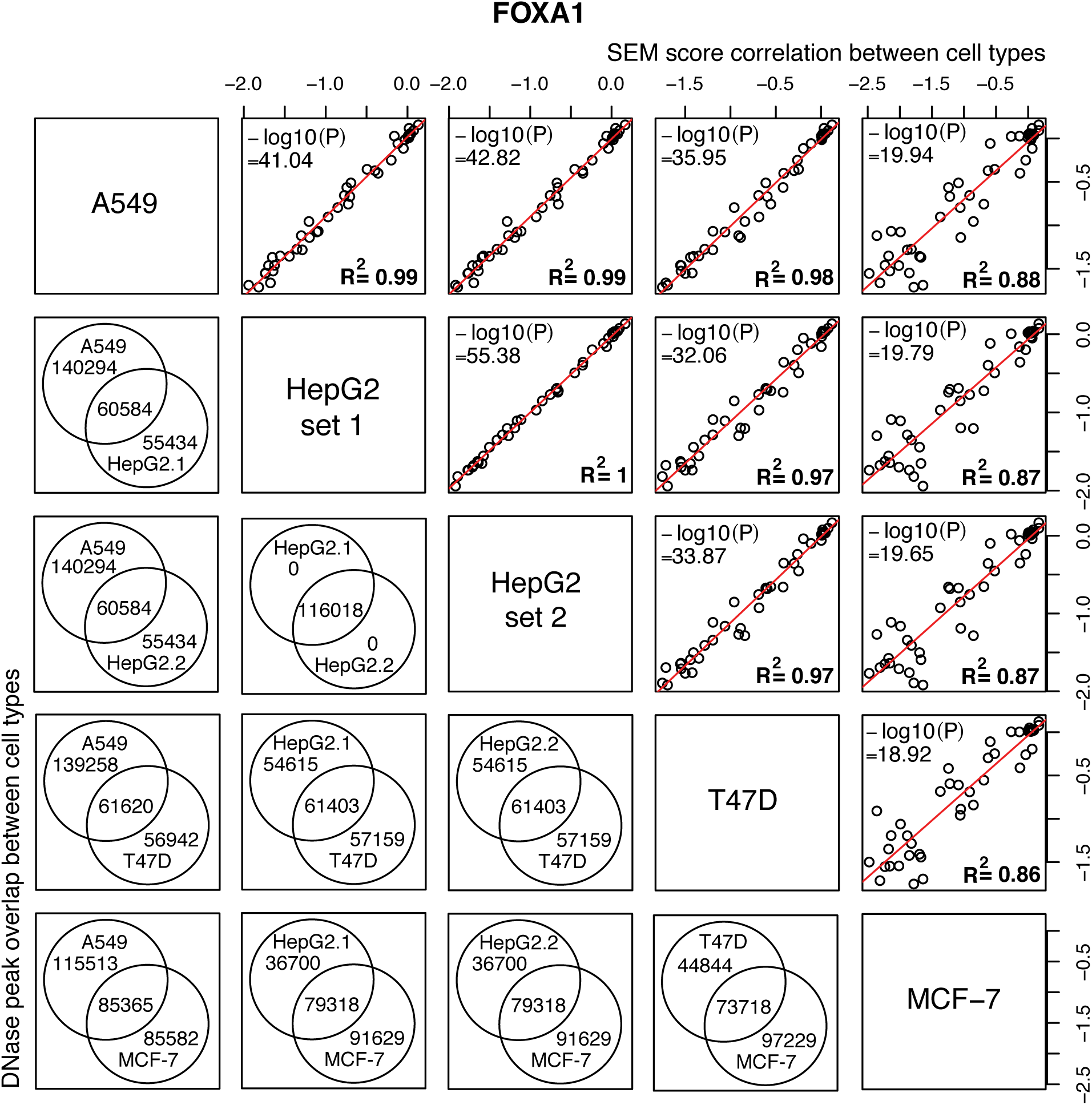
Different ChIP-seq input produce similar SEMs. The top right half of the table shows a least square regression analysis which reveals that FOXA1 SEMs are highly correlated across four cell types and two pairs of biological replicates with correlations between samples ranging from R^2^= 0.86 and R^2^=1. The bottom left half of the table shows overlapping DNase peaks between cell types. A549, lung carcinoma cell line; HepG2, hepatocellular carcinoma cell line; T47D, breast tumor cell line; MCF-7, breast adenocarcinoma cell line; HEK293, embryonic kidney cell line

It has been proposed that there may be binding affinity differences between cell types when a transcription factor has known cell type specific functions or cofactors. To address this, we investigated the proto-oncogene *MYC,* which encodes for the transcription factor c-myc known to have distinct functions and co-factors between differing cell types [36, 37]. Interestingly, we found c-myc yielded a highly similar pattern between almost all cell types observed, but a distinct SEM plot in HeLa cells that cannot be explained by low data quality (Supplementary Figure 4). This suggests that SEMpl can also be used to help identify transcription factors that have distinct cell type specific functions. However, this seems to be the exception rather than the rule as the majority of SEMs we observed were cell-type agnostic.

Finally, we asked if the starting PWM for a TF would influence the final SEM output. We found no appreciable difference in SEMpl outputs when using different starting PWMs, given that the starting PWMs represent the general binding of the transcription factor of interest (Supplementary Figure 5). However, certain PWMs and/or datasets do not contain enough information about the binding of a TF and so do not produce any significant enrichments in the final SEM output and are thus discarded (Supplementary Table 1).

**Figure 5.**
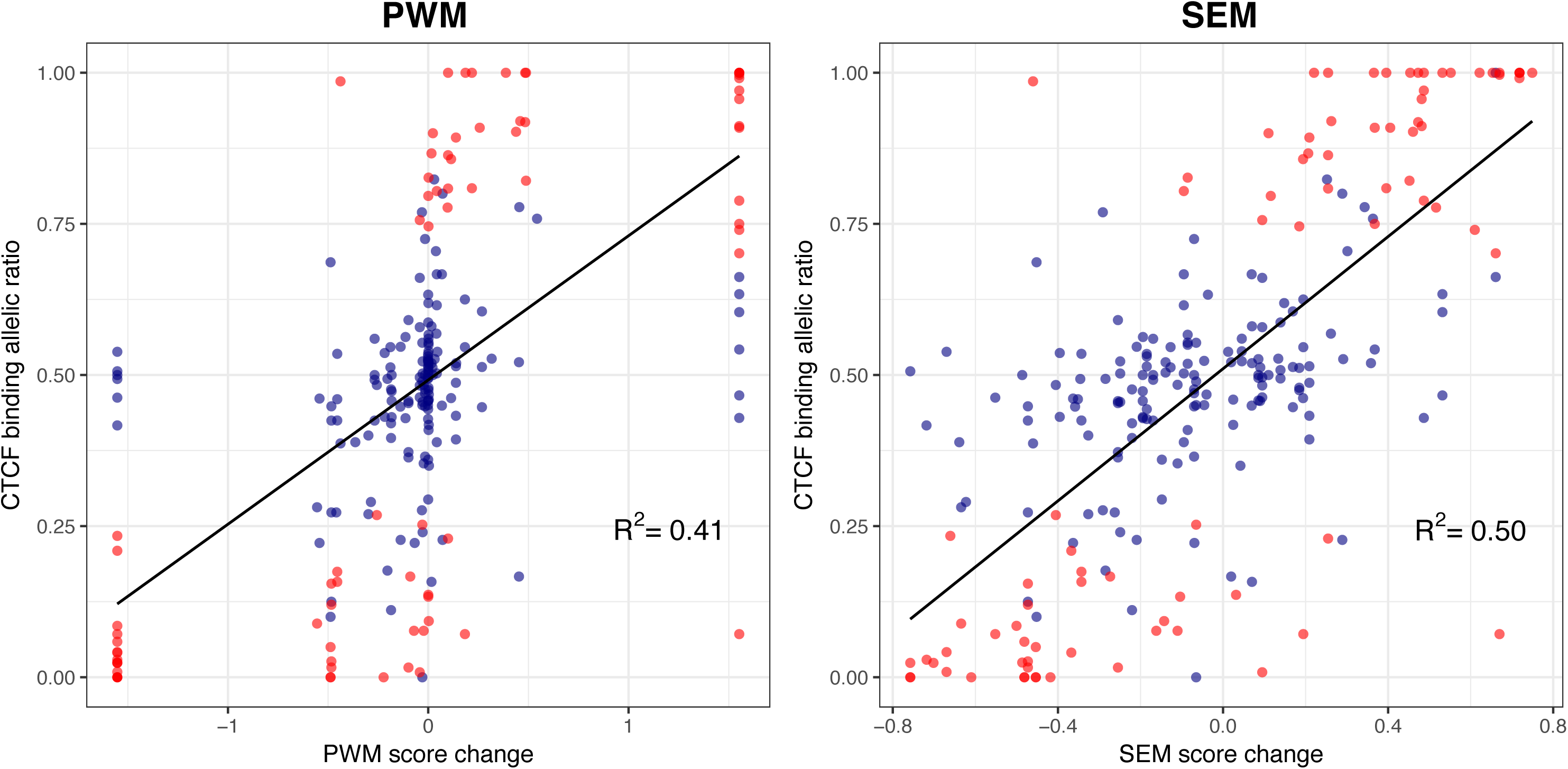
SEMs reflect allele specific CTCF binding patterns. Linear regression reveals a higher correlation between SEM score change and binding affinity change in two alleles of heterozygous sites (R^2^=0.50) than PWM scores (R^2^=0.41). Allele binding affinity change was measured by allelic ratio, which is the ratio between CTCF ChIP-seq read counts from maternal allele and total read counts from two alleles. Allele specific binding sites (red points) generally have larger changes on SEM scores.

### SEMpl recapitulates known allele specific binding patterns

Allele specific binding differences in noncoding regions of the genome have long been associated with regulatory sequence [20, 21]. To compare SEM scores against known allele specific binding data, we annotated heterozygous sites in the GM12878 cell line with ChIP-seq read counts from two alleles using ENCODE CTCF ChIP-seq datasets. Least squares regression analysis of SEM or PWM score changes against ChIP-seq signal changes of these 240 heterozygous sites in CTCF binding sites revealed a higher correlation for SEM score changes with an R^2^ of 0.50 compared to a PWM R^2^ of 0.41 (Figure 5). We also observed a more dispersed distribution of SEM score changes, where the allele specific binding sites have overall larger changes between two alleles (red points in Figure 5). These indicate that the SEM score is more able to capture the change of TF binding affinity compared to PWM.

To validate that SEMpl scores accurately predict transcription factor binding affinity changes *in vitro* we compared SEMpl scores to previously generated ChIP-qPCR data, which measures endogenous transcription factor binding affinity [31]. ChIP-qPCR was generated from 10 allele specific FOXA1 binding sites in the genome. Regression analysis comparing SEMpl scores to changes in transcription factor binding by ChIP qPCR analysis reveal that SEM scores are a better predictor of SNP changes (R^2^= 0.8) than PWMs (R^2^= 0.44) (Supplementary Figure 6). Additional studies of endogenous transcription factor binding in sites not associated with a common SNP will be necessary to demonstrate the full power of SEMpl. These results suggest that SEMpl has the ability to return biologically meaningful results, and can be used to predict the direction and magnitude of allele specific changes.

### SEMpl predictions agree with experimentally validated SNPs from the literature

To verify that SEMpl would allow researchers to identify variants potentially leading to transcription factor binding changes associated with gene expression changes, we validated our method against four published TFBS SNPs found to disrupt transcription factor binding (Supplementary Table 2). In most cases, we found SEMpl predictions agreed with the direction of the validated changes, as well as the magnitude, when available. For example, a T to G change in position 12 of a TCF7L2 binding site was found to increase binding affinity by 1.3 fold by mass spec, where SEMpl predicted a 1.27 fold increase [14]. Only one of the four SEMpl predictions that we identified did not match the experimentally determined variant. This C/T allele in position11 of a FOXA2 binding site was predicted to decrease binding affinity by FAIRE-seq, however SEMpl predicted no difference in binding between the two alleles. Interestingly, PWMs also predicted no difference in binding between the two alleles, suggesting additional factors may be at play.

We also compared SEMpl predictions to predicted variant effects measured through a massively parallel reporter assay (MPRA) [38]. We found a correlation between these previously published expression changes and SEM score changes (Supplementary Figure 7). However, this relationship was not as strong (R^2^= 0.23), though still outperforming PWMs (R^2^= 0.16), possibly due to the non-linear relationship between transcription factor binding, regulatory element use, and gene expression.

## Discussion and Conclusions

A deeper understanding of the role noncoding variants play in altering gene expression is critical to fully illustrate the regulatory complexity of our genome and is an important first step towards developing tools for personalized medicine. Approaches such as the IGR method have expanded our ability to use currently available data to predict SNPs that play a regulatory role and have successfully been implemented in multiple studies to link human disease to specific transcription factors and their binding sites. Since its release the IGR method has been used to successfully identify functional SNPs in TFBSs from GWAS data for breast cancer, atrial fibrillation, and lupus [39-41]. Functional predictions for these SNPs were experimentally validated, suggesting that the IGR process can be a robust method for functional noncoding GWAS SNP prediction. Unfortunately, this method is not accessible for widespread use. By developing a tool which generalizes the IGR methodology to predict the magnitude and direction of effect of all SNPs within a TFBS, we can identify novel variants associated with human disease in TFBSs genome-wide.

In this paper we introduced SEMpl, a new tool designed to identify putative deleterious mutations in TFBSs. SEMpl predictions reflect known patterns of transcription factor binding while providing additional information about magnitude and direction of predicted change. We demonstrate that SEMpl provides more robust and consistent predictions both on a single variant and a TFBS kmer level than the current standard, PWMs. The method leverages simulation and real data to better model strength of binding rather than a consensus sequence. Additionally, SEMpl scores correlate with known ASB sites and agree with previously published variants known to alter transcription factor binding affinity.

SEMpl was designed to be easy to use and accessible. In addition to being available as an open source application, precompiled SEM plots for 89 transcription factors from over 200 PWMs, are available online. While SEMpl is currently limited to transcription factors with available ChIP-seq and PWM data we may be able to eliminate the use of PWMs to guide TFBS loci in future versions of our pipeline, reducing bias and expanding our list of compatible transcription factors. In addition, we are working to include additional genomic features, such as DNA methylation which would allow the inclusion of additional bases to SEM plots and a more nuanced understanding of transcription factor binding.

SEMpl’s ability to better predict the impact of genomic variation on transcription factor binding has broad implications to the cross disciplinary study of the regulatory genome. SEMpl has great usability for prioritizing GWAS SNPs for experimental follow-up. With the increased need for experimental validations following large-scale genomics studies, we anticipate that annotation tools, such as SEMpl, will be critical in revealing developmental and disease-associated regulatory SNPs.

## Methods

### Usage/Accessibility

SEMpl is open access and can be downloaded from github: https://github.com/Boyle-Lab/SEM_CPP. Over 200 pre-computed SEMpl scores can be found in Supplemental Material.

### SEM pipeline methodology

SEMpl starts reads in the user-given PWM which is expanded into all valid kmers greater than a weak p-value cutoff of 4^−5^, established by TFM-score, to create a list of endogenous kmers. This enumerated kmer list is used to generate 4 × kmer length additional lists in which all possible SNPs are systematically created *in silico*. In addition, the kmers in the endogenous kmer list are randomly scrambled to create a scrambled baseline where transcription factor binding would not be expected. These kmer lists (endogenous, SNP, scrambled) are all aligned to the human autosomes (hg19) using bowtie.[42] Matching ChIP-seq loci from bigwig files downloaded from the ENCODE database are matched to the aligned kmer loci and the greatest ChIP-seq value within a 50bp window are used to define the maximum ChIP-seq value. Values not within a DNase-peak from narrowPeak files downloaded from the ENCODE database are filtered out. Resulting aligned loci and ChIP-seq values stored in a cache, which allows for a quick lookup of non-unique kmers without realignment. Average ChIP-seq values are calculated across each kmer list to create a matrix. Additionally, the enumerated baseline is calculated from the average of the enumerated kmer list, and the scrambled baseline from the average of the scrambled baseline kmer list. These scores are then log2 transformed and normalized to set the enumerated baseline to 0. This normalized matrix represents an SEM, which is then plotted via R.

The resulting matrix from the first SEM iteration is used to generate a PWM-style matrix and is used as the starting PWM for the second iteration. All iterations after the first use a slightly more stringent p-value cutoff of 4^−5.5^ to generate kmers. This process continues until the number of kmers from the endogenous kmer list does not change or until 250 iterations

SEMpl output files include error messages during the run (.err), the cache, a tally of kmer similarity between iterations (kmer_similarity.out), and an output file containing information on run time and where the program is in the run (.out). Additionally, within each iteration, output files include the alignments for the SNP kmer lists (alignment folder) and endogenous and scrambled kmer lists (baseline folder) which include the aligned loci and ChIP-seq signal. A quality control file is also provided within each iteration file that provides the number of kmers mapped within the iteration, as well as a –log10(p-value) representing the average of 100 T-tests value from 1000 randomly chosen kmers from the SNP signal files versus 1000 randomly chosen kmers from the scrambled signal file. We used a threshold of 2.5 to report confidence in a SEM run.

SEMpl options include –readcache, which can be used to speed up a run for which a cache has already been created. SEMpl is written in C++ and R. PWMs were created using the R package seqLogo [43].

### Correlation with ChIP-seq data

All possible kmers from the original FoxA1 PWM were generated. For each unique kmer, average ChIP-seq signal and standard error was calculated. PWM and SEM scores were calculated for each kmer. Correlations cutoffs were calculated for PWMs above the standard TFM p-value cutoff (p-value = 4^−8^) typically used for PWM visualization. Correlation cutoffs for SEMs were defined as the average scrambled baseline across all iterations for a single transcription factor run.

### SEM correlation across runs

SEM outputs from different staring ChIP-seq or PWM data were compared using least square regression in R. More details about the datasets used for analysis can be found in Supplemental Table 1. Overlapping DNase-seq peaks were downloaded from ENCODE and calculated using bedtools [44]. SEMpl runs from the same cell type, and therefore using the same DNase dataset, show no difference in DNase peak overlap.

### Allele specific CTCF binding pattern analysis

Allele specific binding sites were defined as loci containing one or more heterozygous SNPs while showing significant differences in ChIP-seq signal from two alleles. We applied the AlleleDB pipeline to count the number of ChIP-seq reads from two alleles respectively for each heterozygous site and identified 468 allele specific binding sites at an FDR of 5% [45]. CTCF ChIP-seq data from GM12878 cell line was used in this analysis (Accession number: ENCSR000DZN). For all heterozygous sites within CTCF ChIP-seq peaks in GM12878 cell line, 240 of them also have matching CTCF PWMs, which we further used for the comparison of SEM and PWM scores. For those 240 heterozygous sites, we calculated the allelic ratio defined by the ratio between the number of ChIP-seq reads from the maternal allele and the total number of reads from two alleles. We then evaluated the correlation between the change of SEM or PWM scores and allelic ratios.

## Supporting information

All SEMs

## Acknowledgements

This project was supported by NIH U41 HG009293 to APB and through the University of Michigan Genome Science Training Program T32 HG00040 to SSN. CM and CW were supported by the University of Michigan Undergraduate Research Opportunity Program. We also thank Jessica Switzenberg and Courtney Asman for their feedback and support of this project.

