## Supplemental Information for "Predicting the effects of SNPs on transcription factor binding affinity"

**Supplementry Figure 1.** Correlation of PWM scores for full kmers versus average ChIP-seq signal.

**Supplementry Figure 2.** FOXA1 shows robust correlations across different ChIP-seq input data.

**Supplementry Figure 3**. NFKb shows robust correlations across different ChIP-seq input data.

**Supplementry Figure 4.** FOS shows robust correlations across different ChIP-seq input data.

**Supplementry Figure 5.** Different starting PWMs yield highly similar SEMs.

**Supplementry Figure 6.** SEMs scores are a better predictor of transcription factor binding changes from SNPs than PWMs when compared to ChIP qPCR analysis for FoxA1.

**Supplementary Figure 7.** SEM scores are correlated with reporter expression changes.

**Supplementry Table 2.** Known variants affecting transcription factor binding affinity.

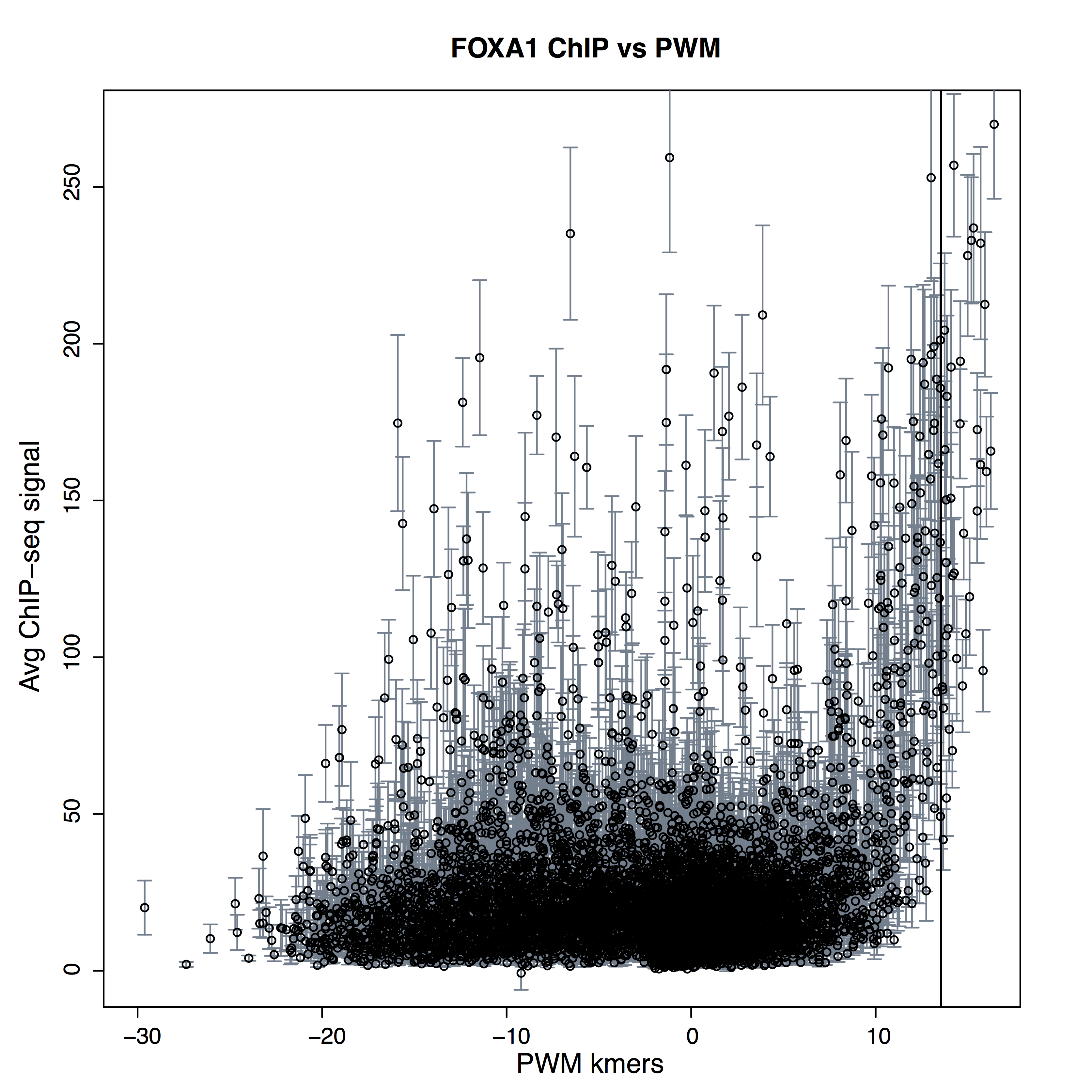

Supplementry Figure 1. Correlation of PWM scores for full kmers versus average ChIP-seq signal. All possible kmers from a starting FOXA1 PWM were included. This is an extension of Figure 3, with x-axis bounds from -30 to 15.

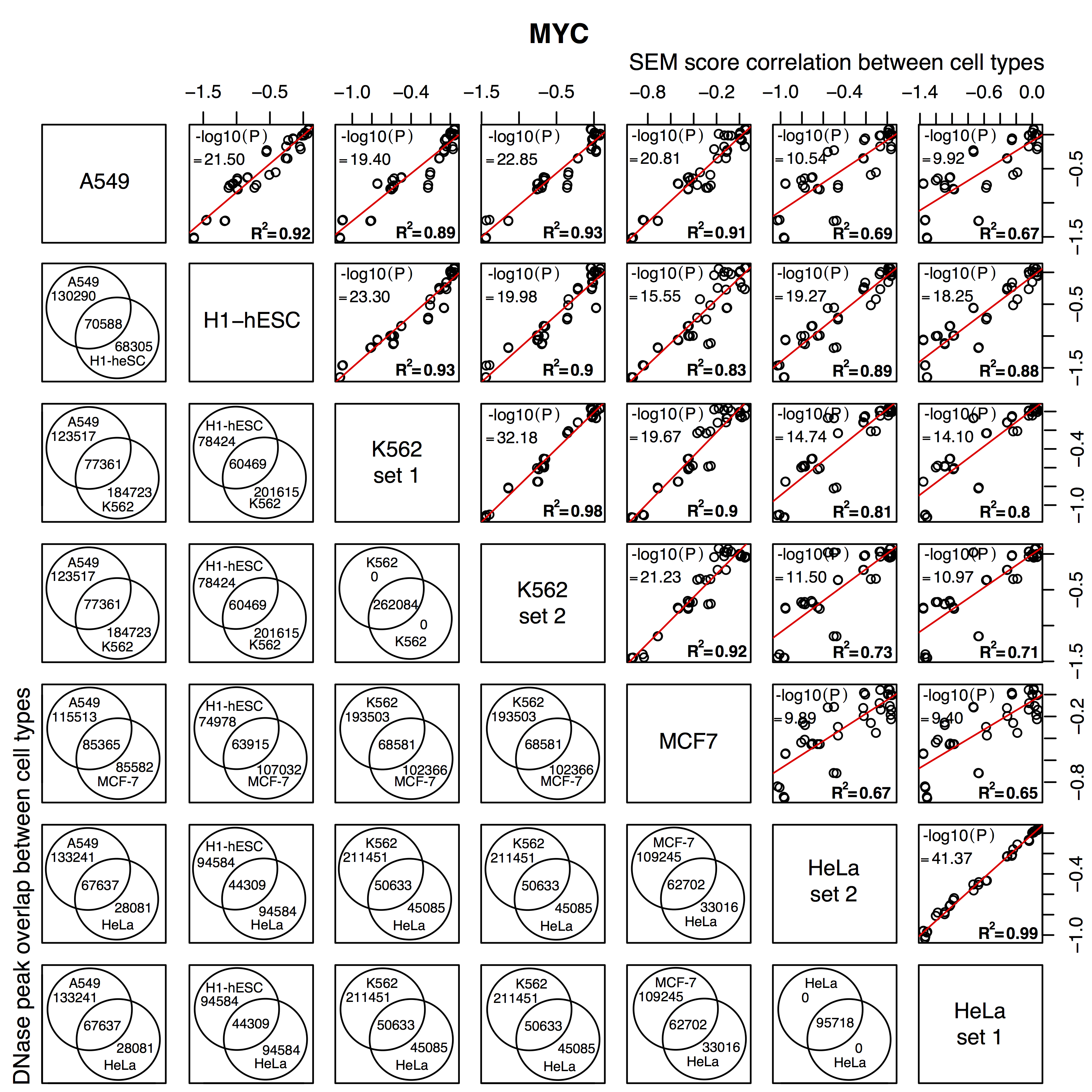

Supplementry Figure 2. MYC shows cell-line specific binding affinity across different ChIP-seq input data (HeLa v. others). The top right half of the table shows a least square regression analysis, while r-squared analysis and p-values can be found on the bottom left.

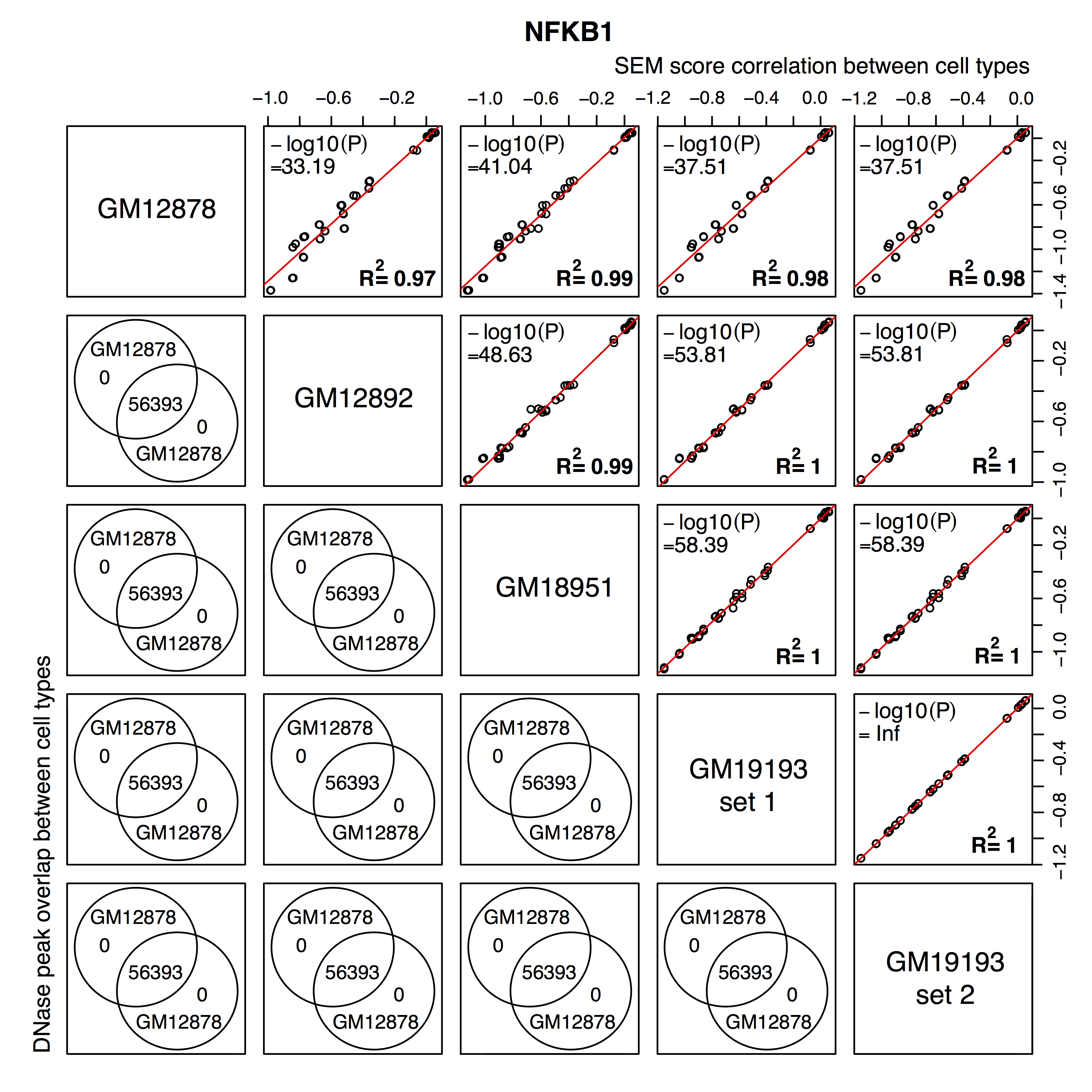

Supplementry Figure 3. NFKb shows robust correlations across different ChIP-seq input data. The top right half of the table shows a least square regression analysis, while r-squared analysis and p-values can be found on the bottom left.

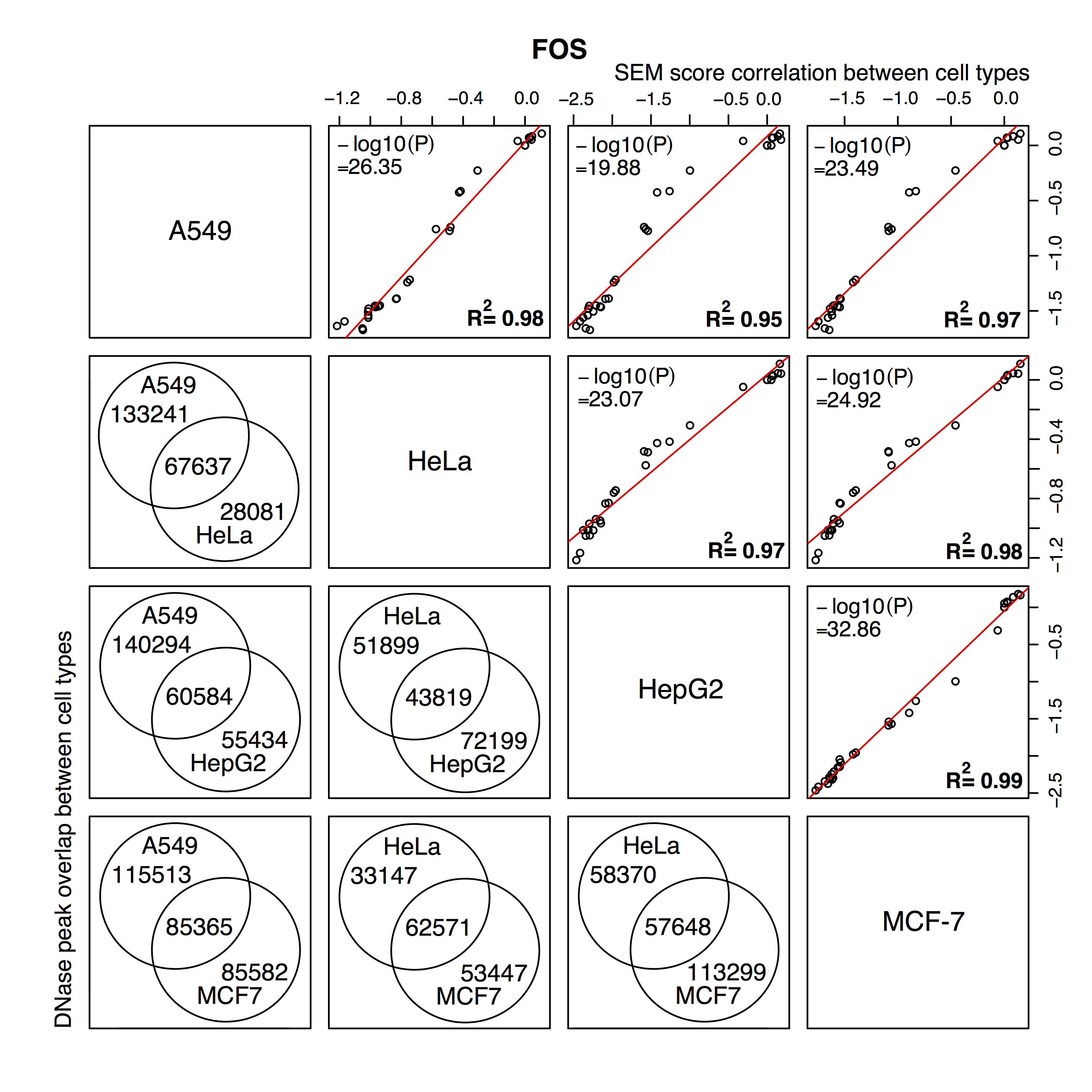

Supplementry Figure 4. FOS shows robust correlations across different ChIP-seq input data. The top right half of the table shows a least square regression analysis, while r-squared analysis and p-values can be found on the bottom left.

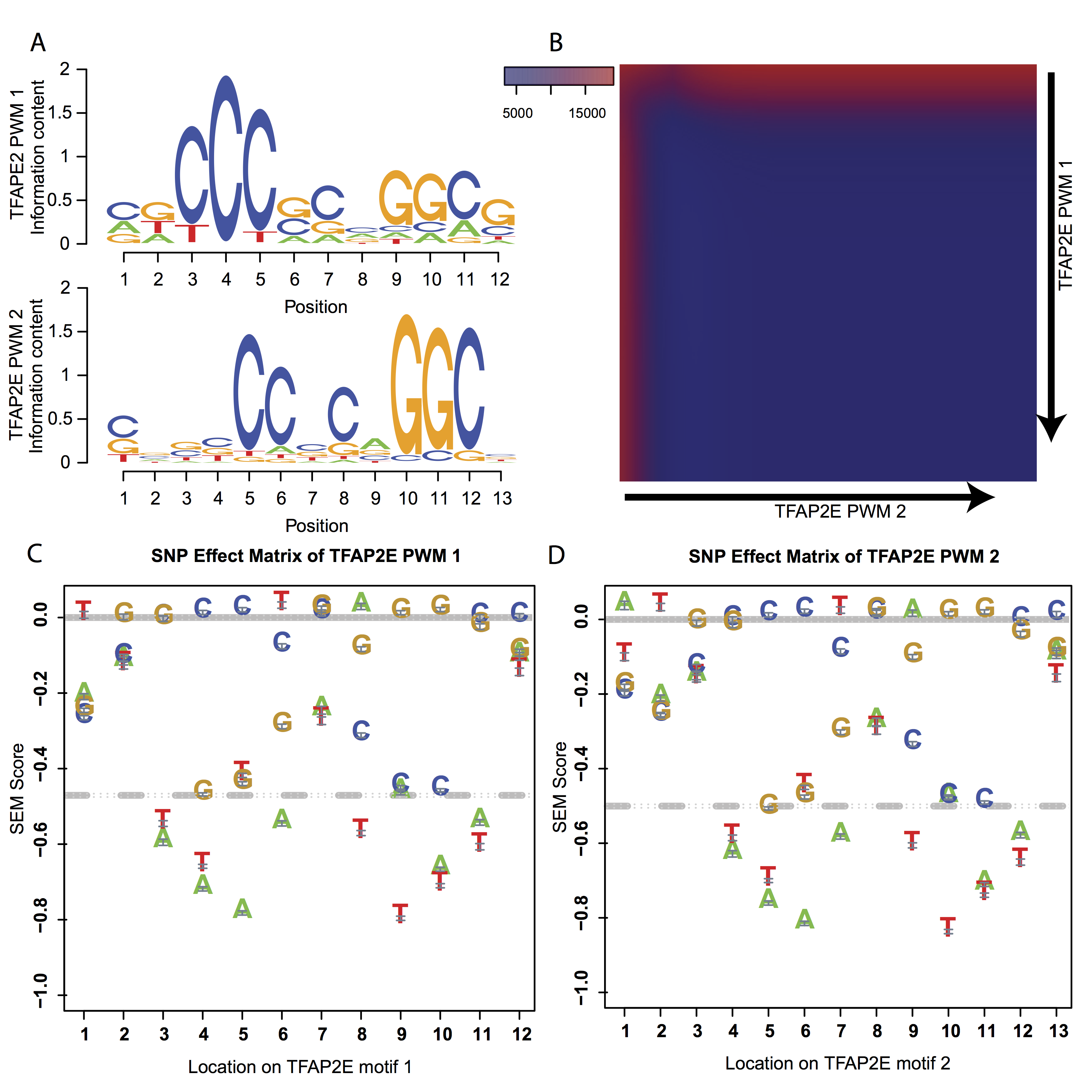

Supplementry Figure 5. Different starting PWMs yield highly similar SEMs. A. Two starting PWMs were chosen, representing distinct binding profiles and motif length (TFAP2E PWM 1, M00189; TFAP2E PWM 2, M00915). B. Both PWMs were run using SEMpl, and similarities between kmers generated from PWMs were tracked over 250 iterations. Red represents more differences between kmer sets and deep blue represents fewer. C&D. Final SEMs from starting with PWM 1 or PWM 2 appear highly similar, with the SEM of PWM 2 contianing an extra nucleotide in position 1 of the motif.

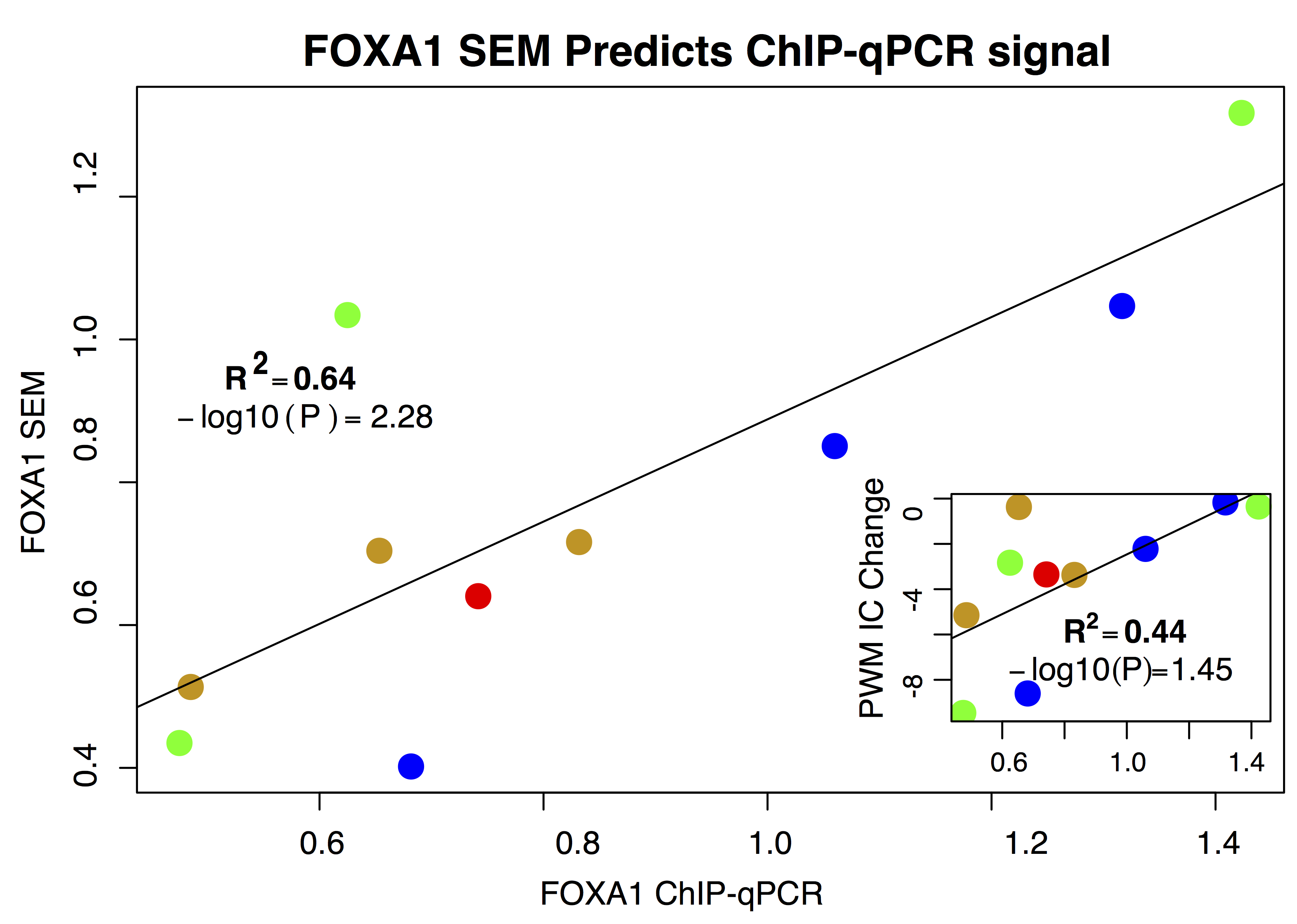

Supplementry Figure 6. SEMs scores are a better predictor of transcription factor binding changes from SNPs than PWMs when compared to ChIP qPCR analysis for FoxA1 [1]. ChIP-qPCR data for 10 FOXA1 SNPs from the IGR paper was used as an endogenous measure of binding affinity change. Colors represent the SNP change (green, A; blue, C; yellow, G; red, T). The SEM was generated using HepG2 cell line data for the ChIP-seq and DNase. PWM change is measured in change to information content (IC).

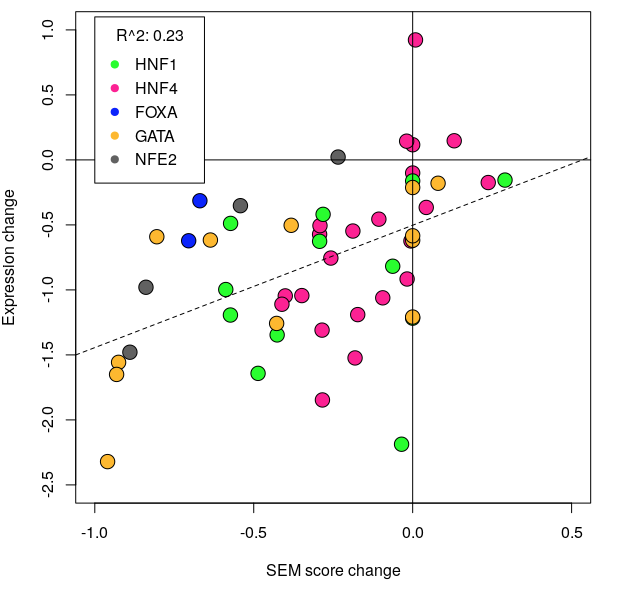

Supplementary Figure 7. SEM scores are correlated with reporter expression changes. Reporter assay expression changes across 5 transcription factors from Kheradpour et al. were compared to SEM score changes [2].

| SNP | TF | Sequence | Reference/ Variant allele | SEMpl prediction | Published validation | Reference |
| --- | --- | --- | --- | --- | --- | --- |
| rs4784227 | FoxA1 | TGTTTGC[C]GAT | C/T | 0.247x inc | inc | Cowper-Sal Iari, 2012 [1] |
| rs6983267 | TCF7L2 | AGTG[A]CTTTCATCTGCT | A/C | 0.297x inc | inc | Pomerantz, 2009 [3] |
| rs7903146 | FoxA2 | TCGATA[C]TATATA | C/T | 0.037x dec | inc | Gaulton, 2010 [4] |
| rs11257655 | FoxA2 | GGGCAAGTGT[C]TACTGGGCAT | C/T | 0.585x inc | inc | Fogarty, 2014 [5] |

Supplementry Table 2. Known variants affecting transcription factor binding affinity.
