## Supplementary figures and images for "Predicting the effects of SNPs on transcription factor binding affinity"

### BHLHB2_GM12878

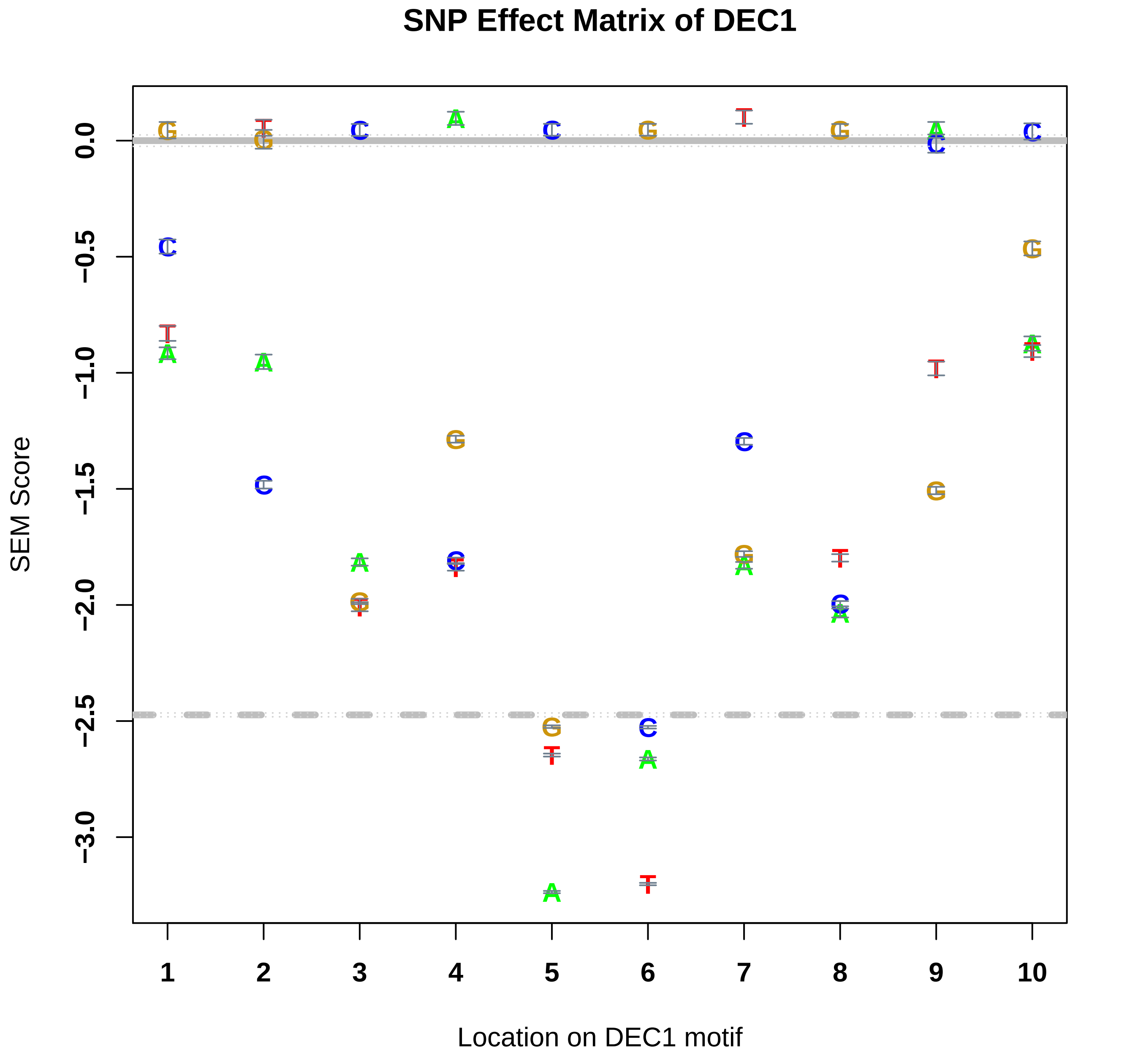

### BHLHB2_HEK293

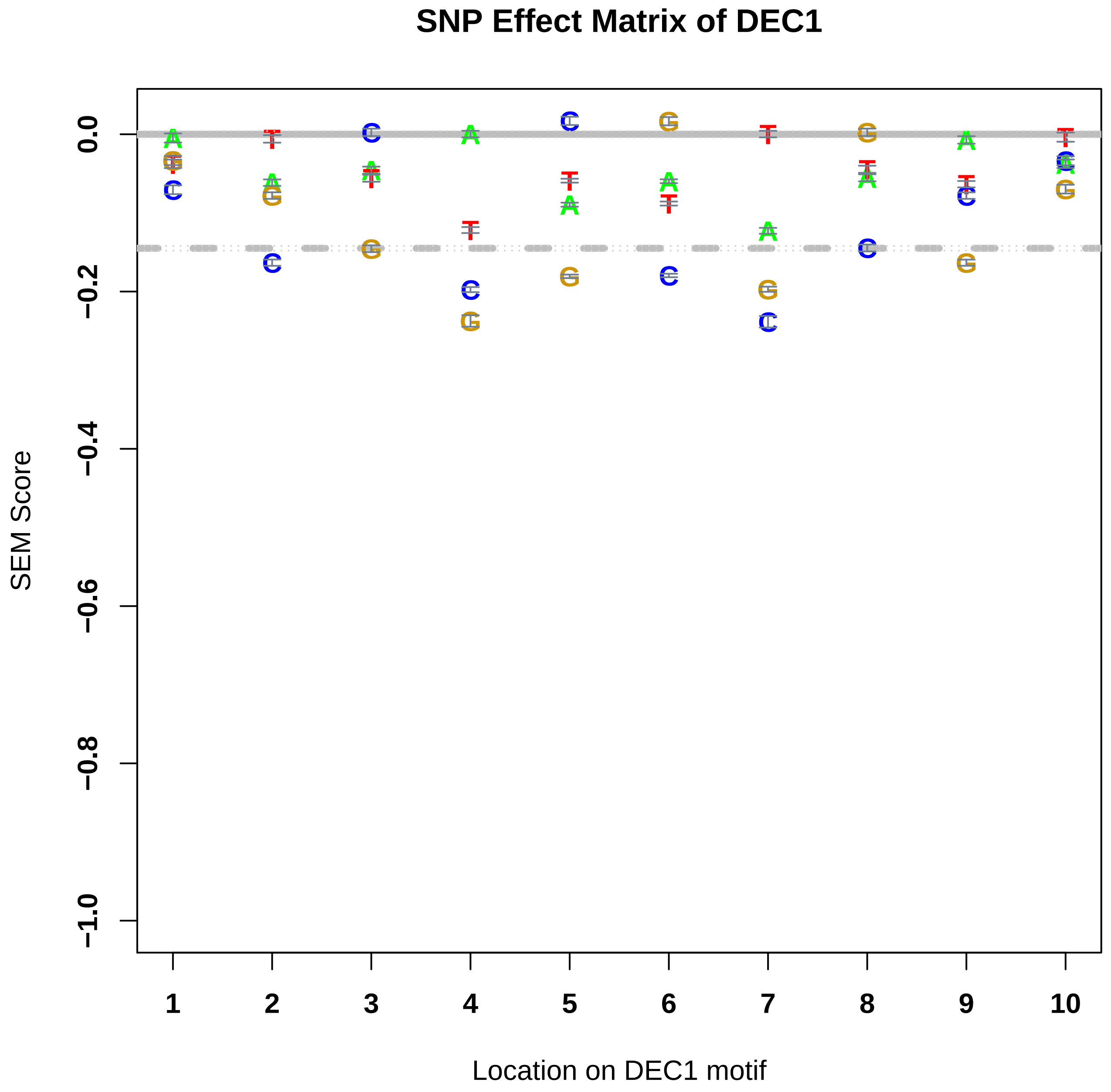

### BHLHB2_HepG2

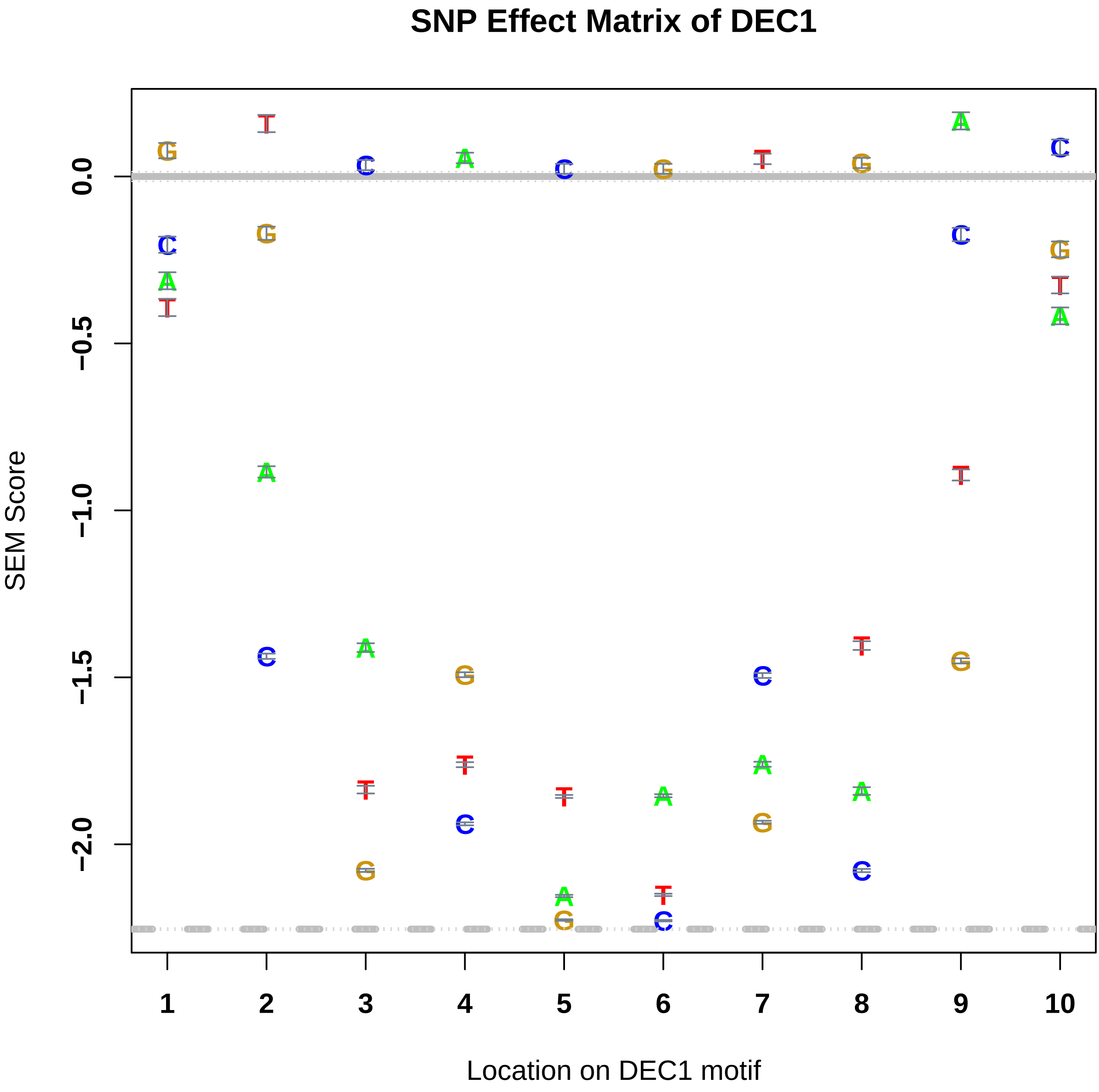

### BHLHB2_K562

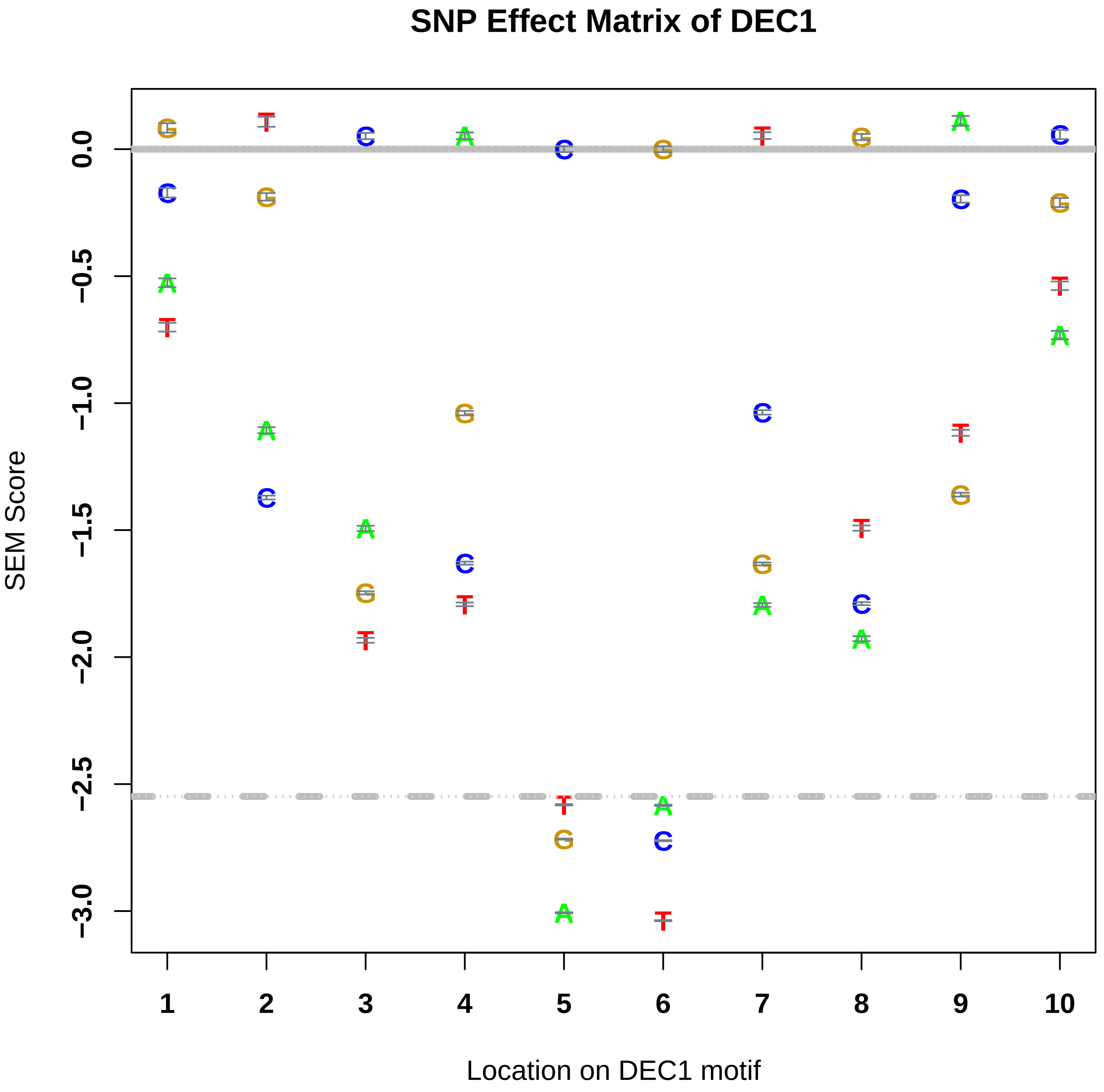

### CREB3L1_2

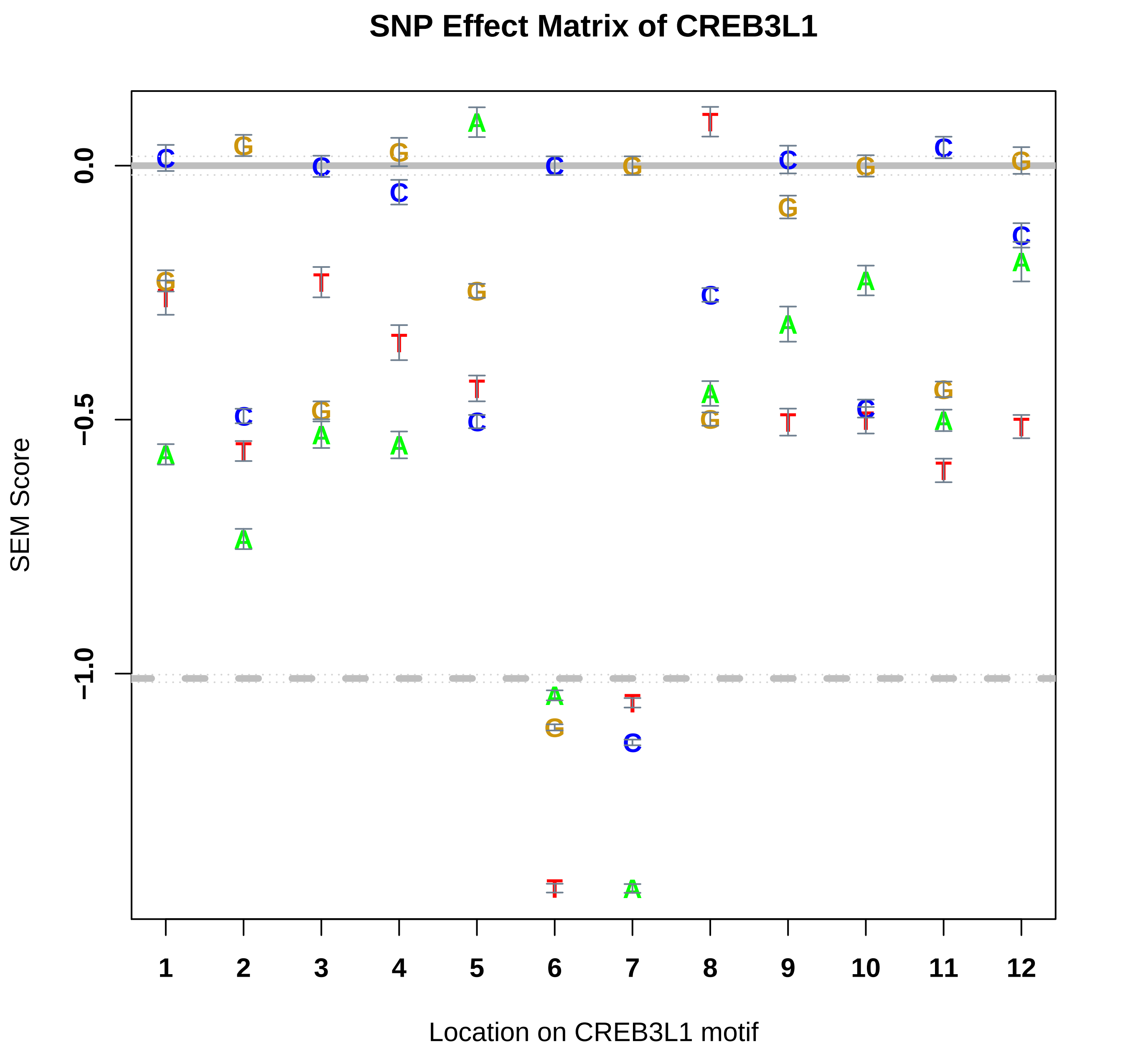

### CREB3L1_3

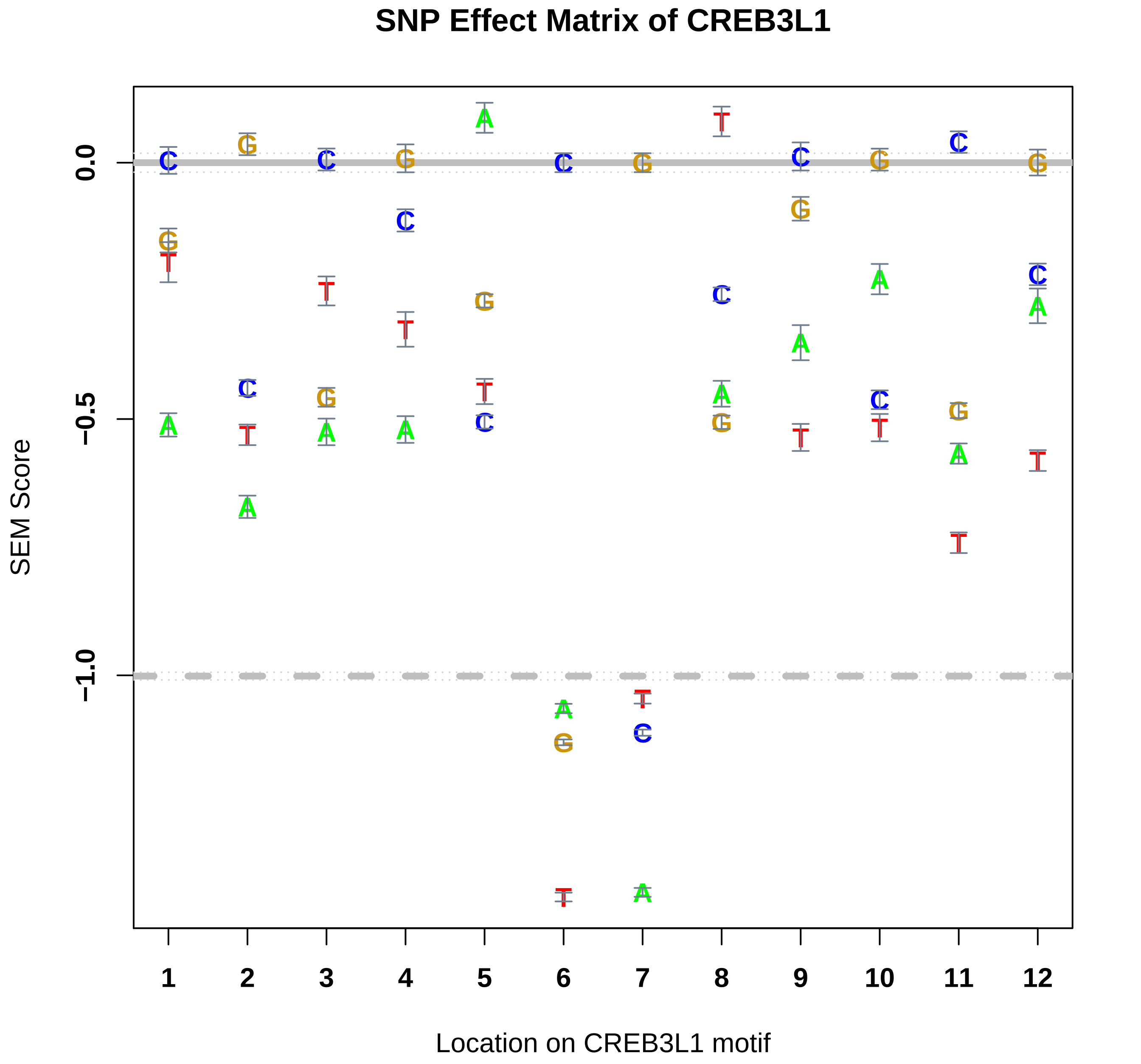

### EGR1_1

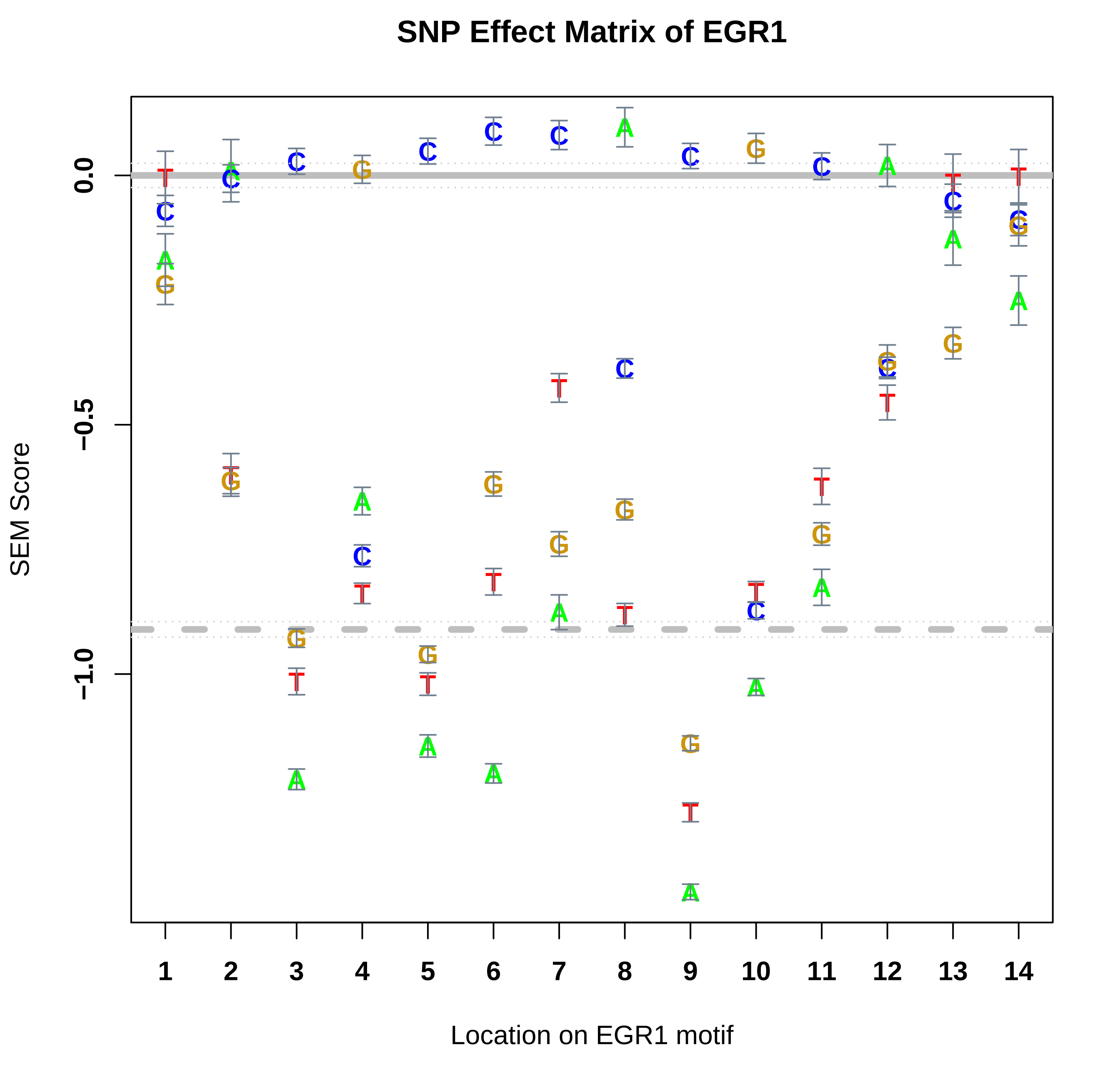

### ELF1_2

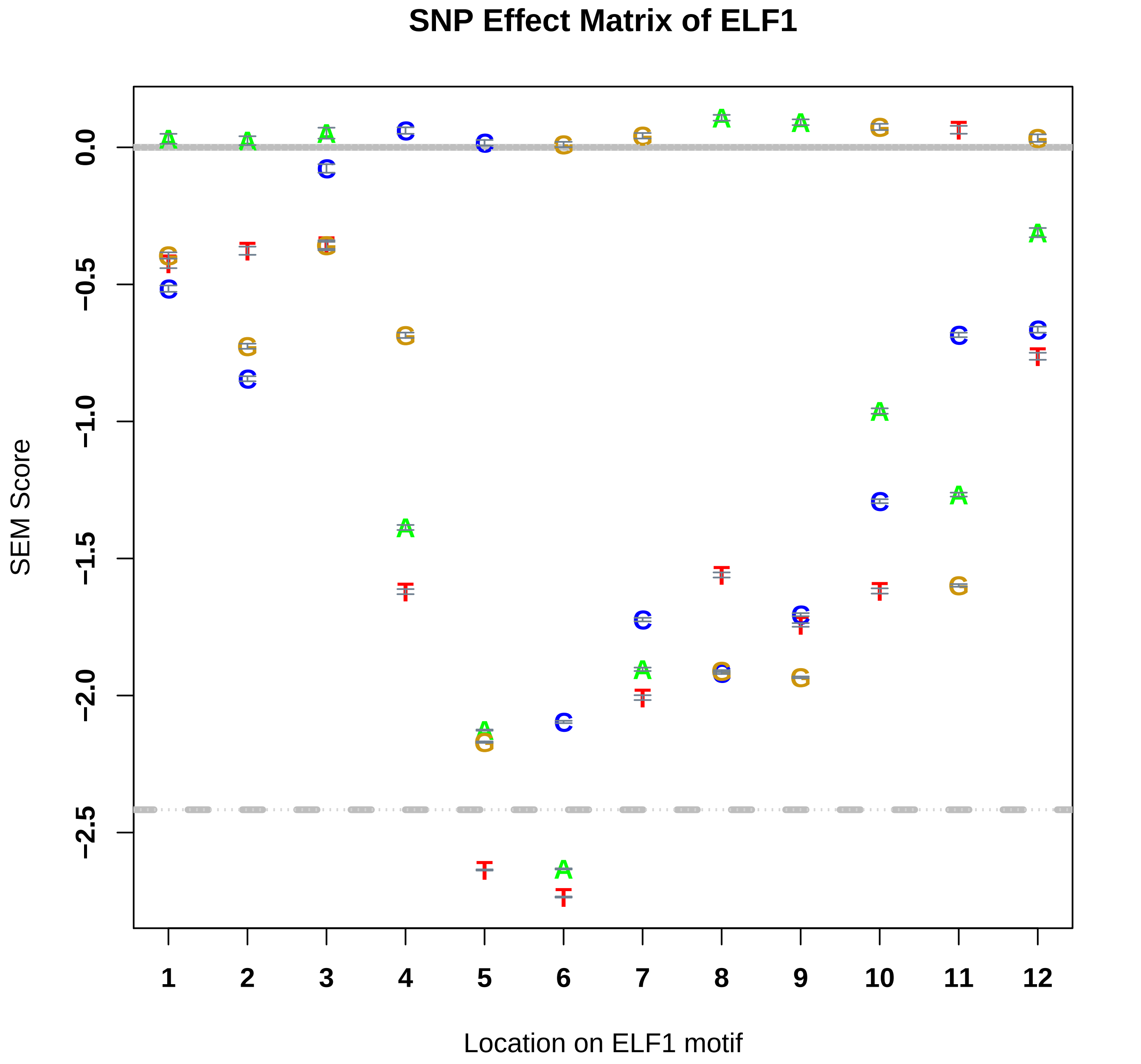

### Elf3_primary

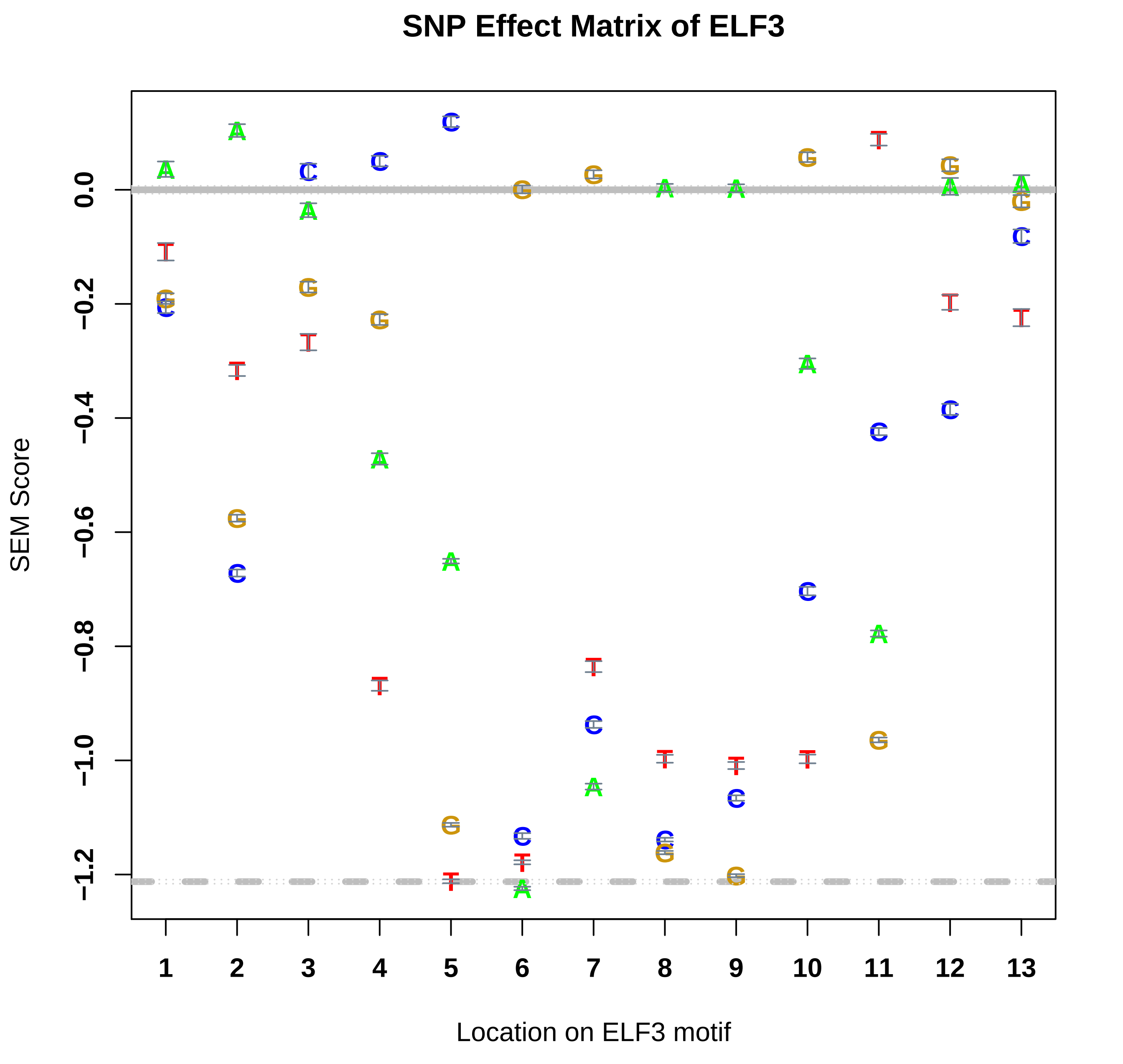

### ELF4_1.2

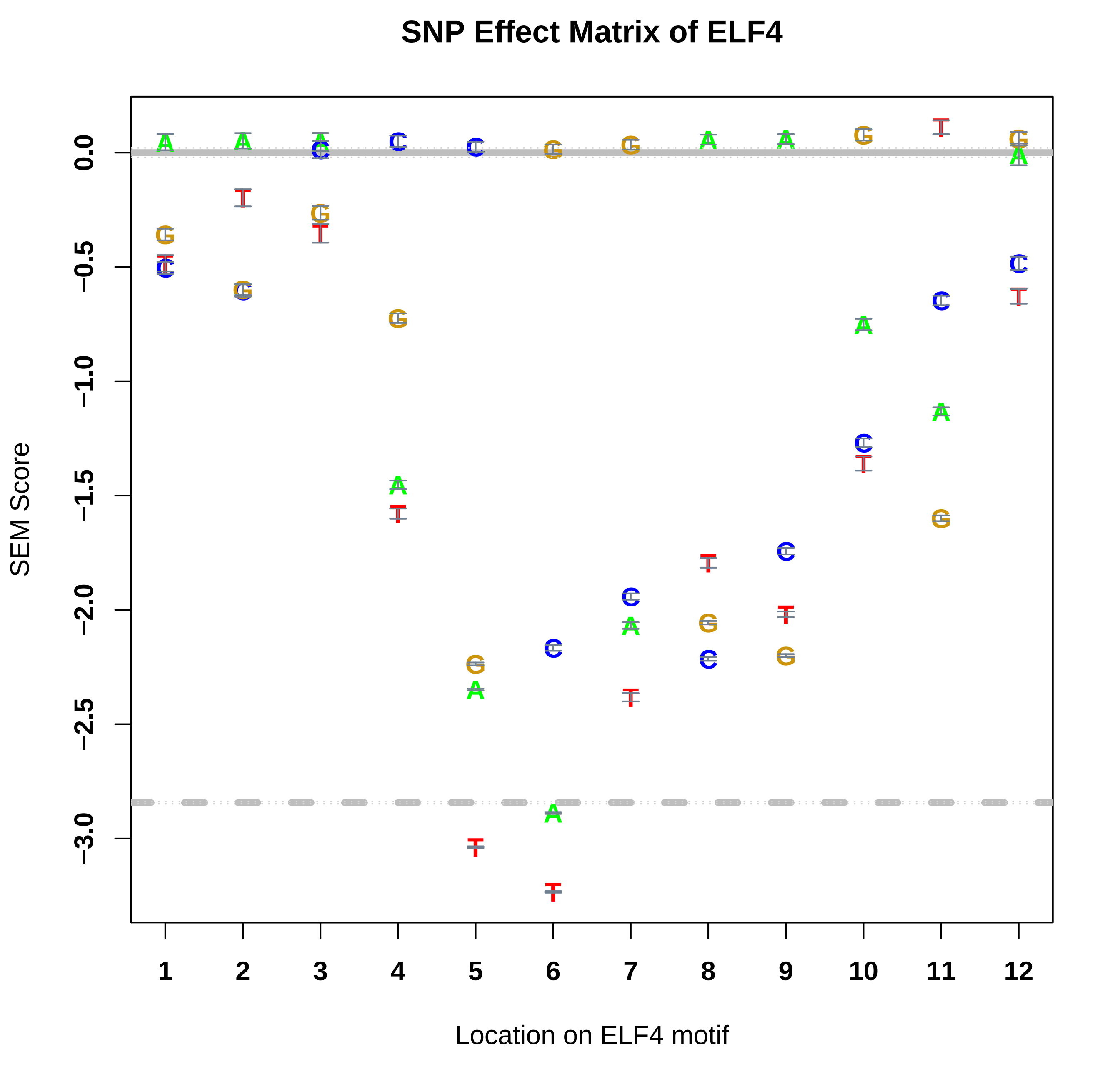

### ESR1_T47D

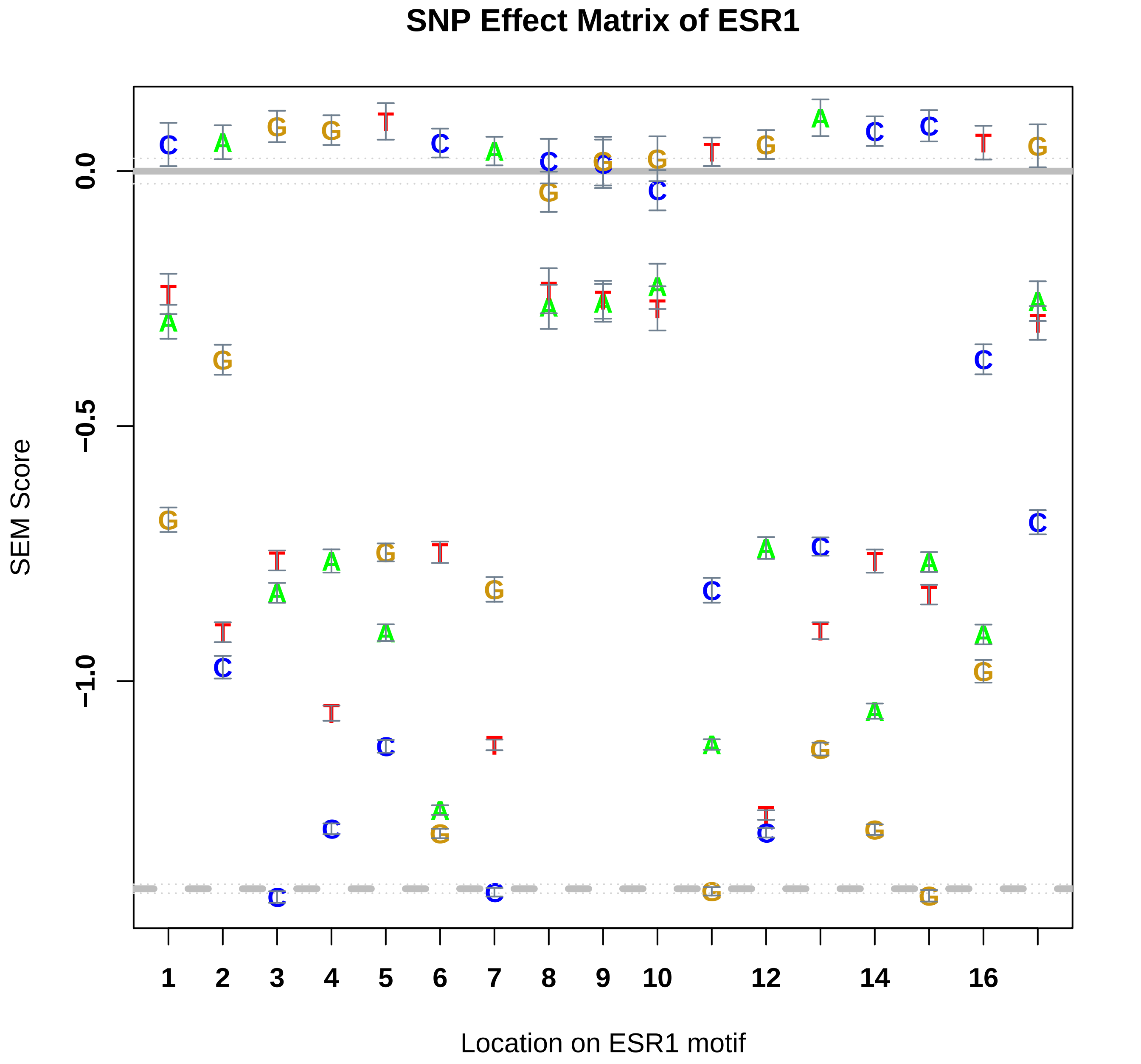

### ETV6_1_GM12878

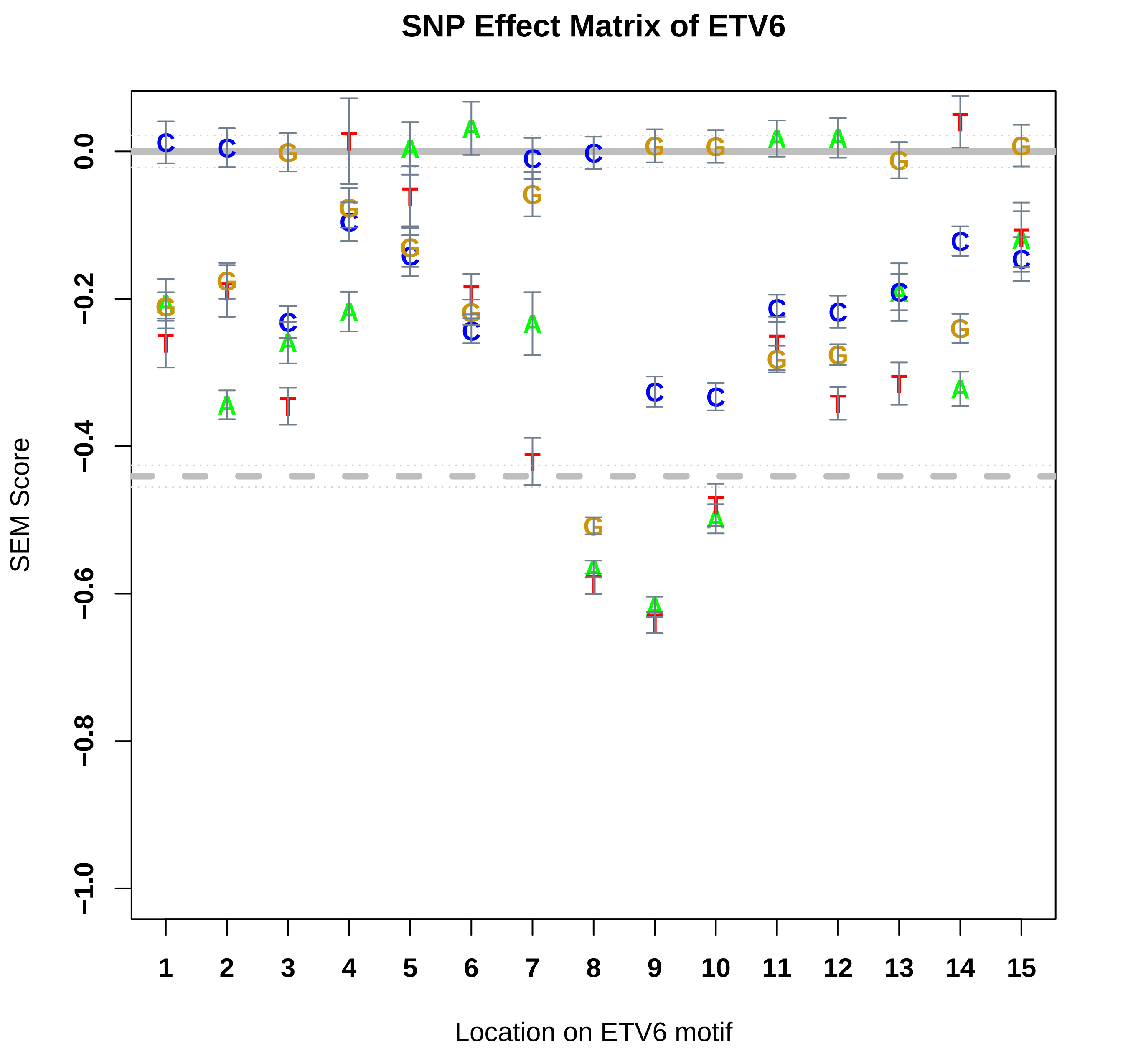

### ETV6_1_K562

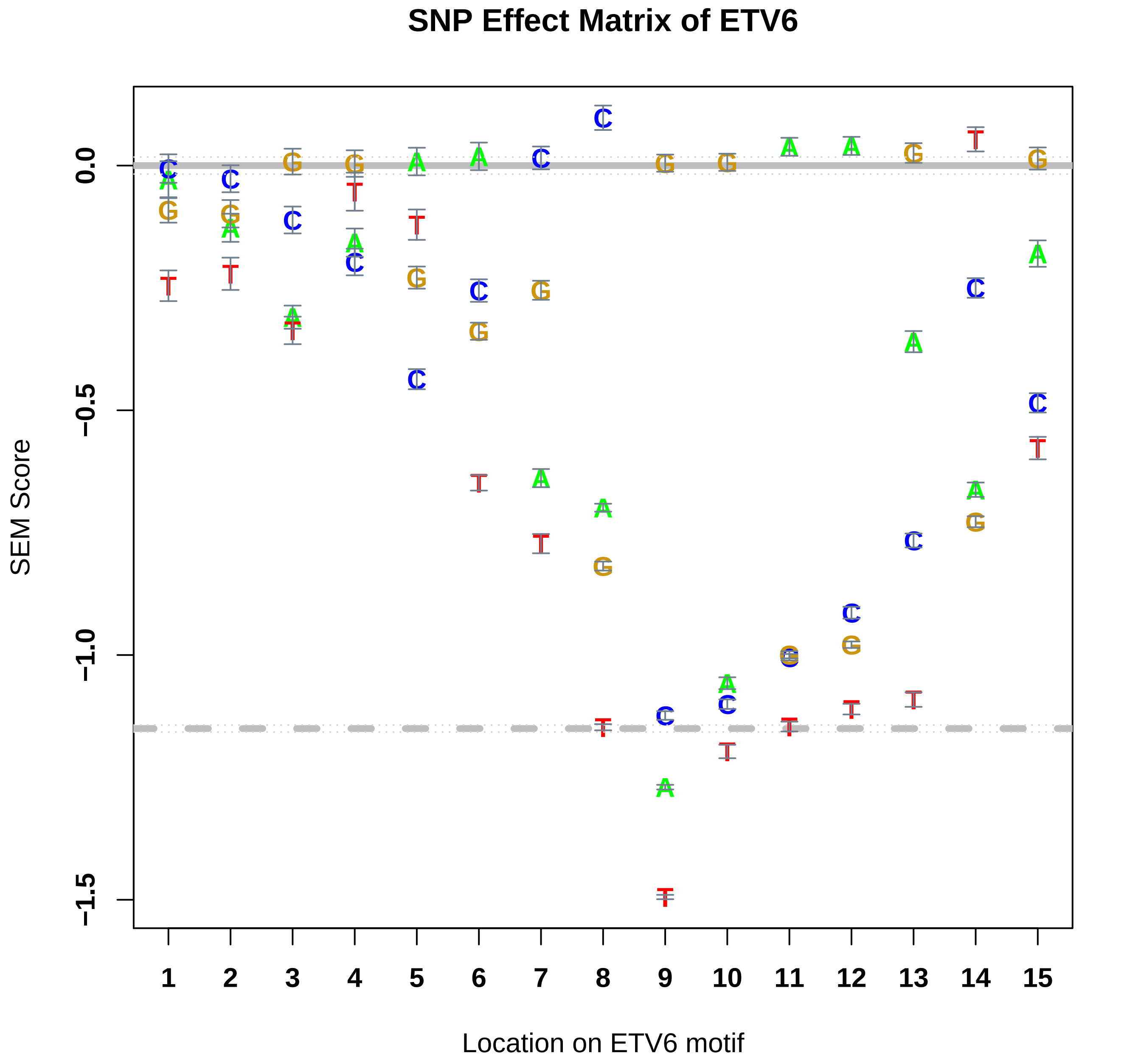

### FOXJ2_3

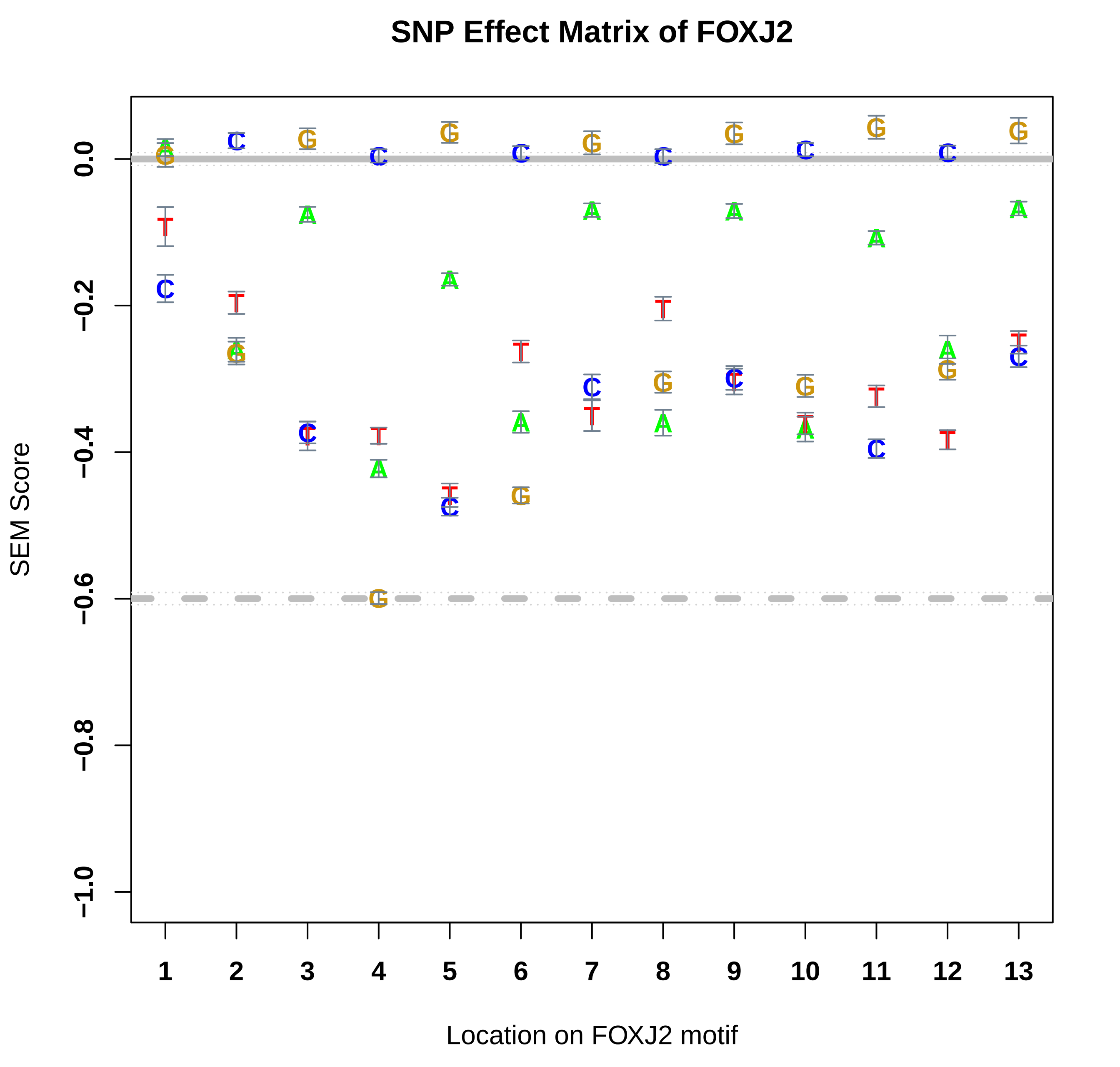

### Gabpa_secondary

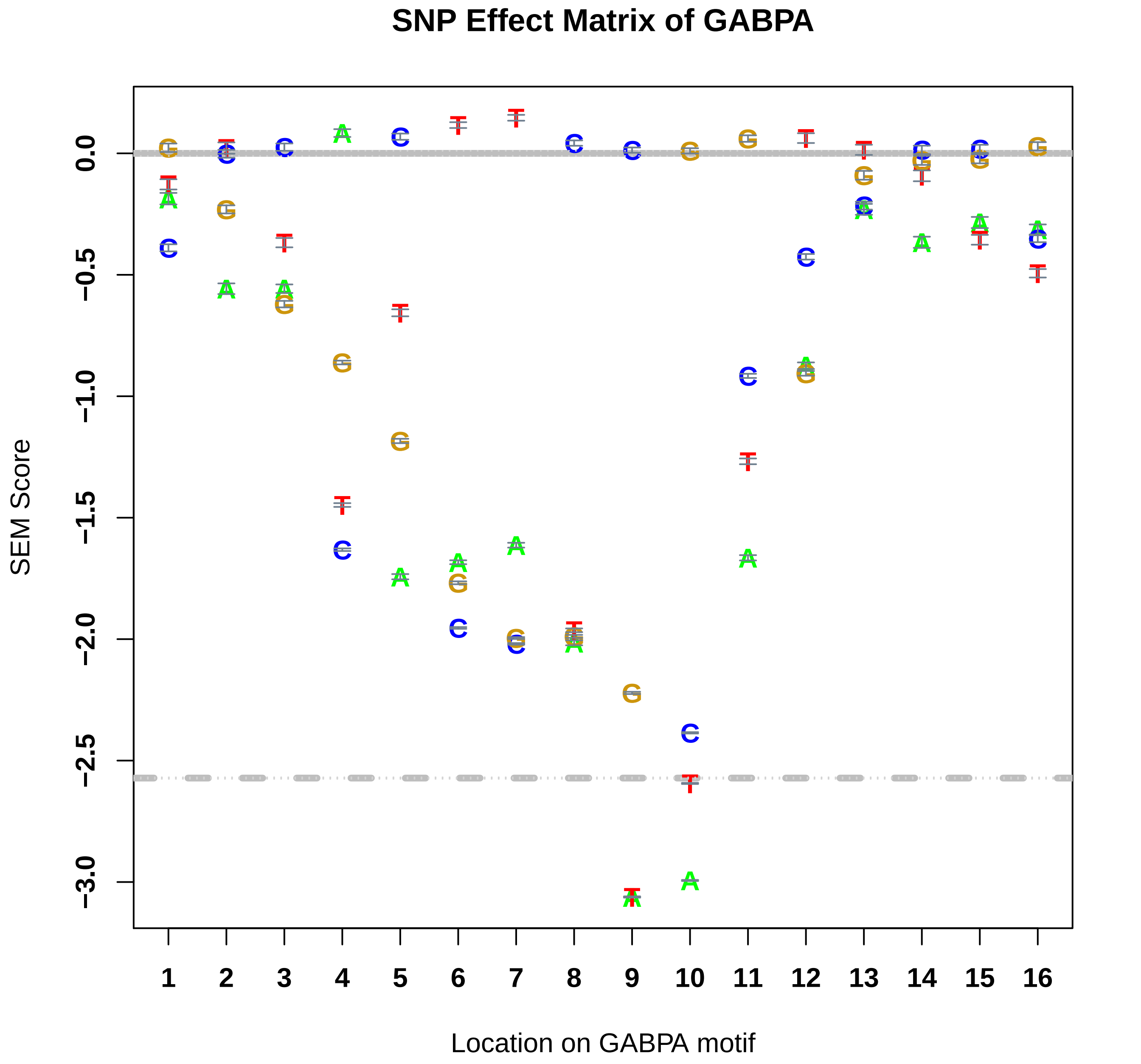

### Klf7_primary

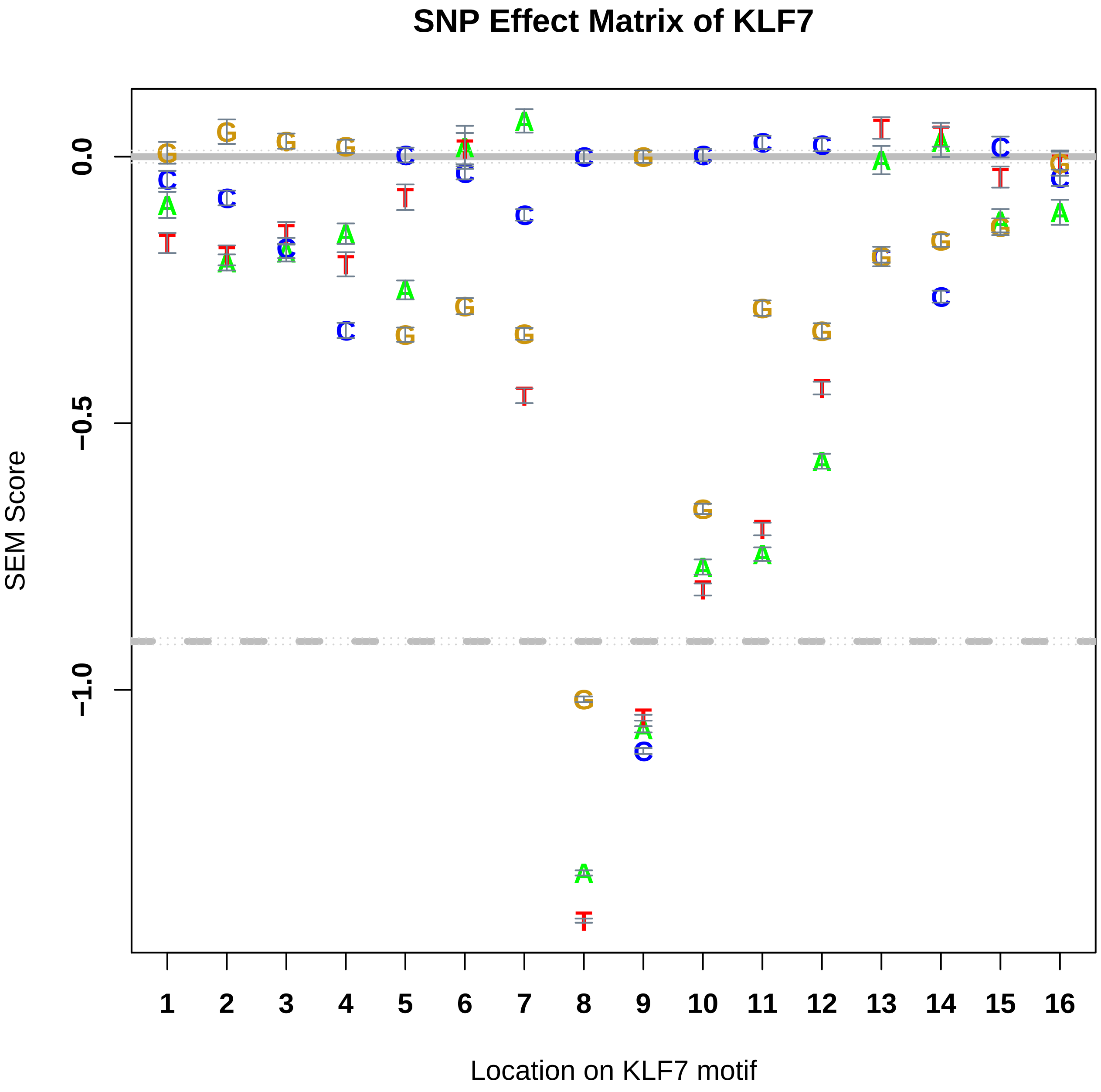

### KLF14_1

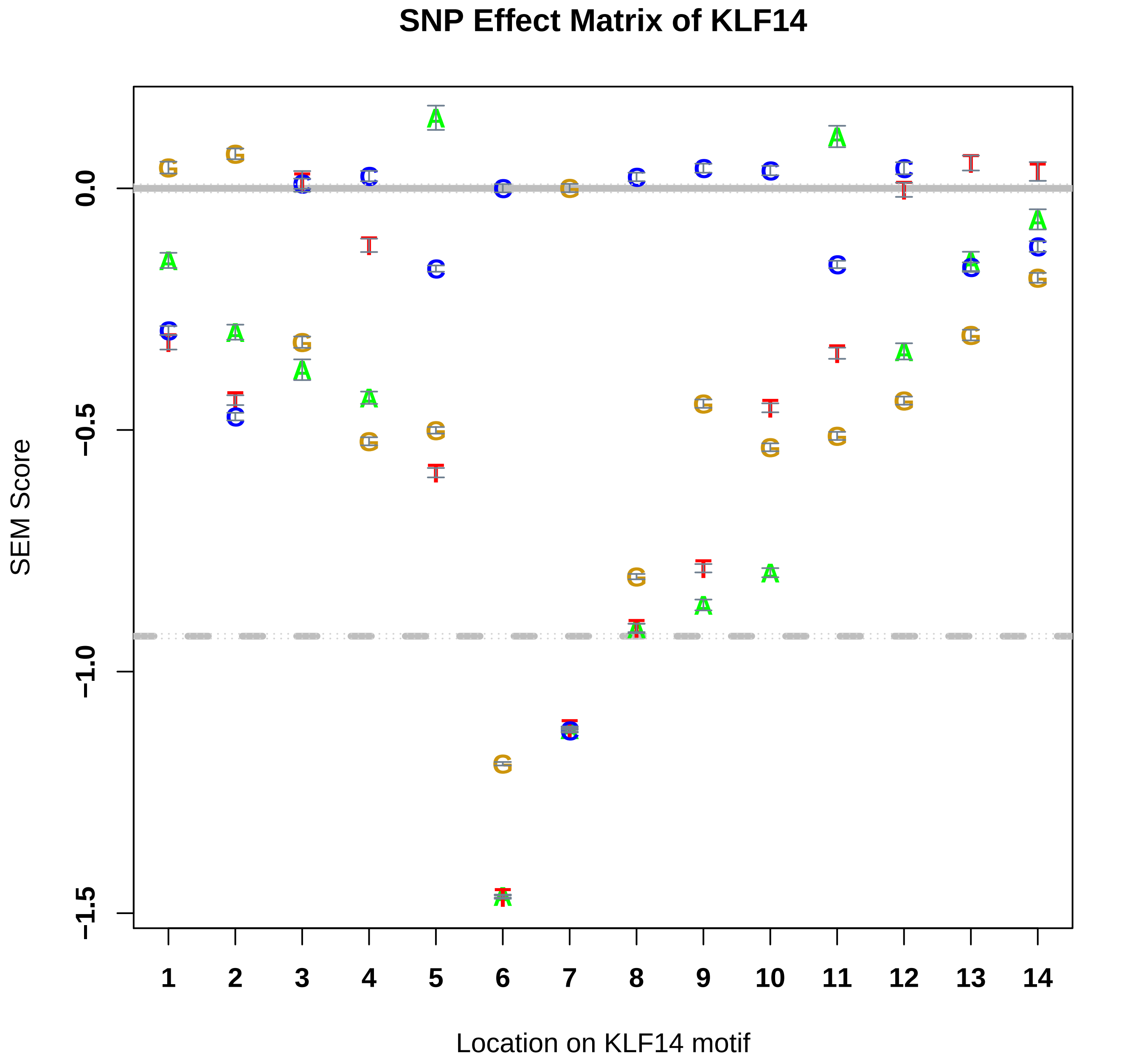

### M00007

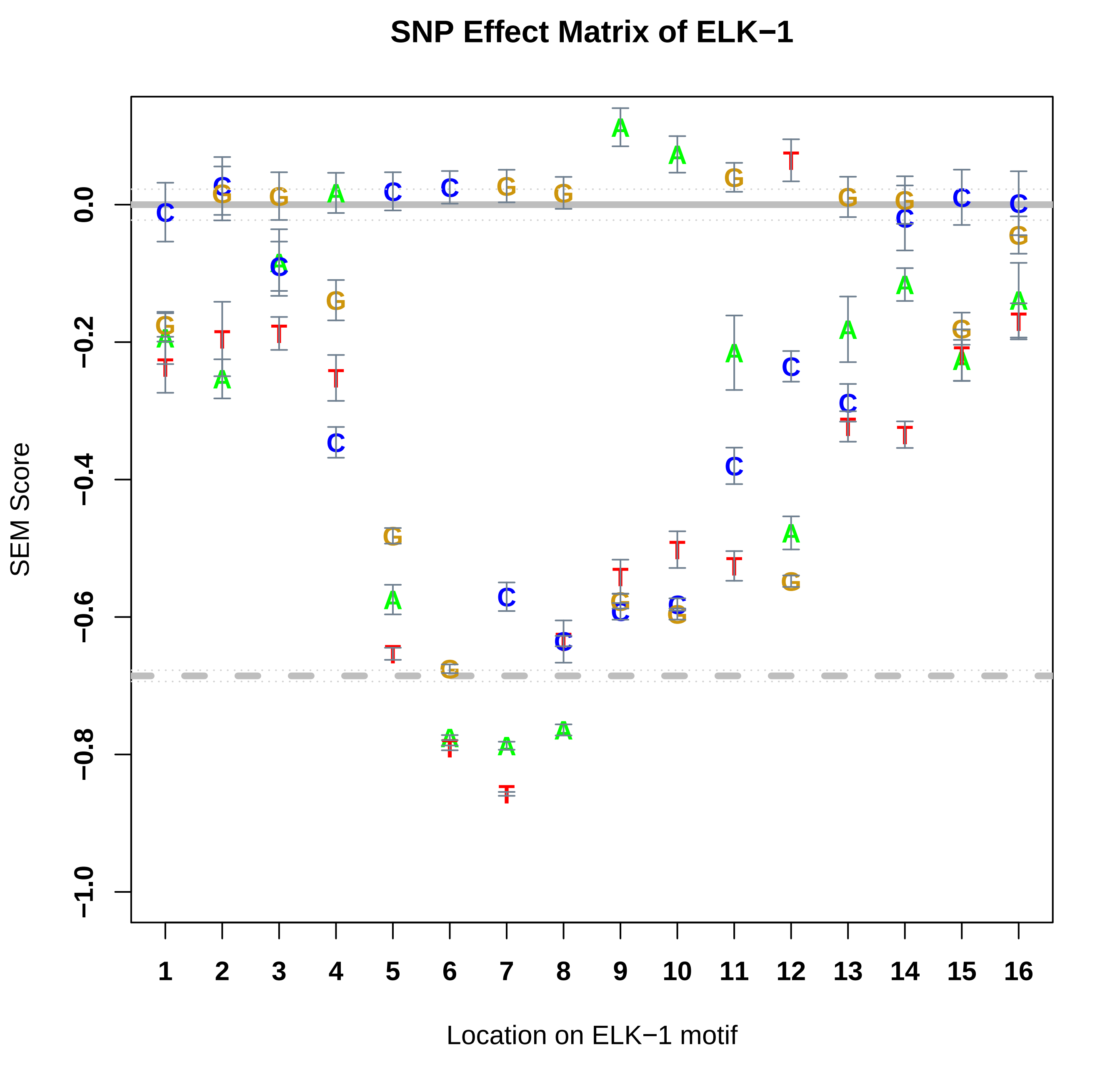

### M00008

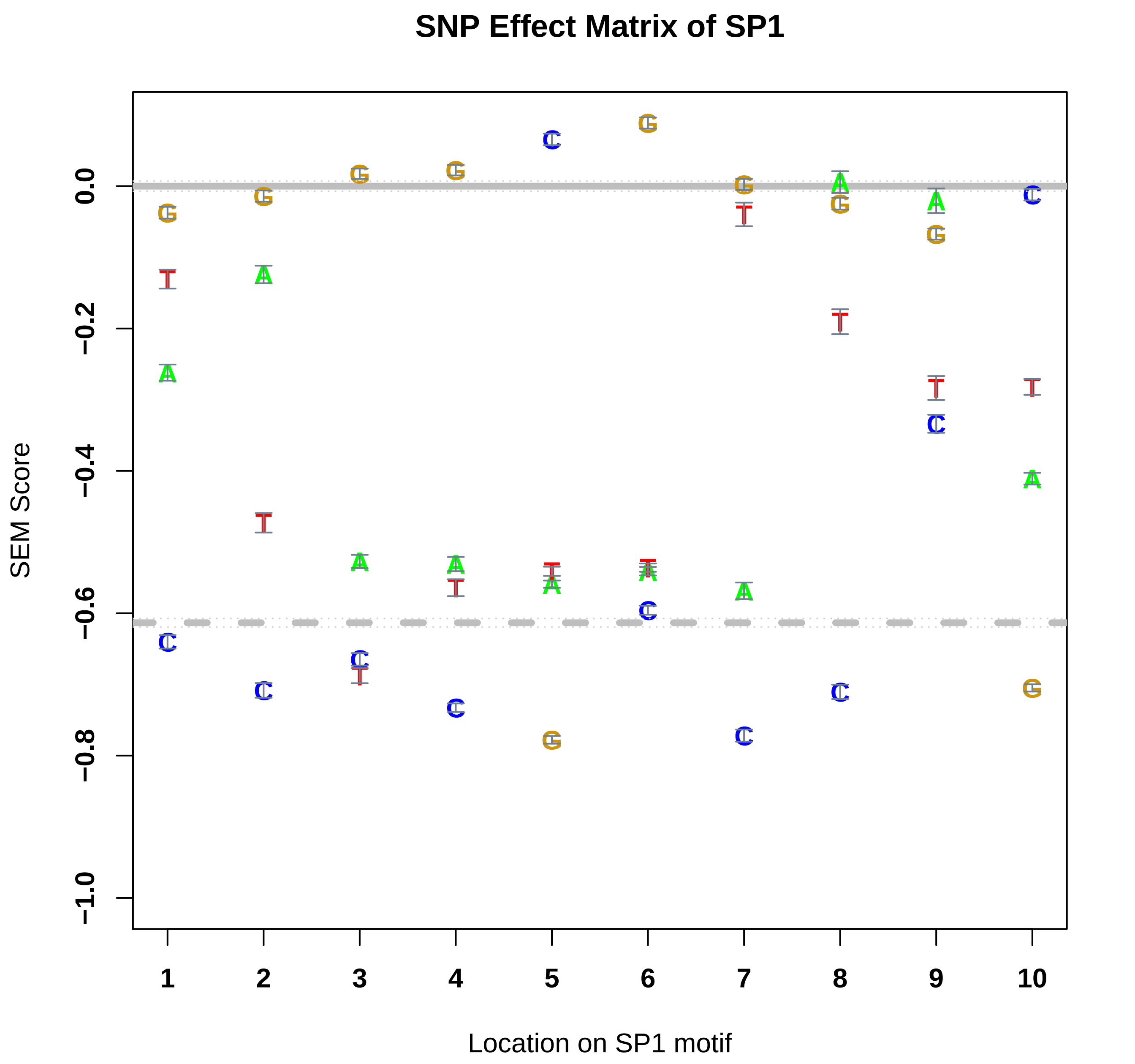

### M00039

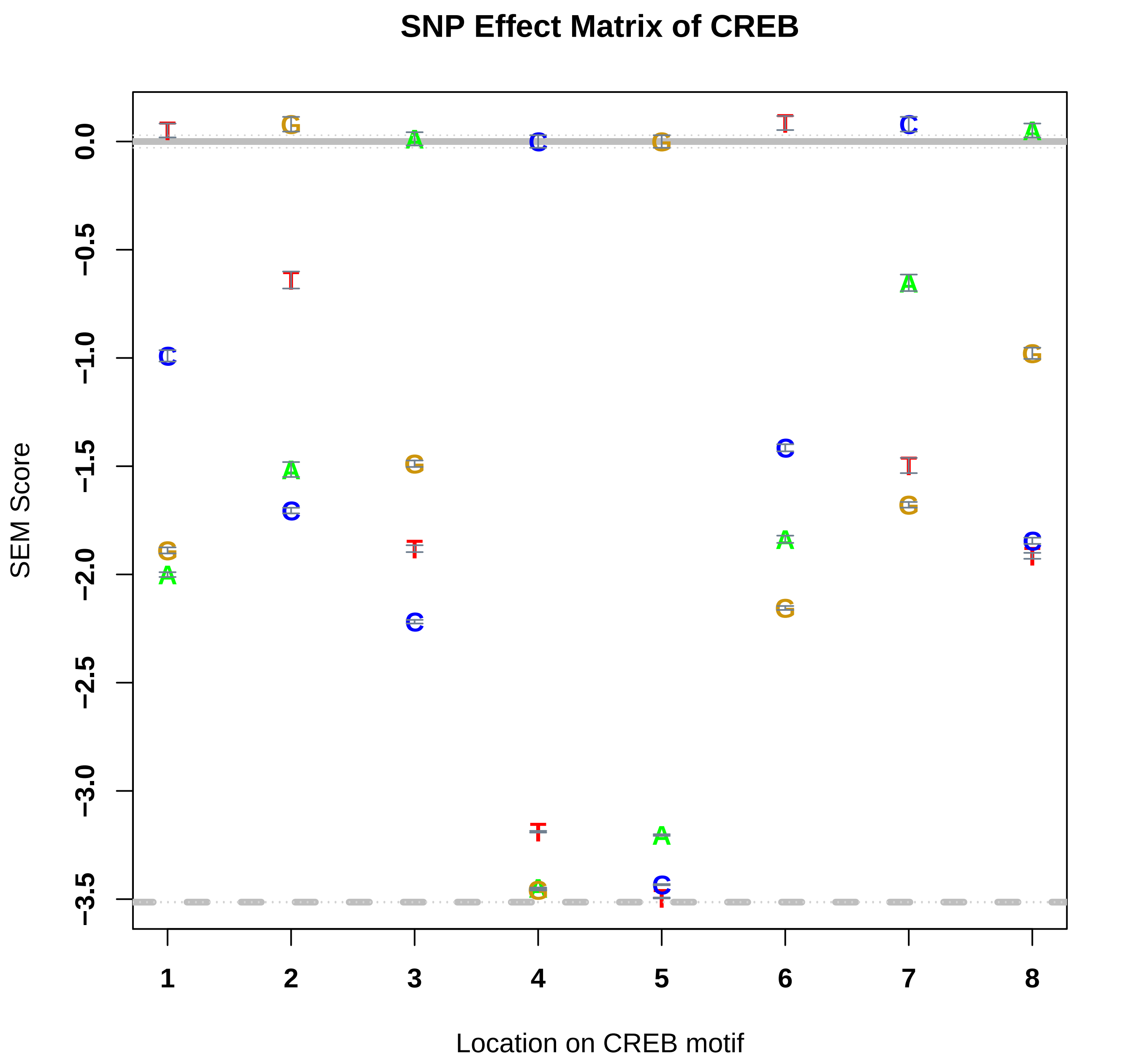

### M00040

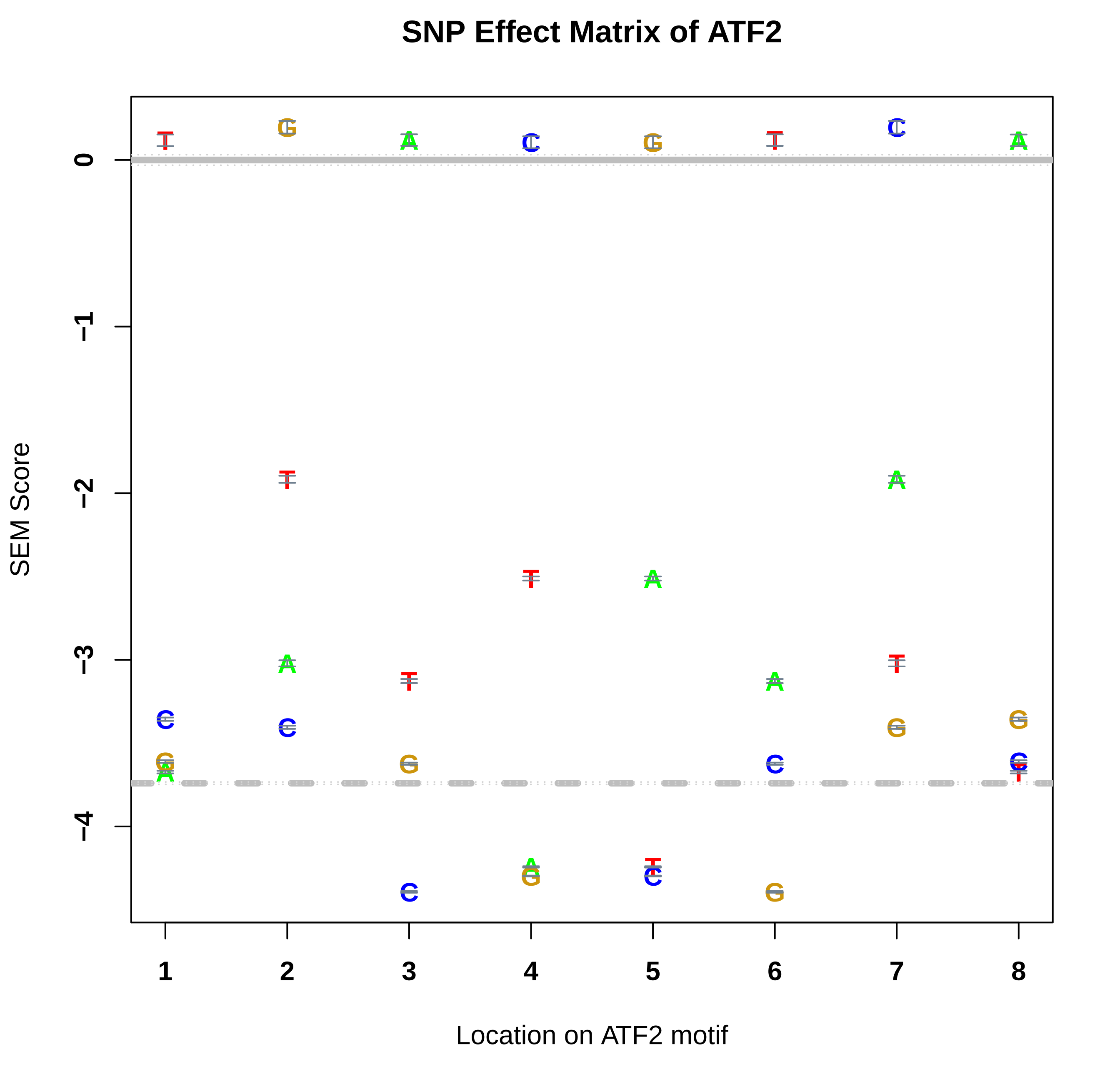

### M00041

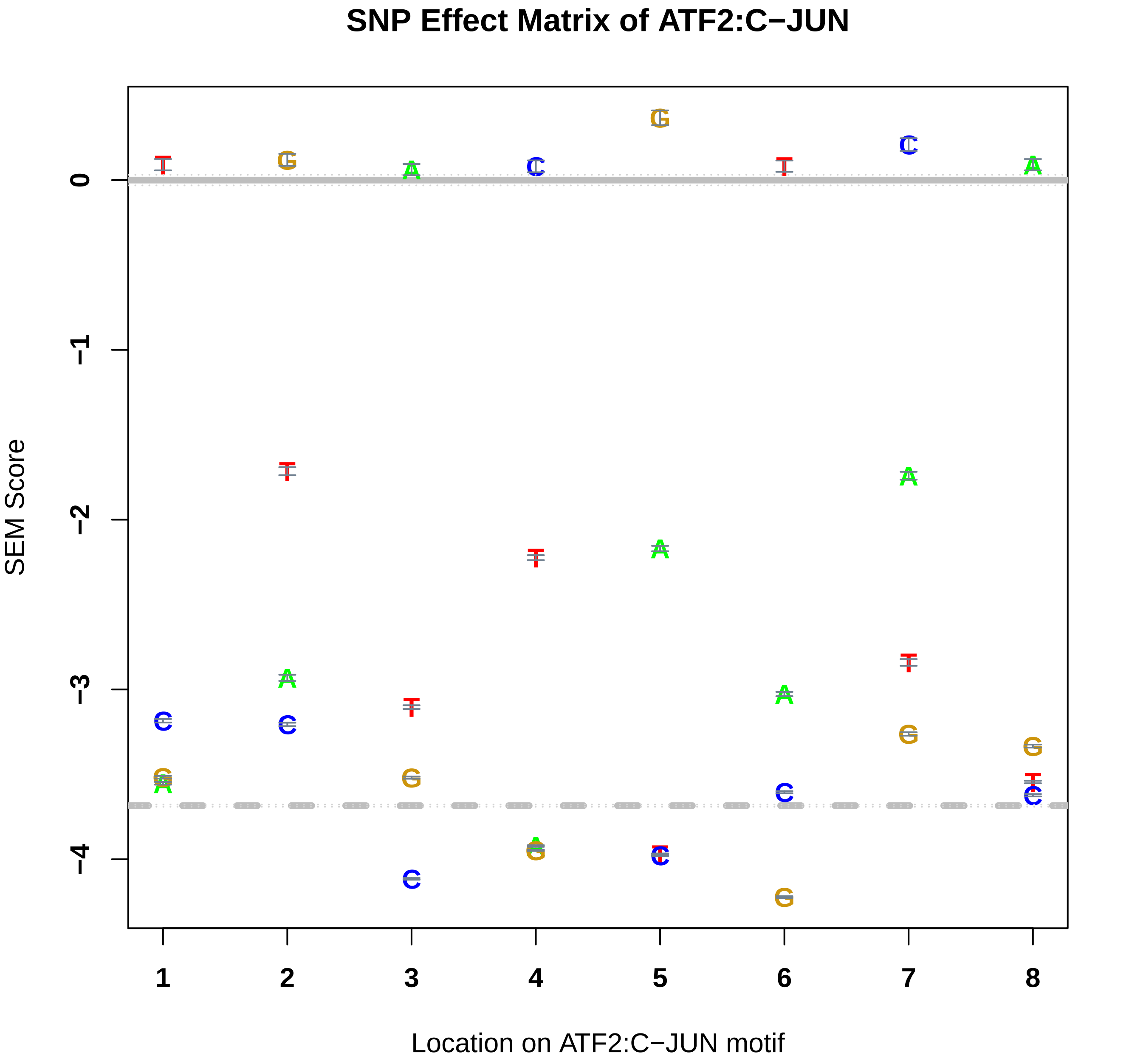

### M00065

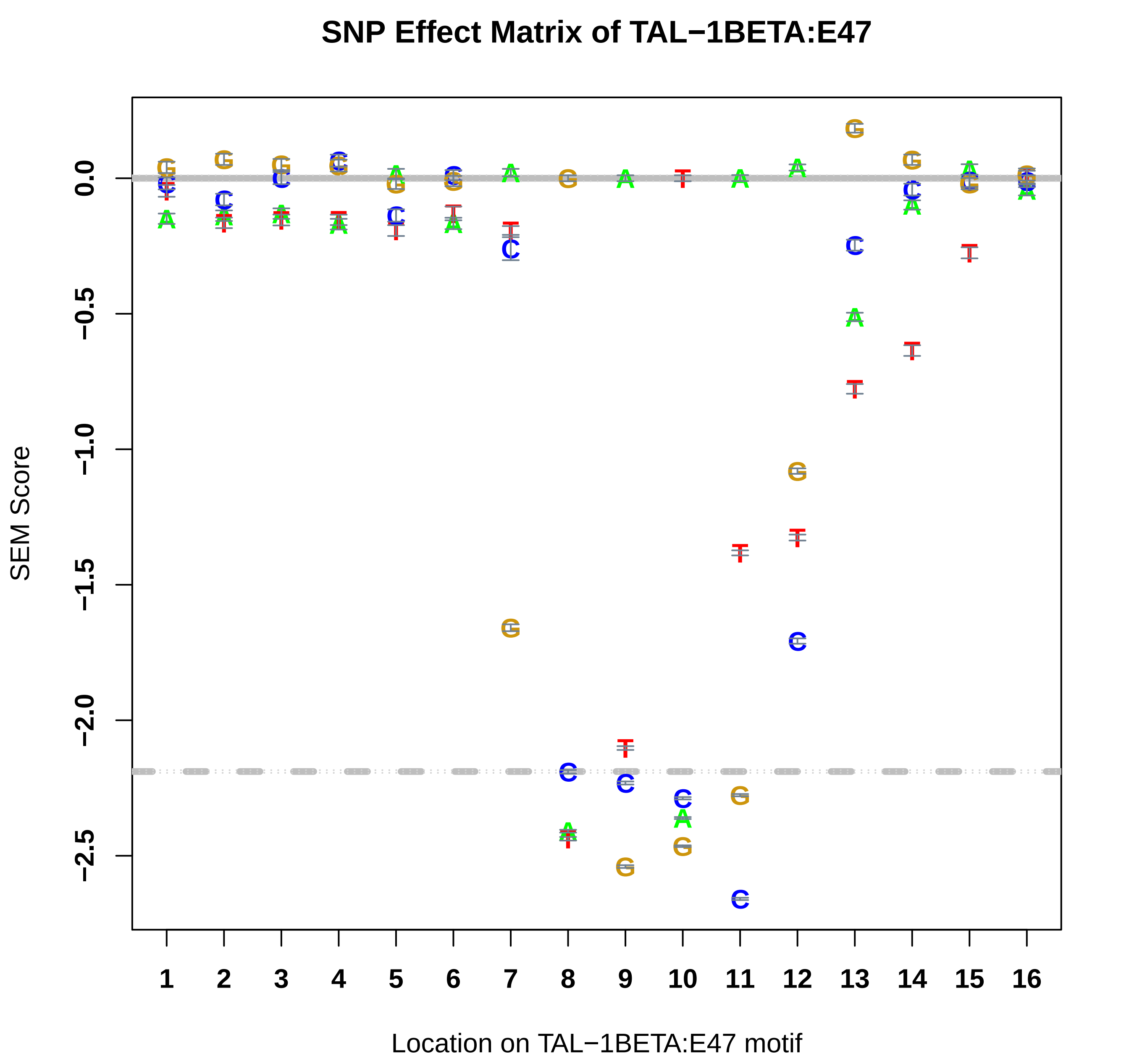

### M00066

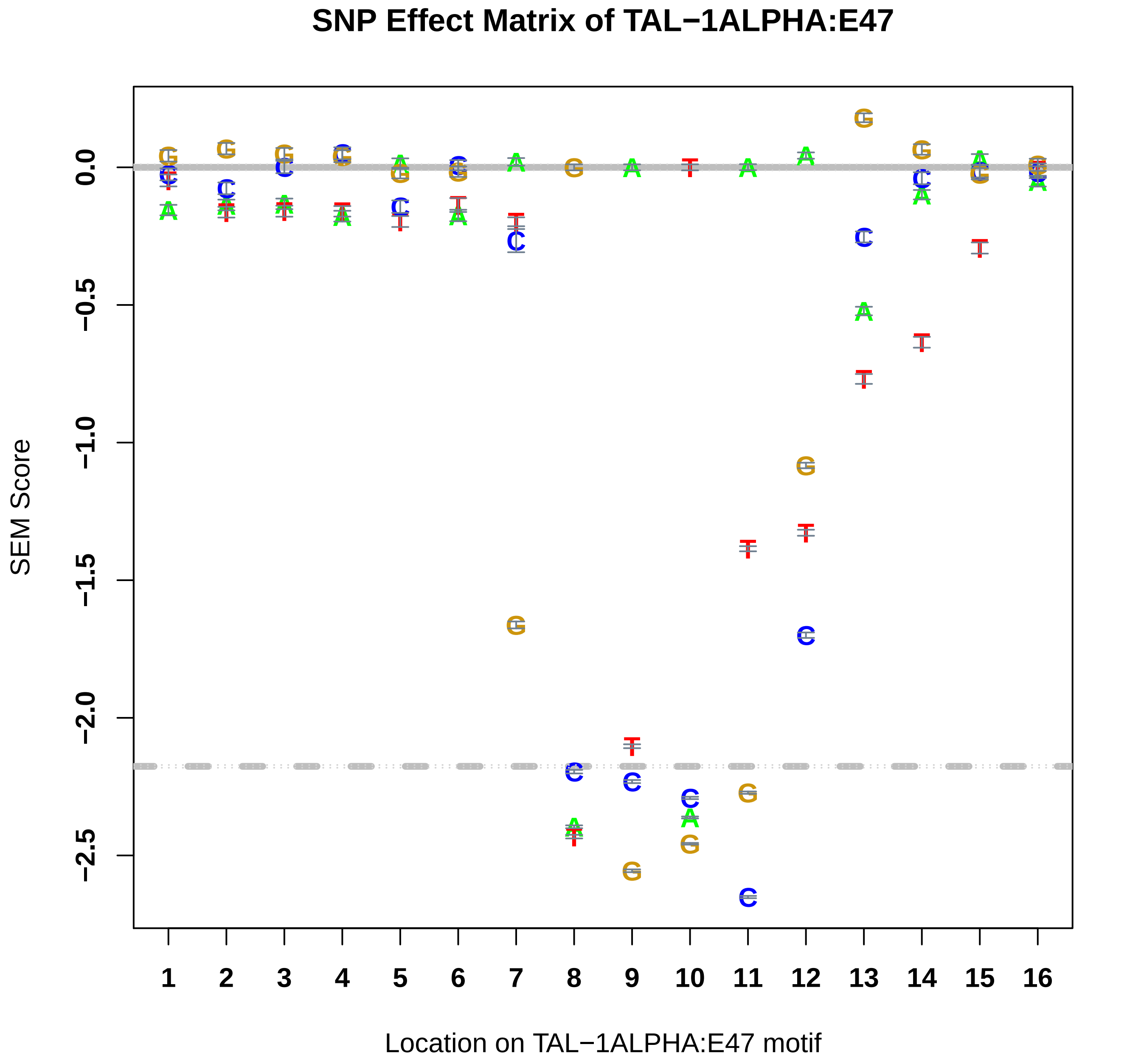

### M00070

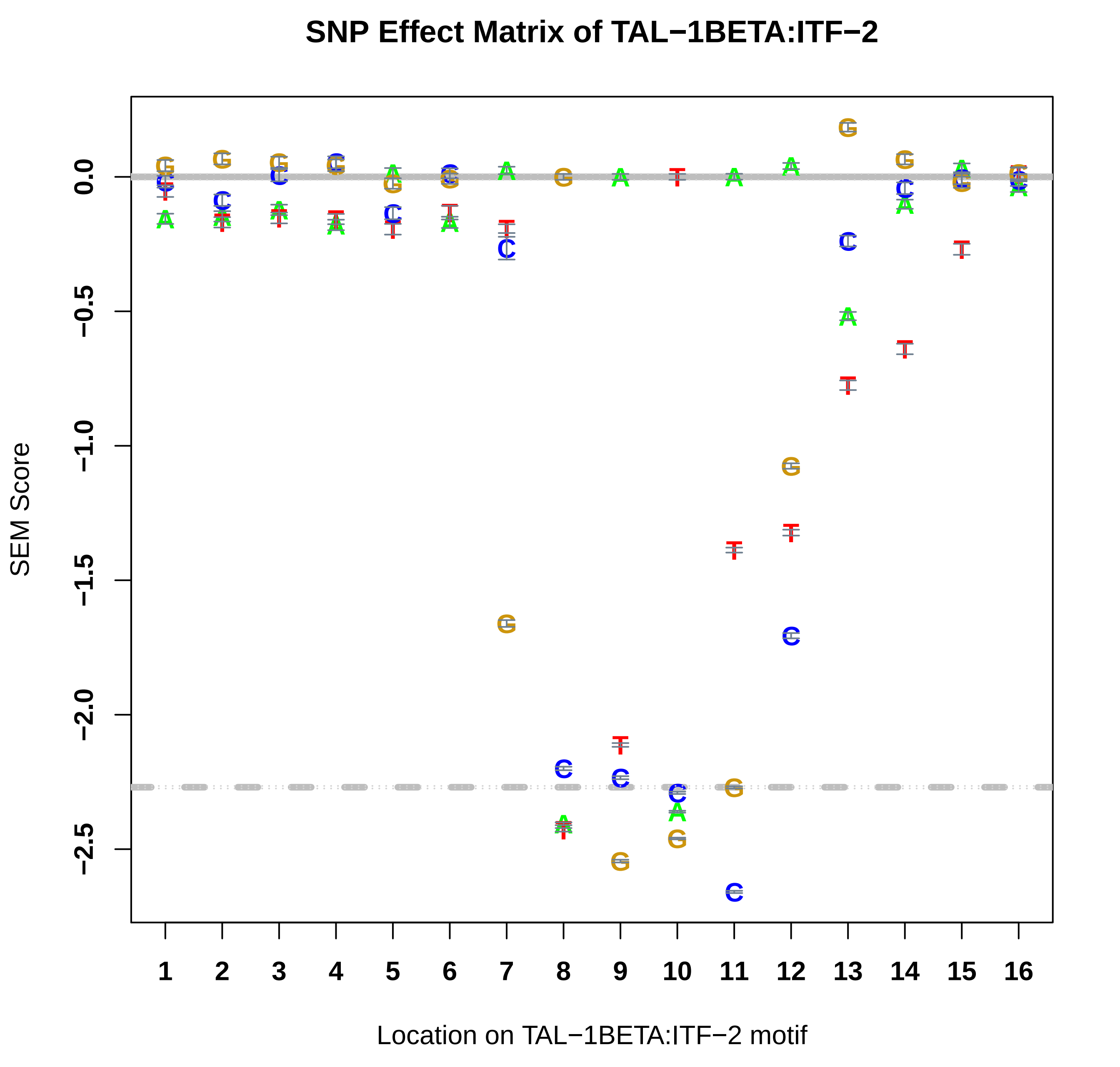

### M00075

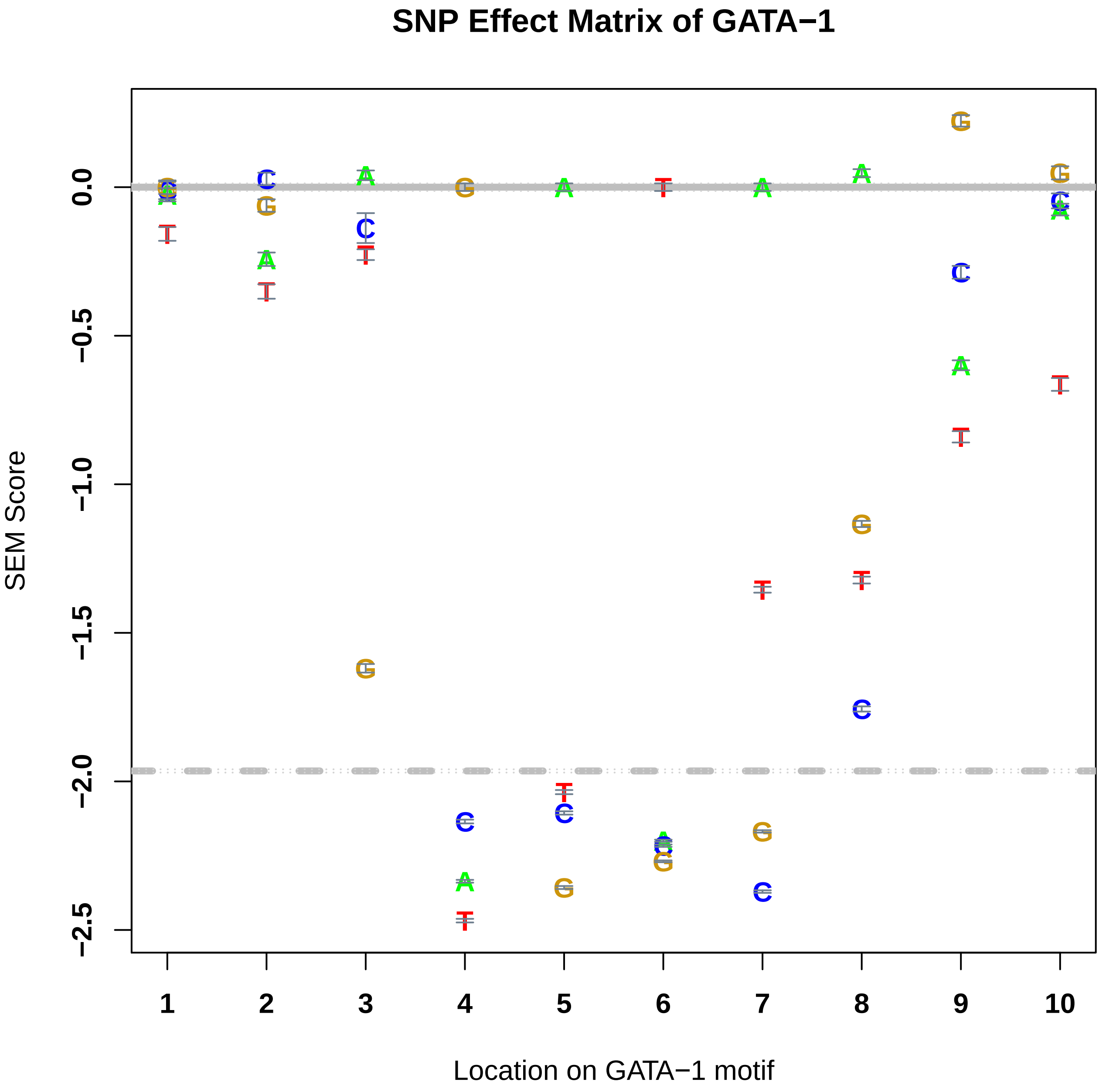

### M00076

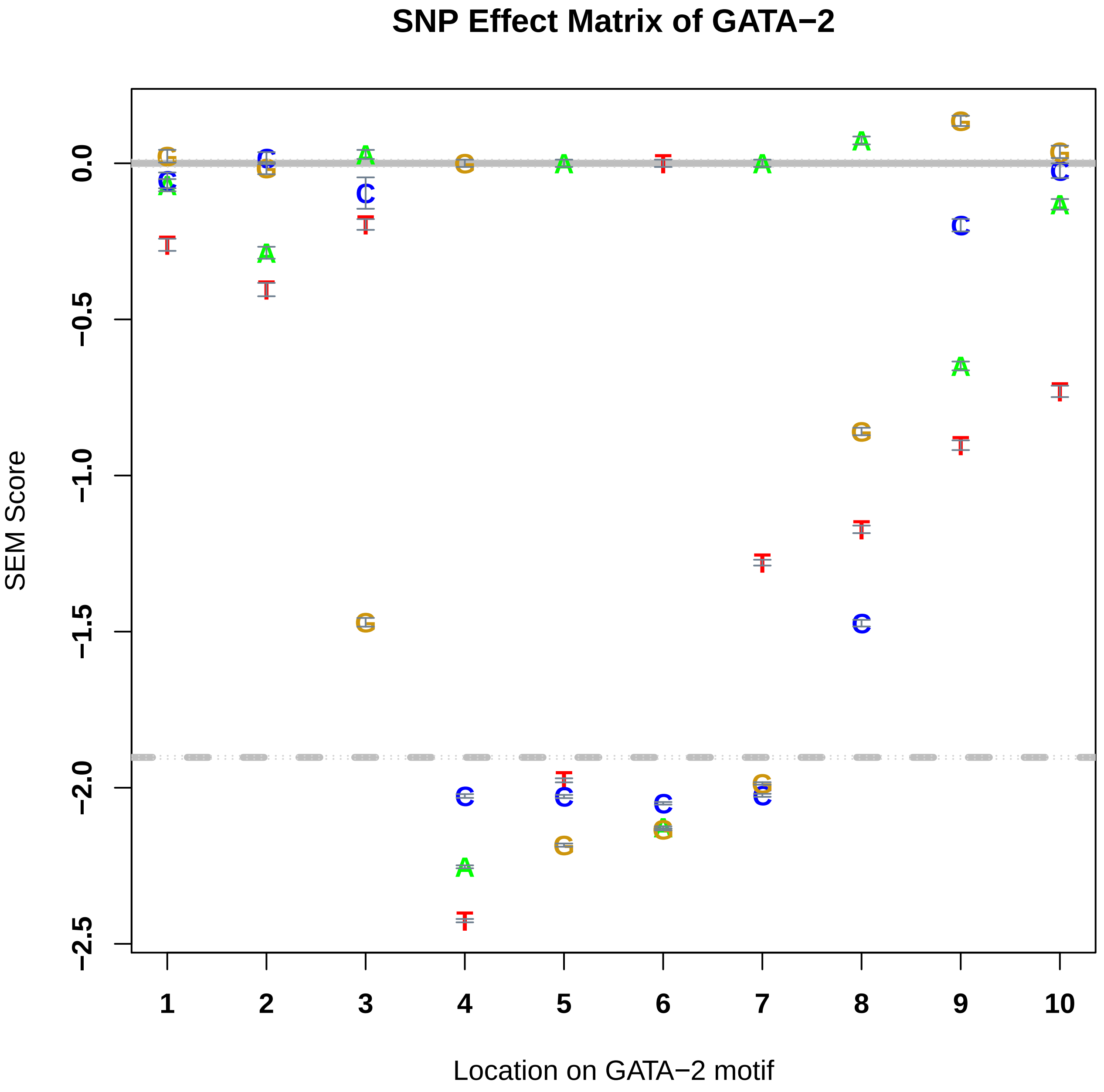

### M00146

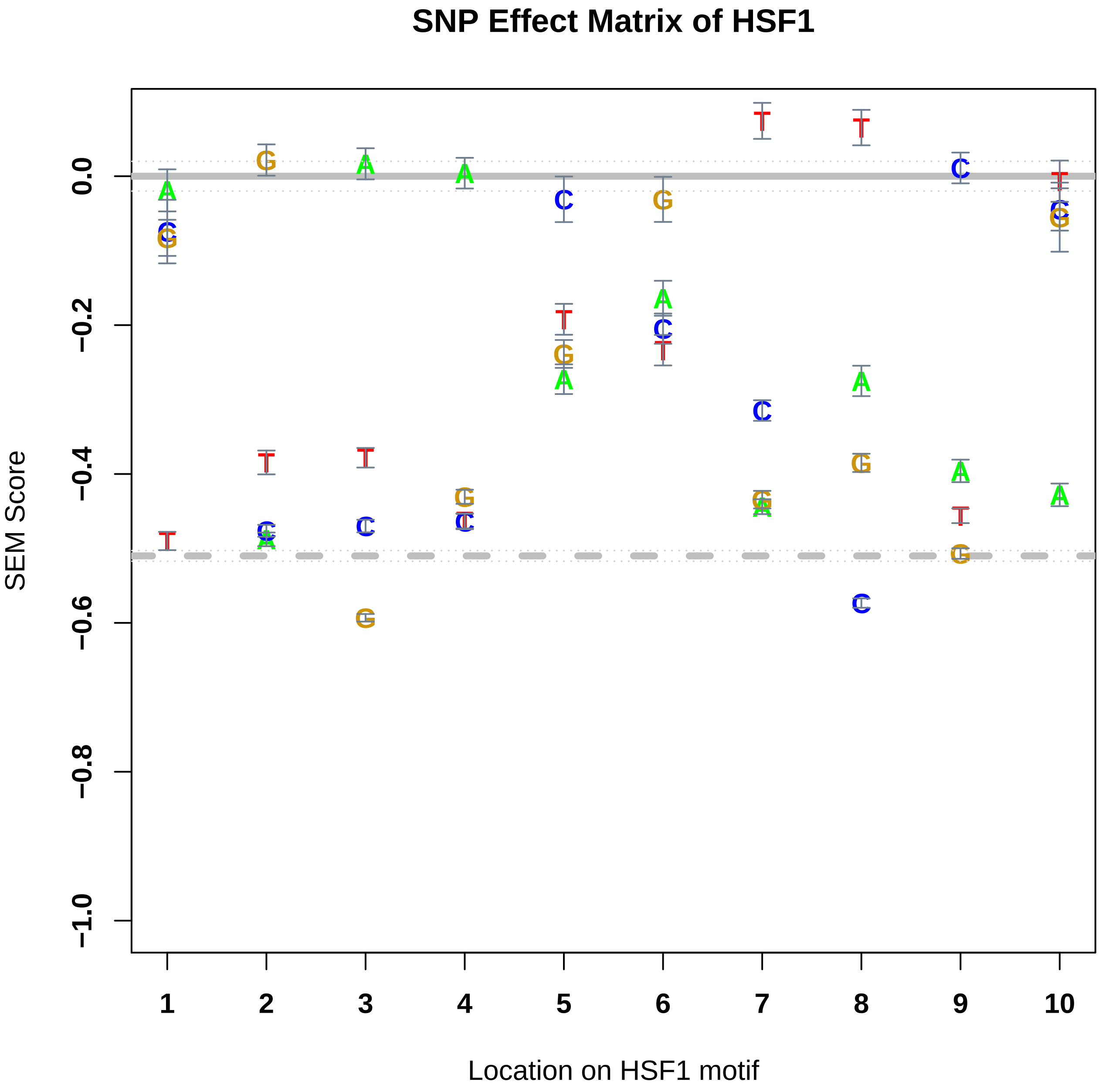

### M00172

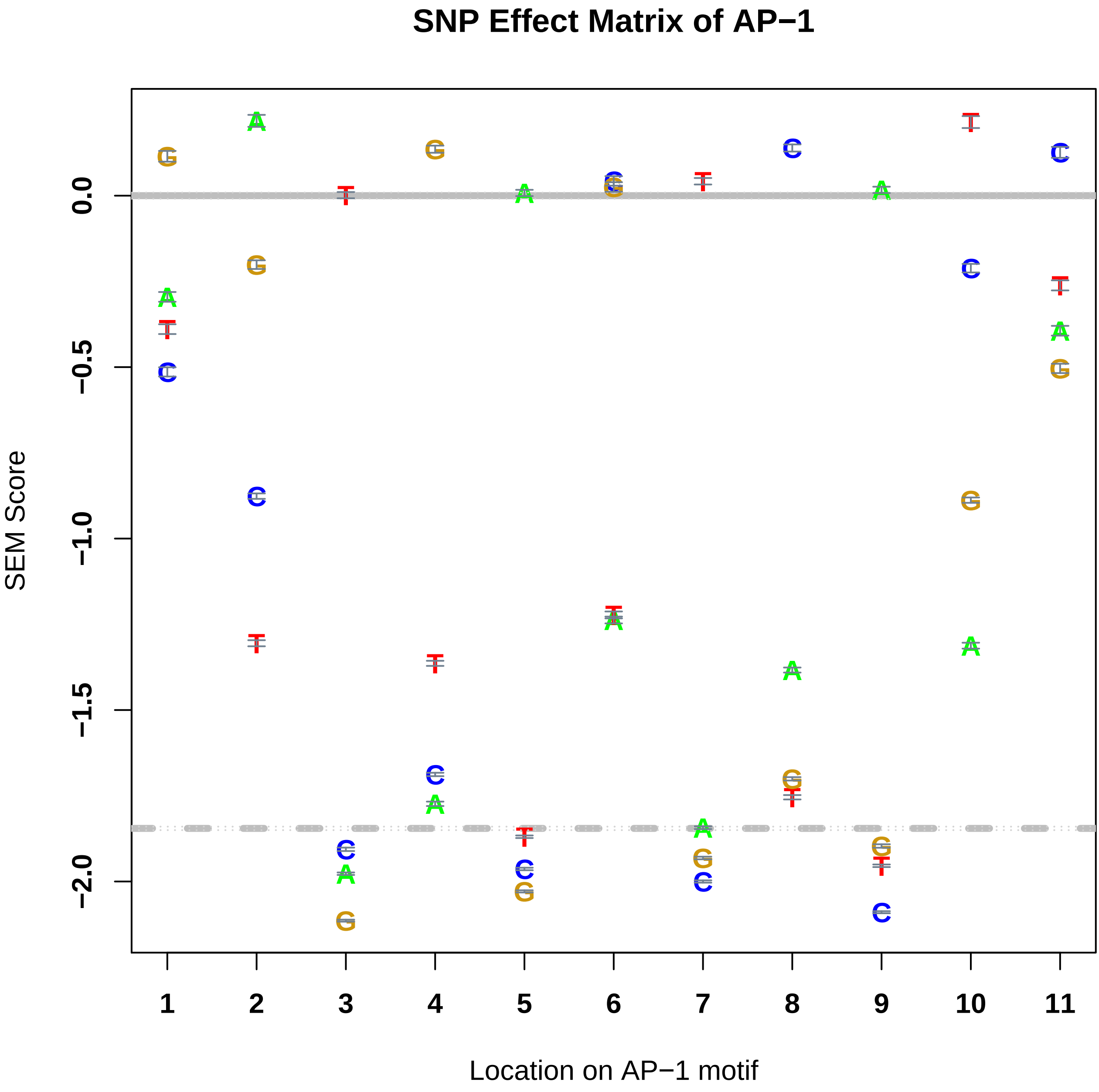

### M00173

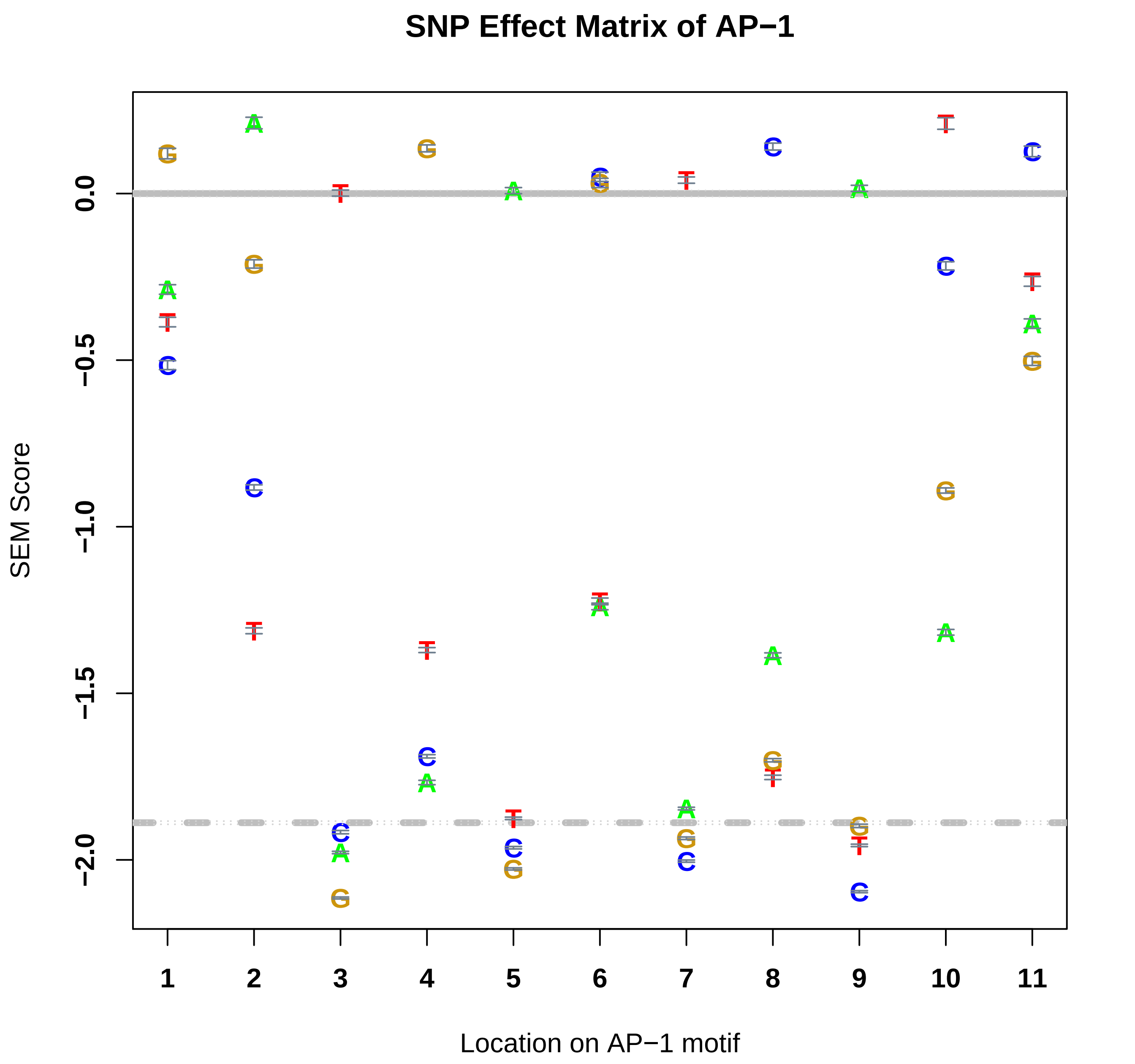

### M00174

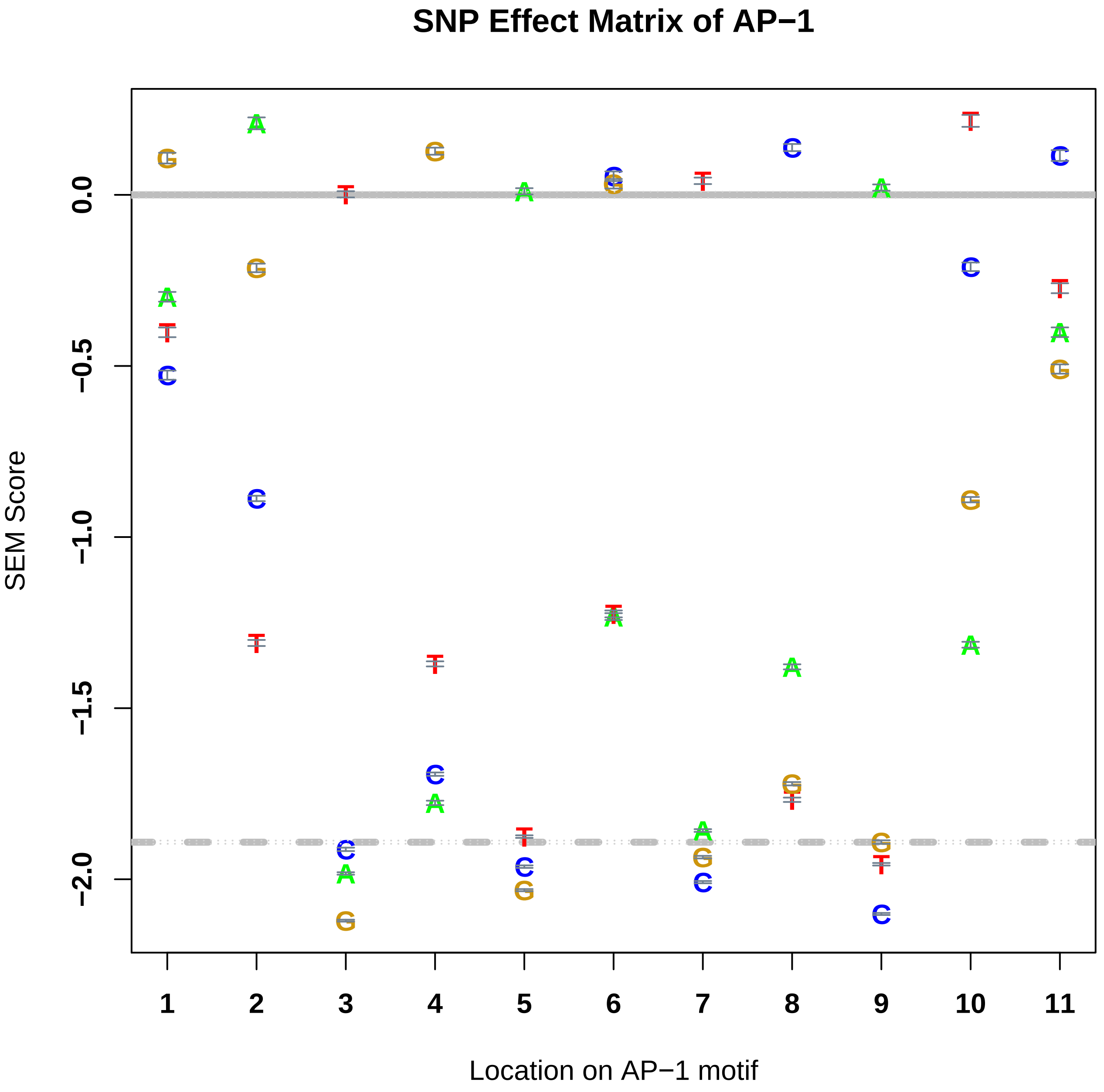

### M00183

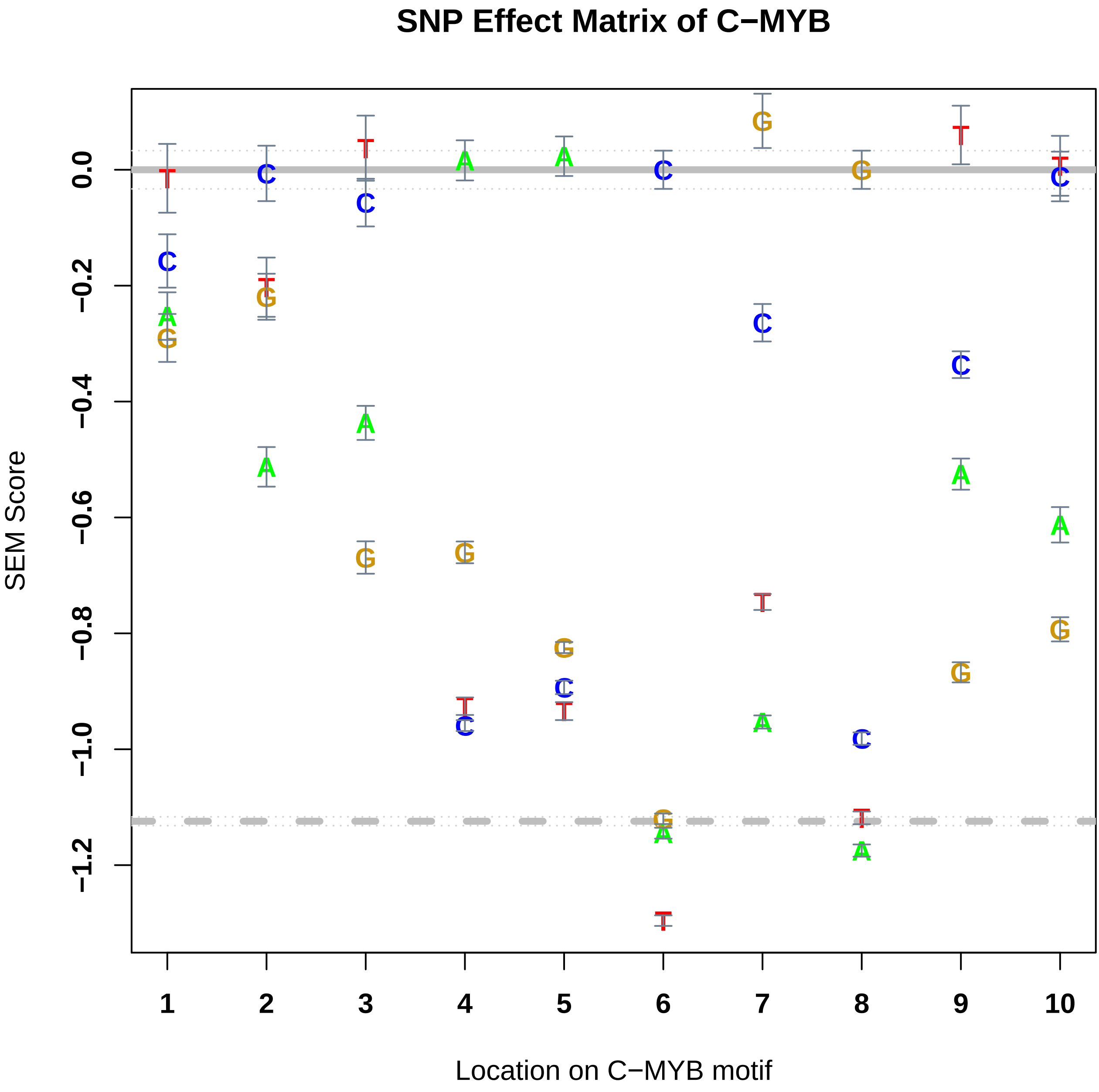

### M00185

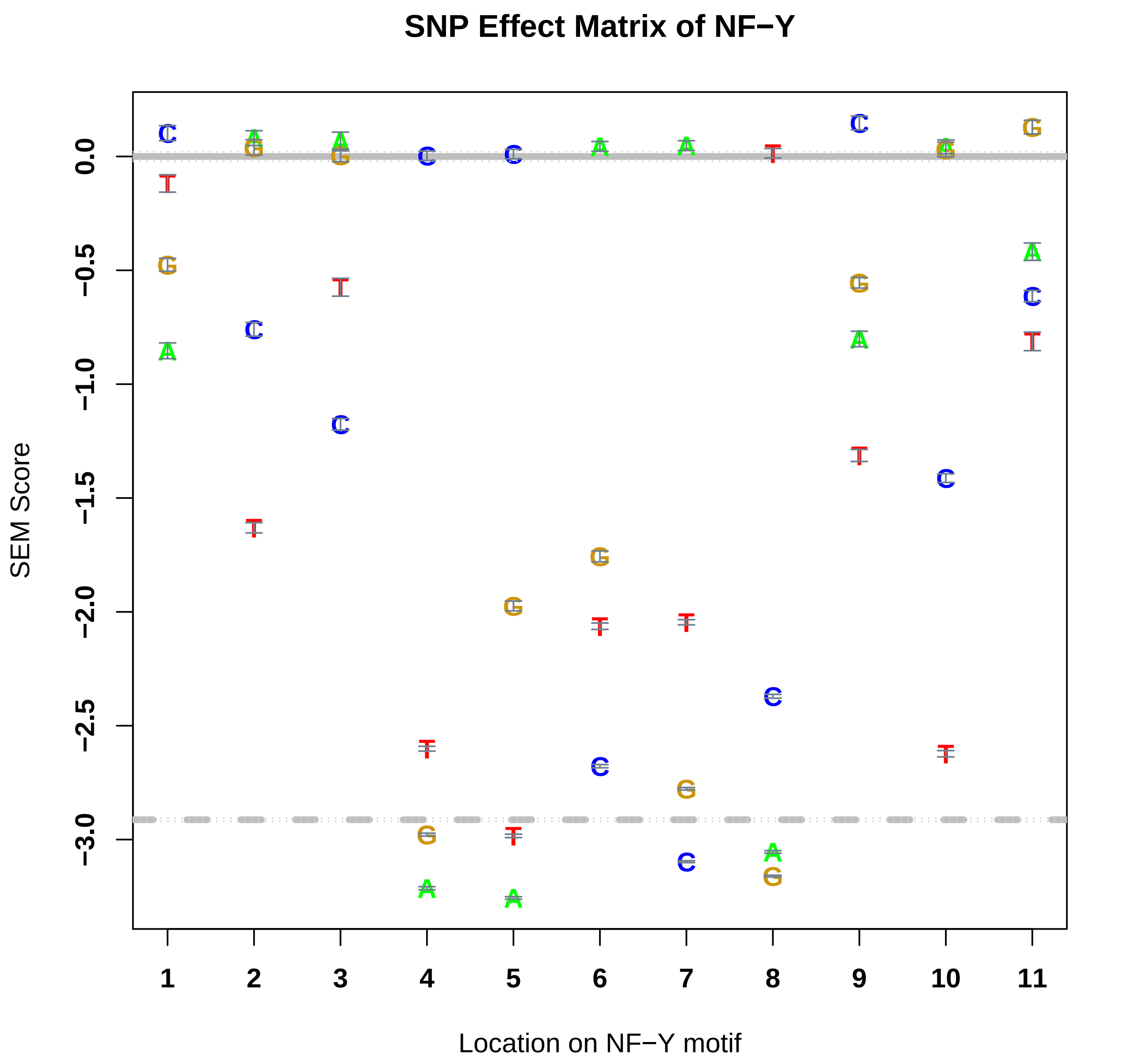

### M00187

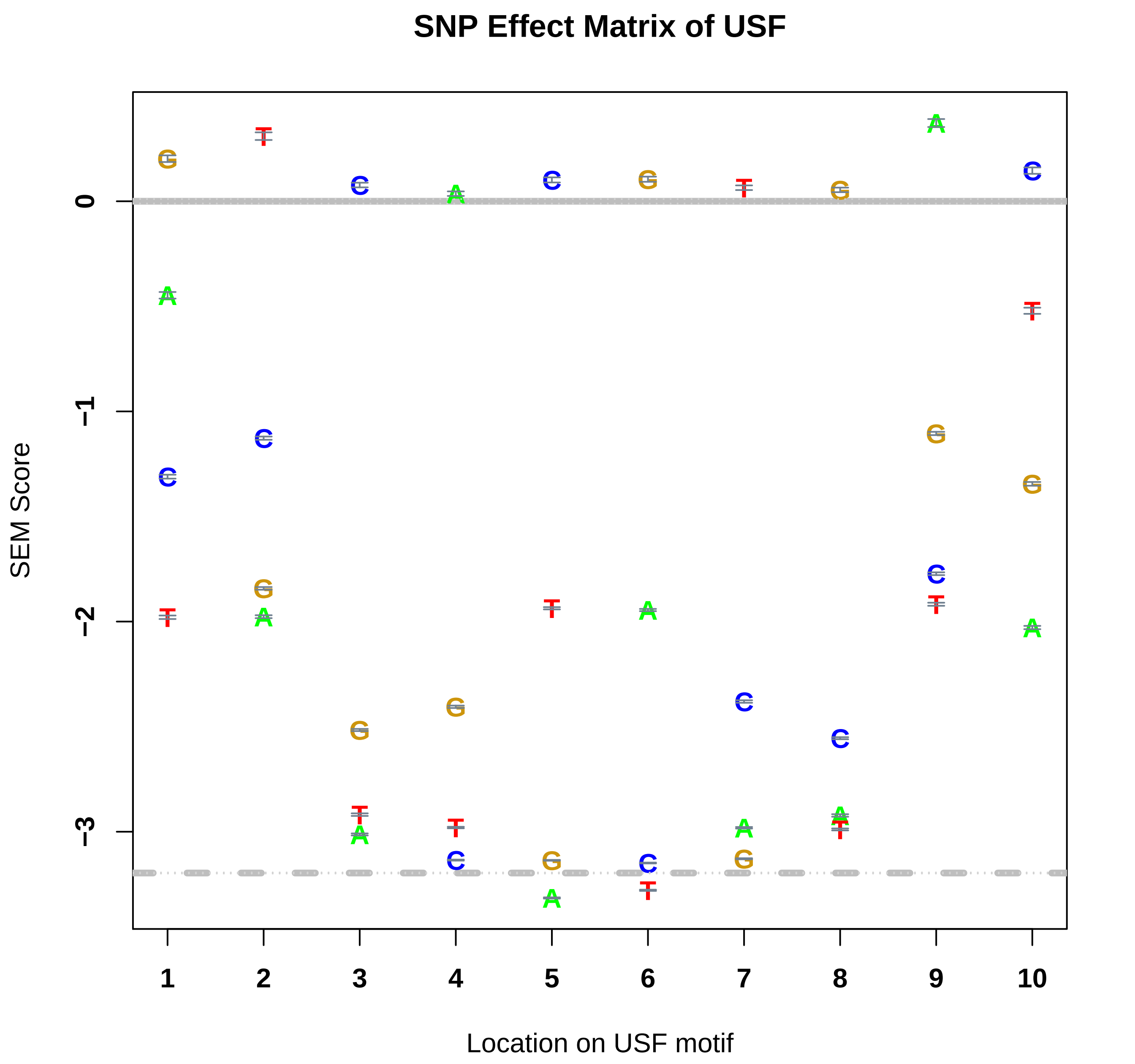

### M00188

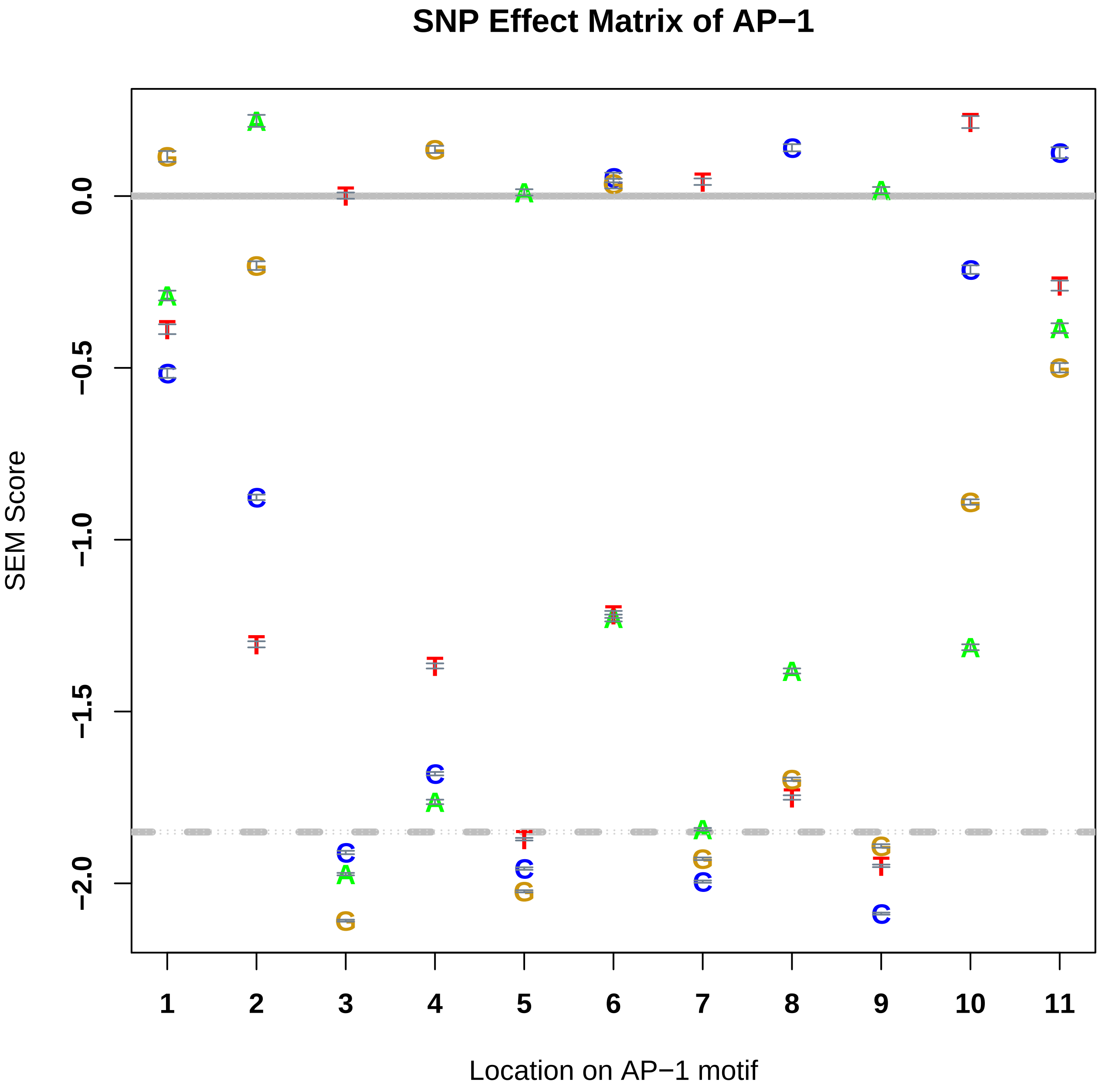

### M00189

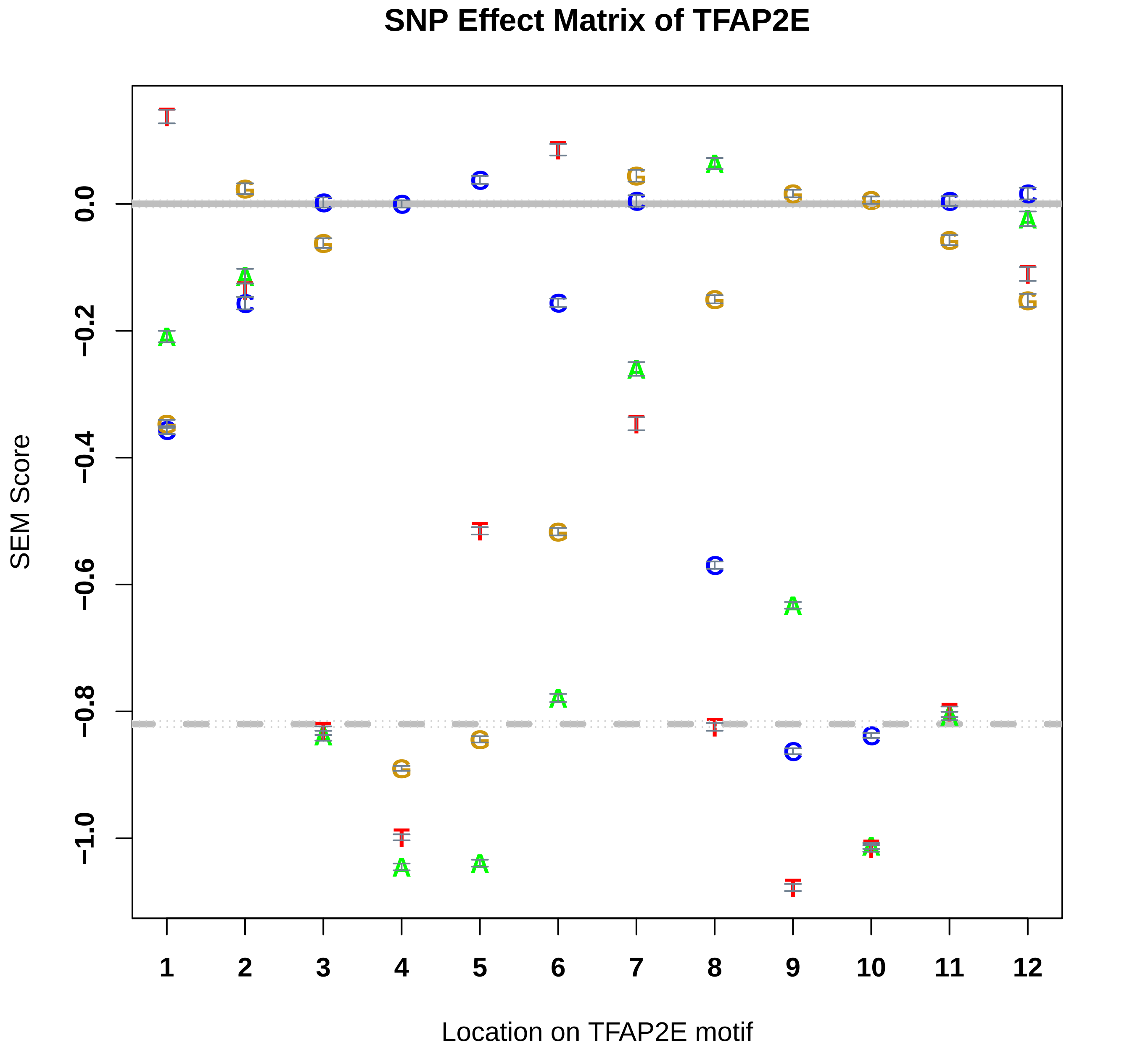

### M00196

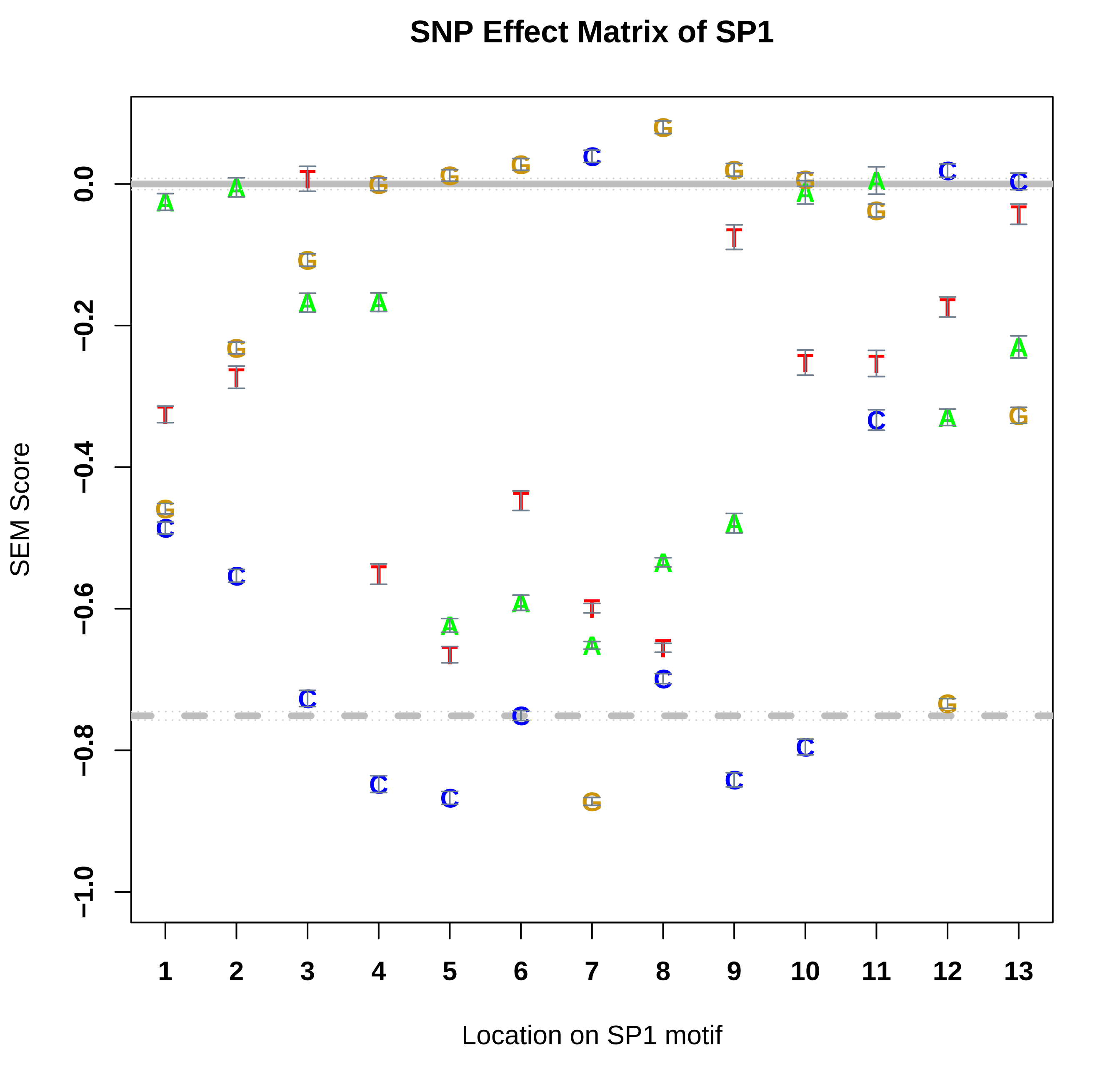

### M00220

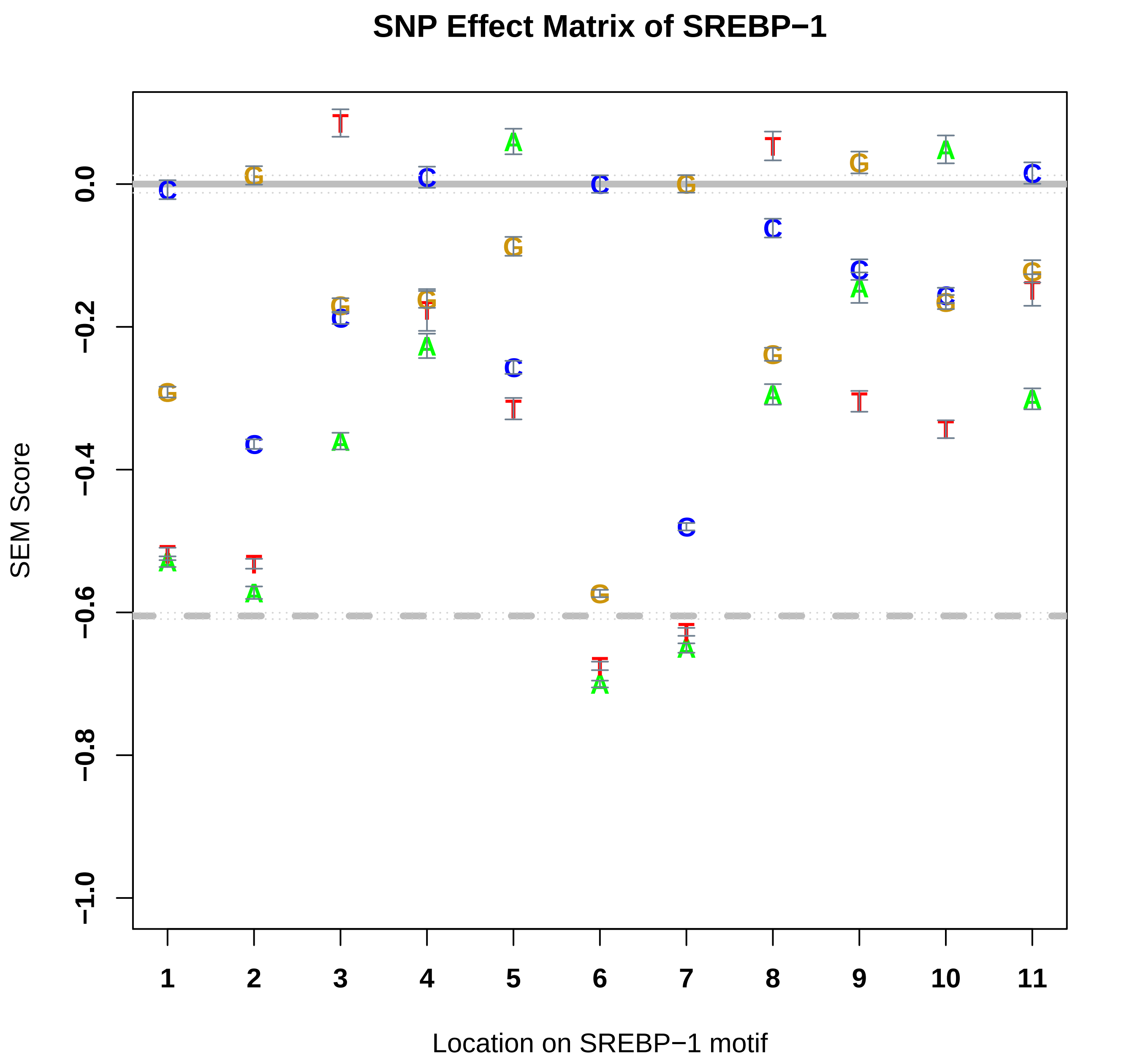

### M00221

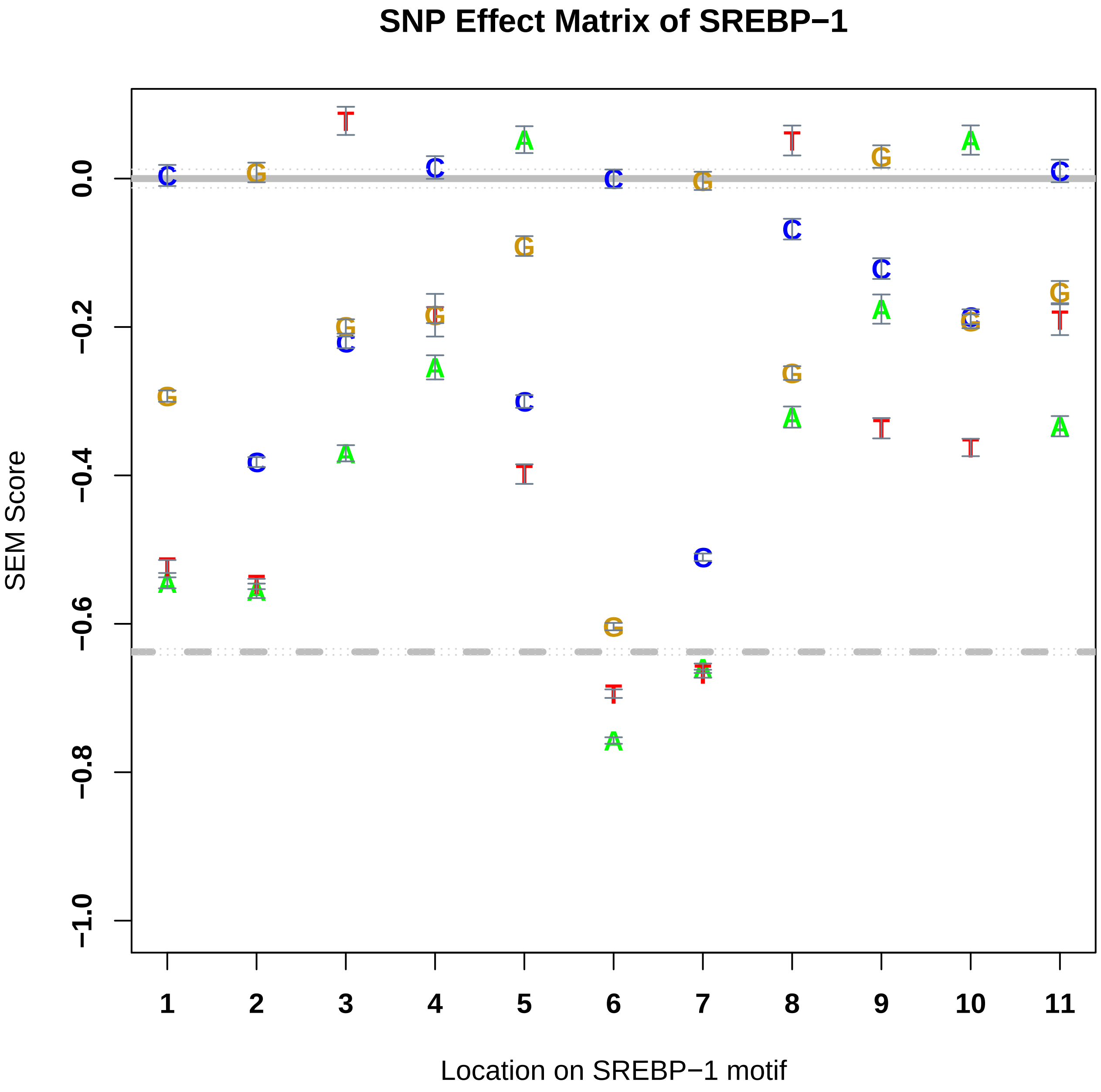

### M00322

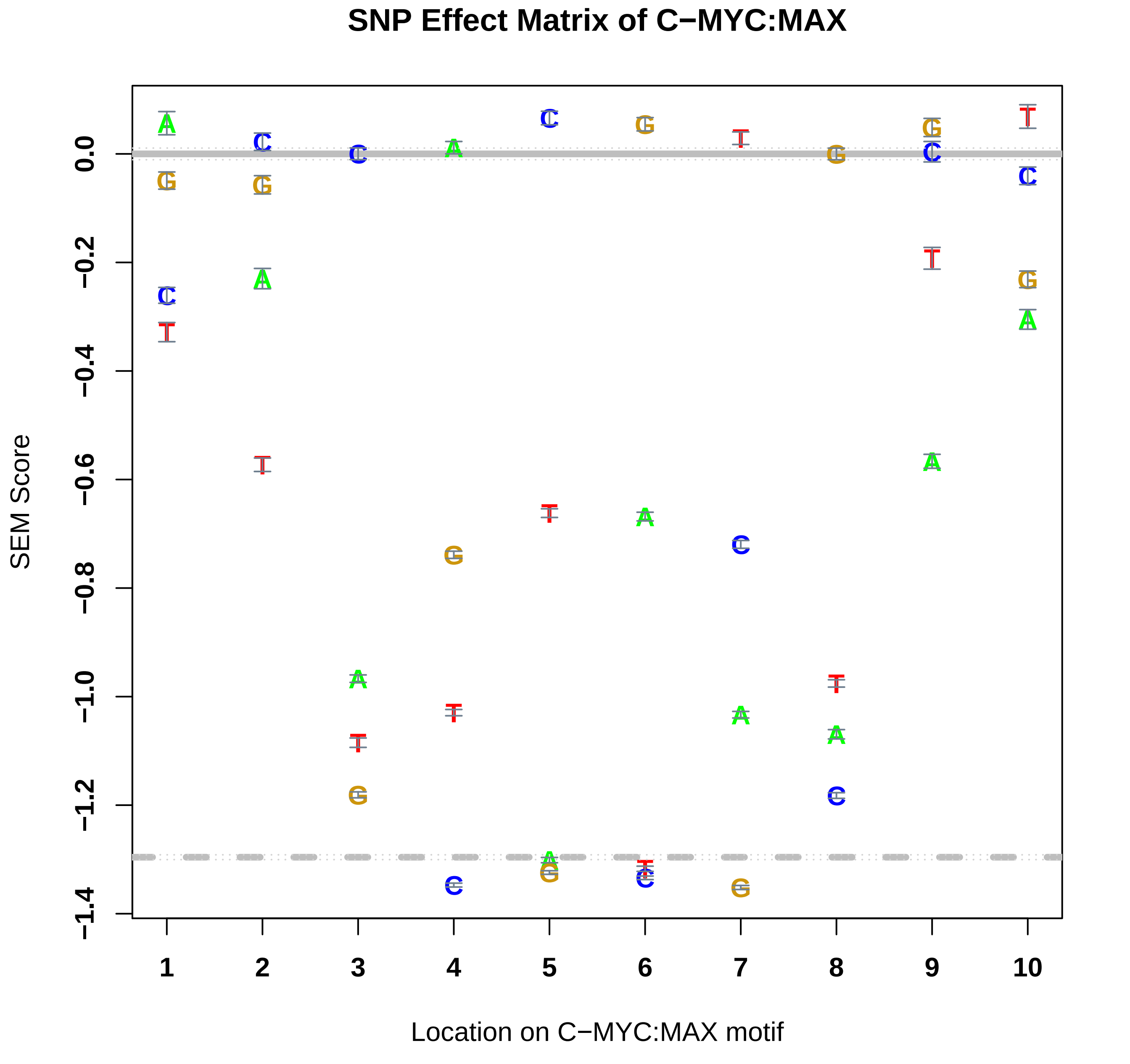

### M00346

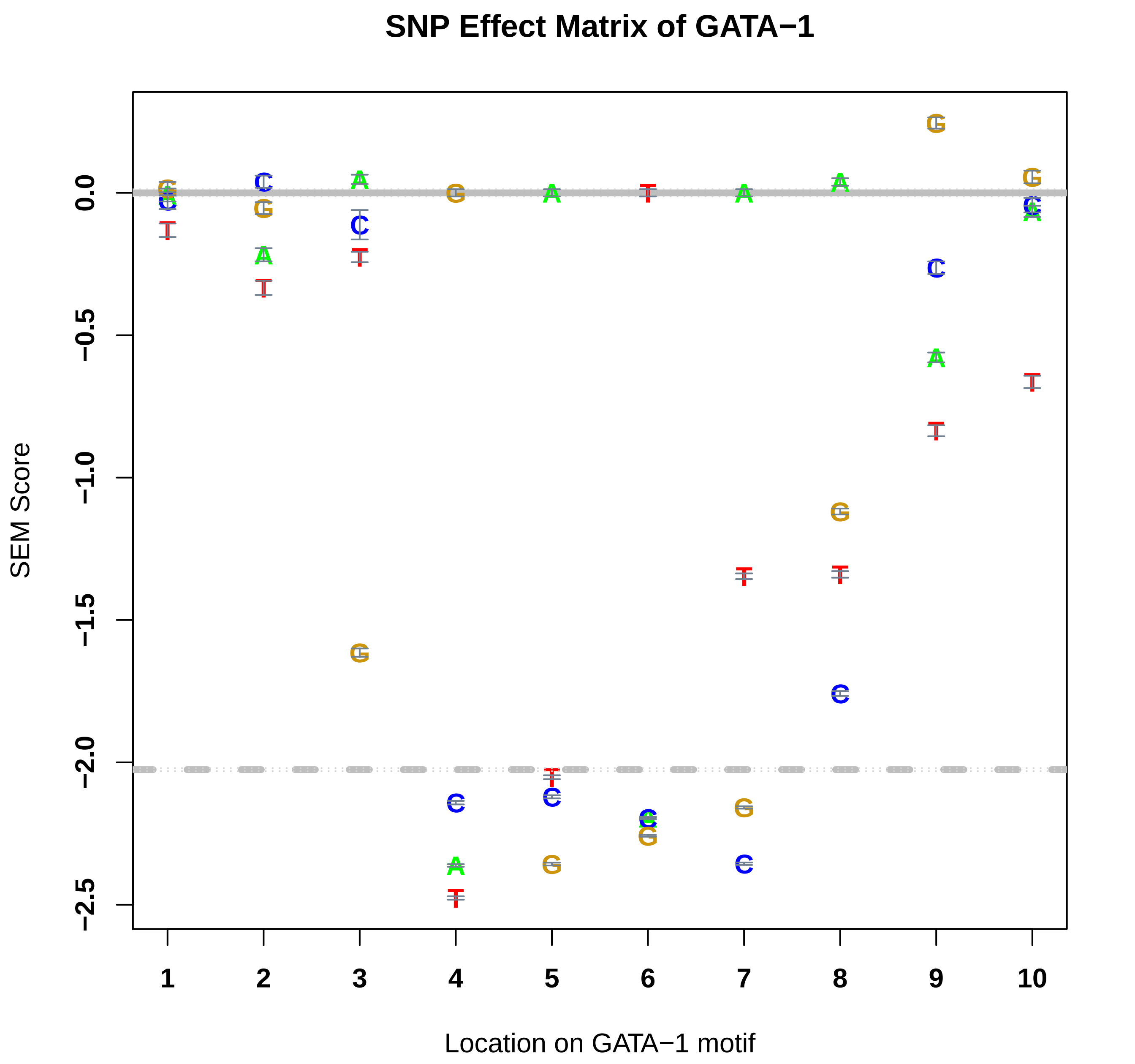

### M00347

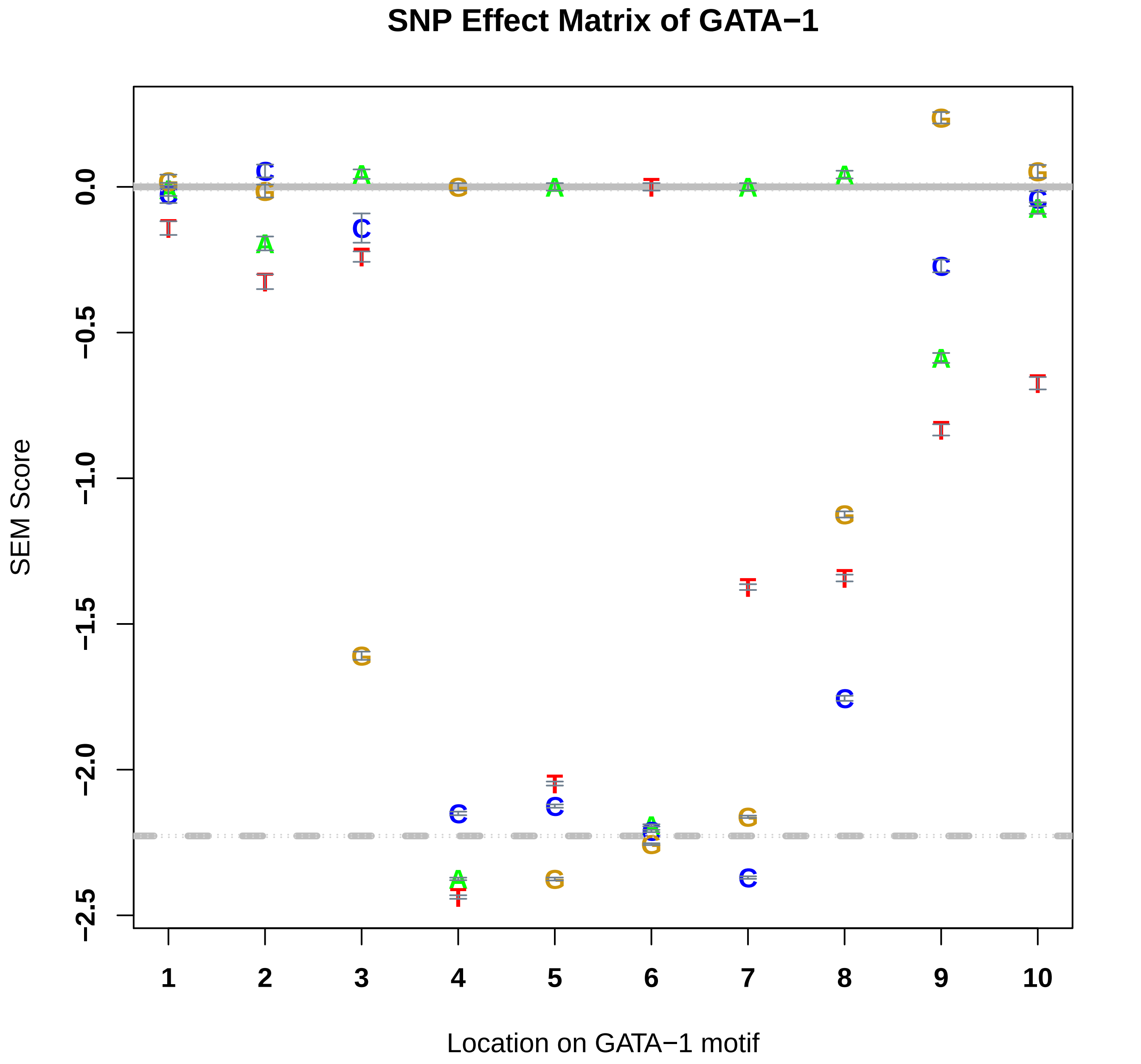

### M00348

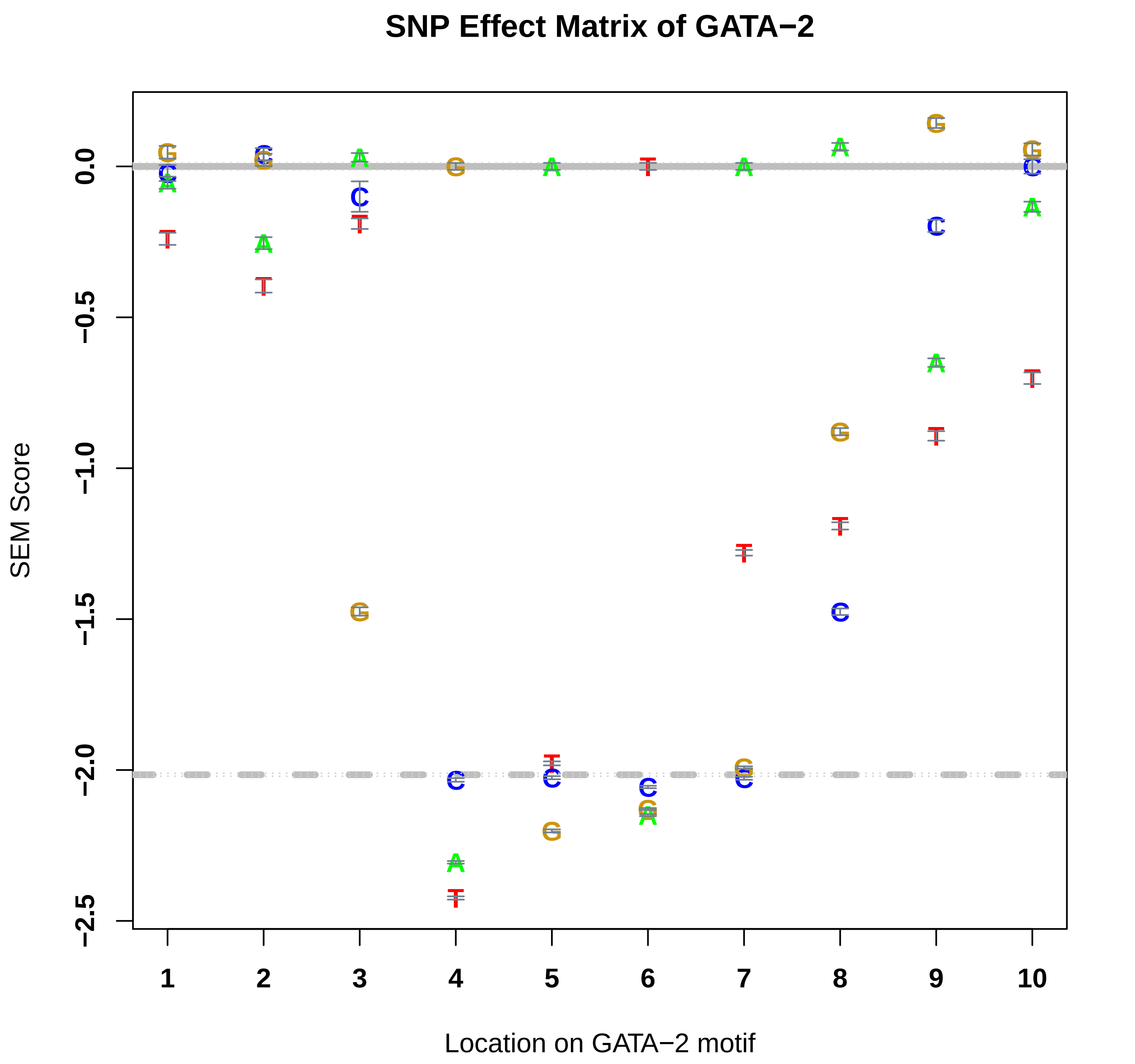

### M00349

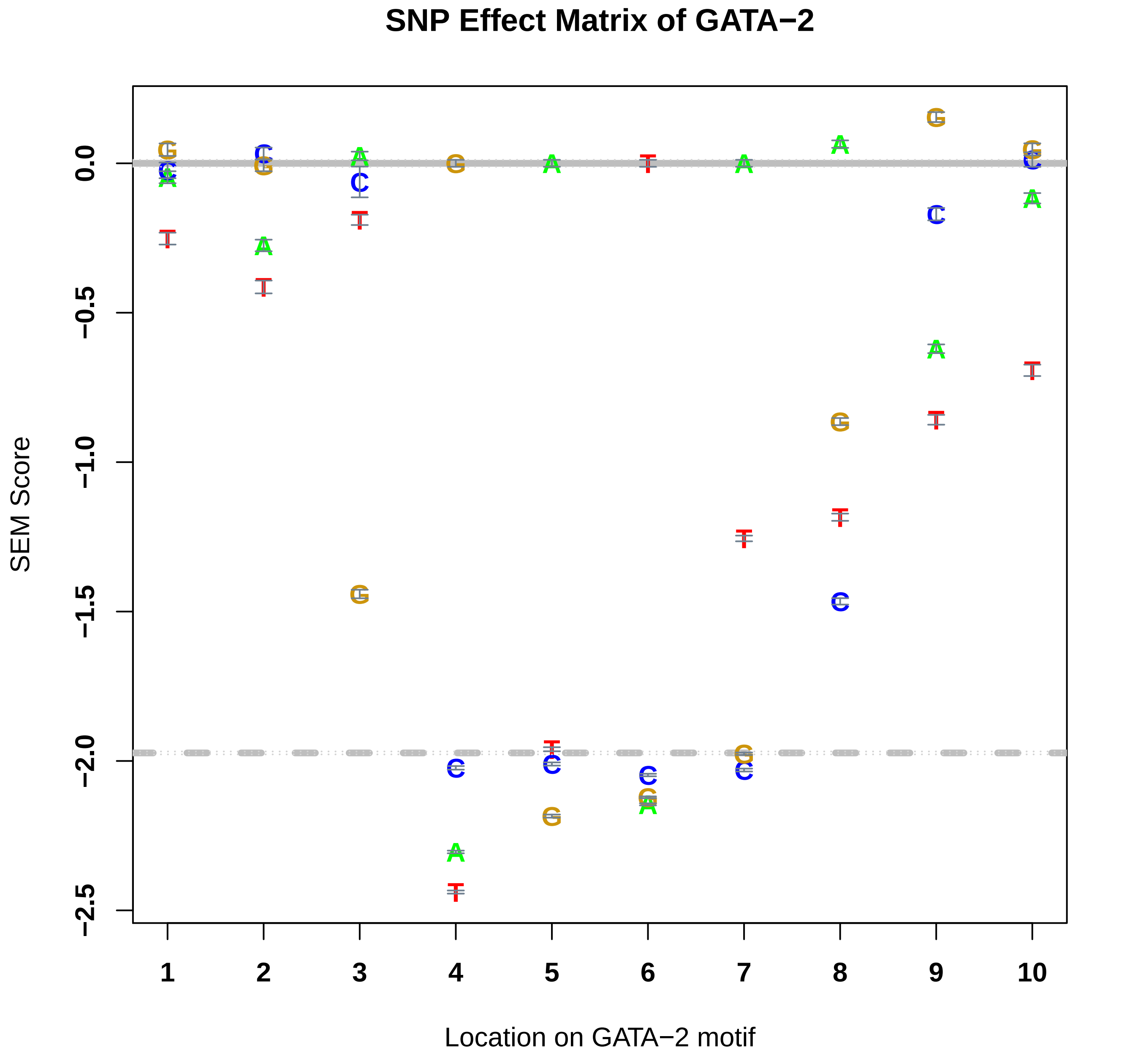

### M00350

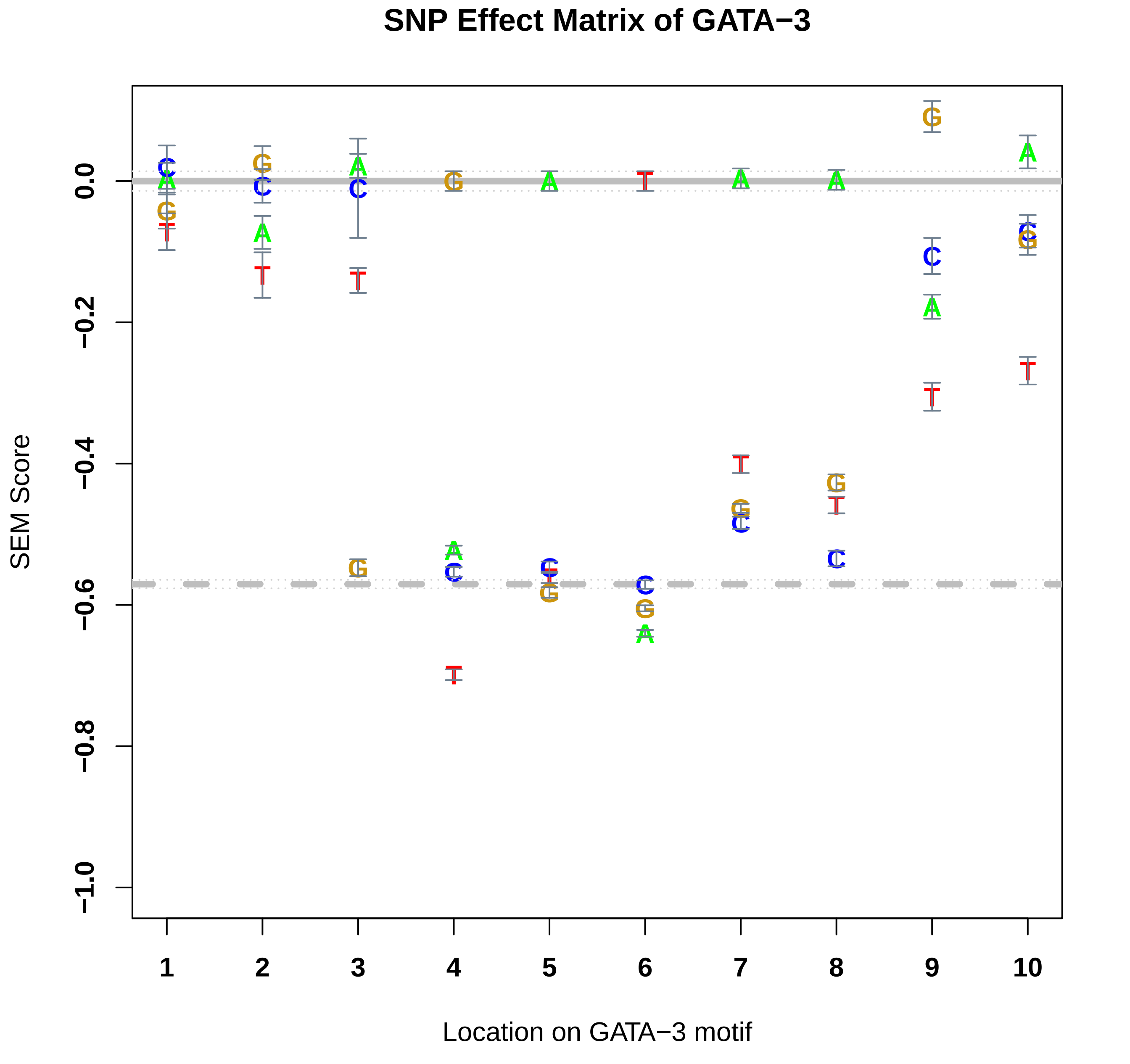

### M00351

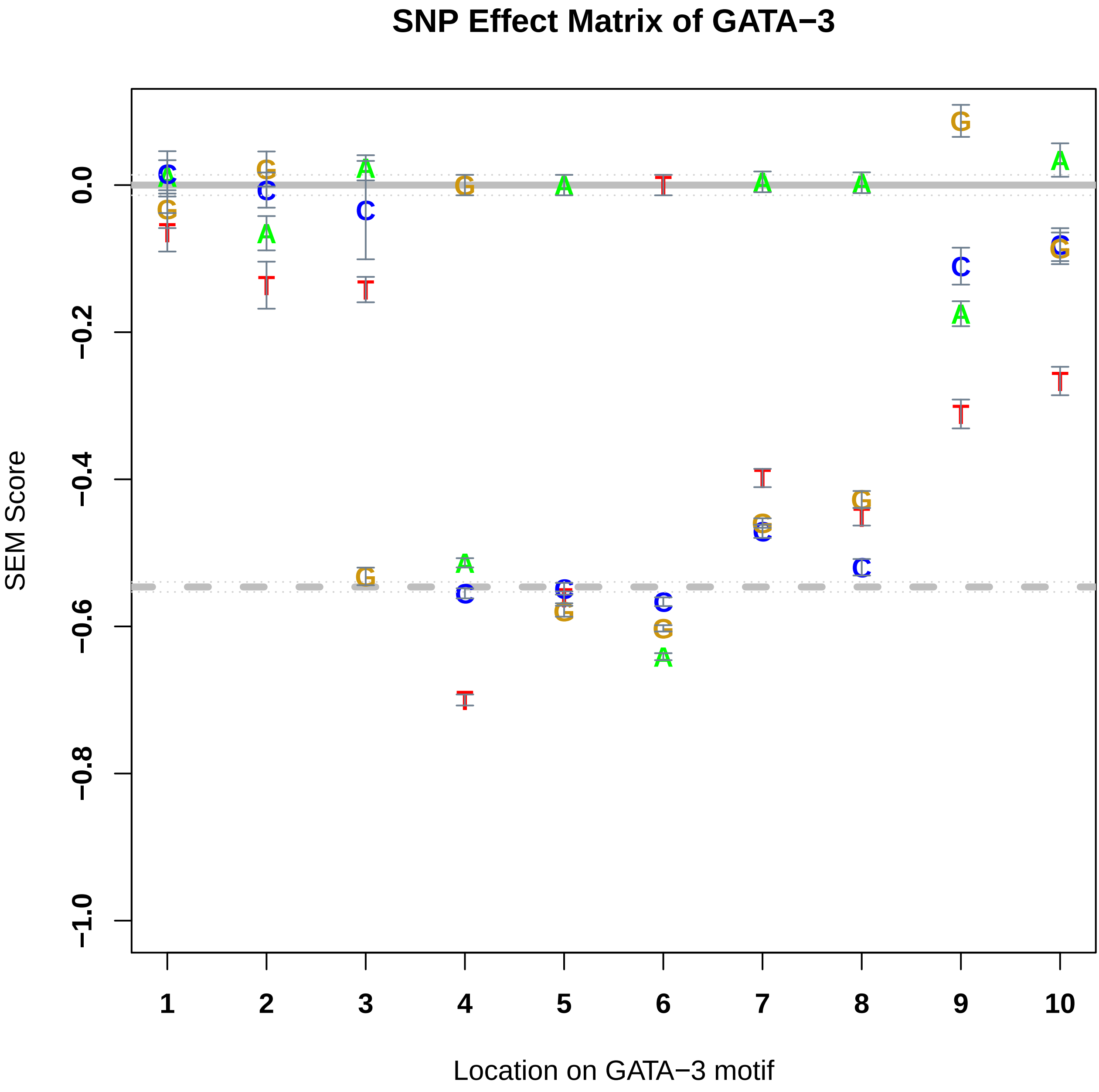

### M00415

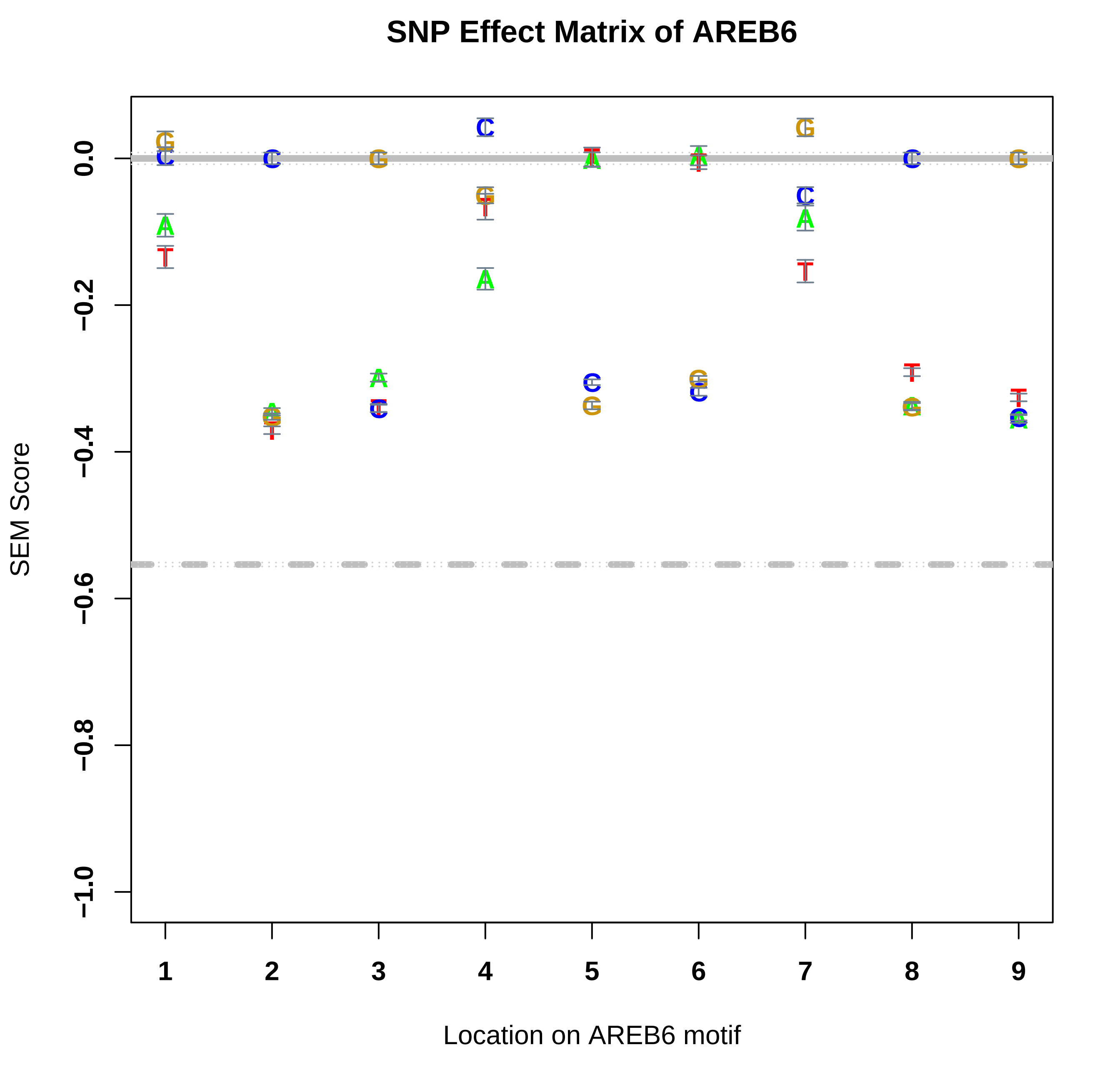

### M00418

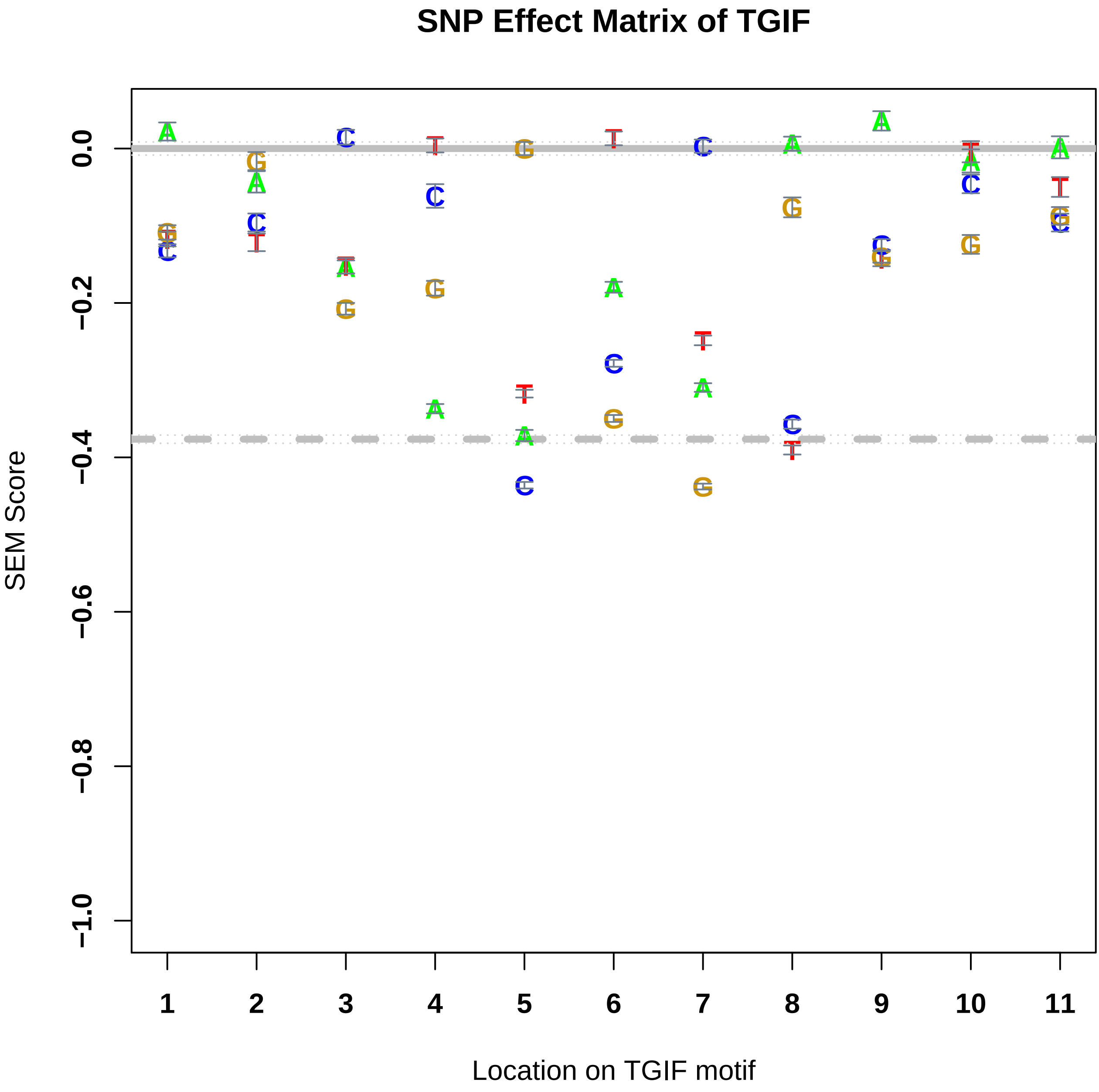

### M00427

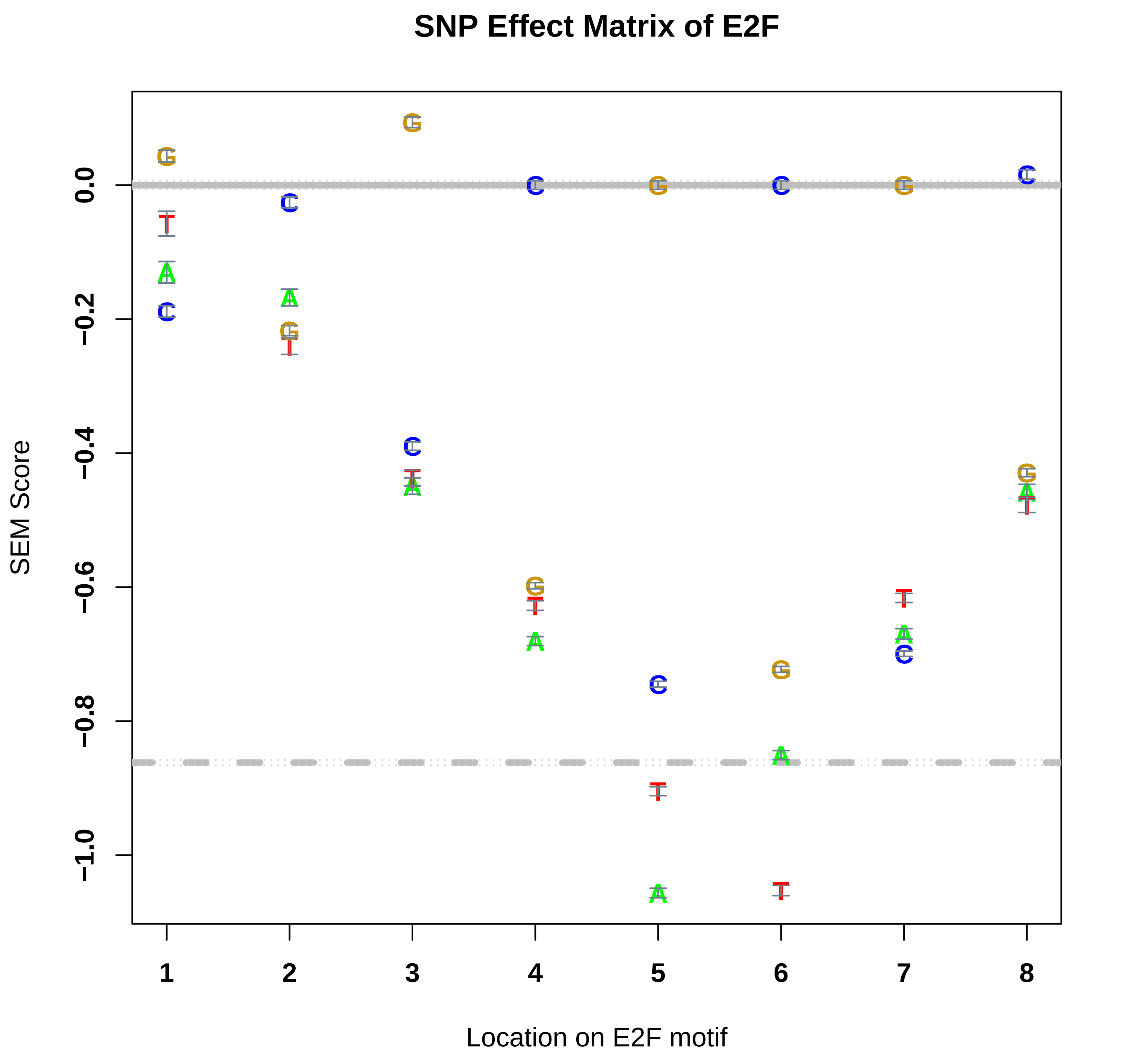

### M00428

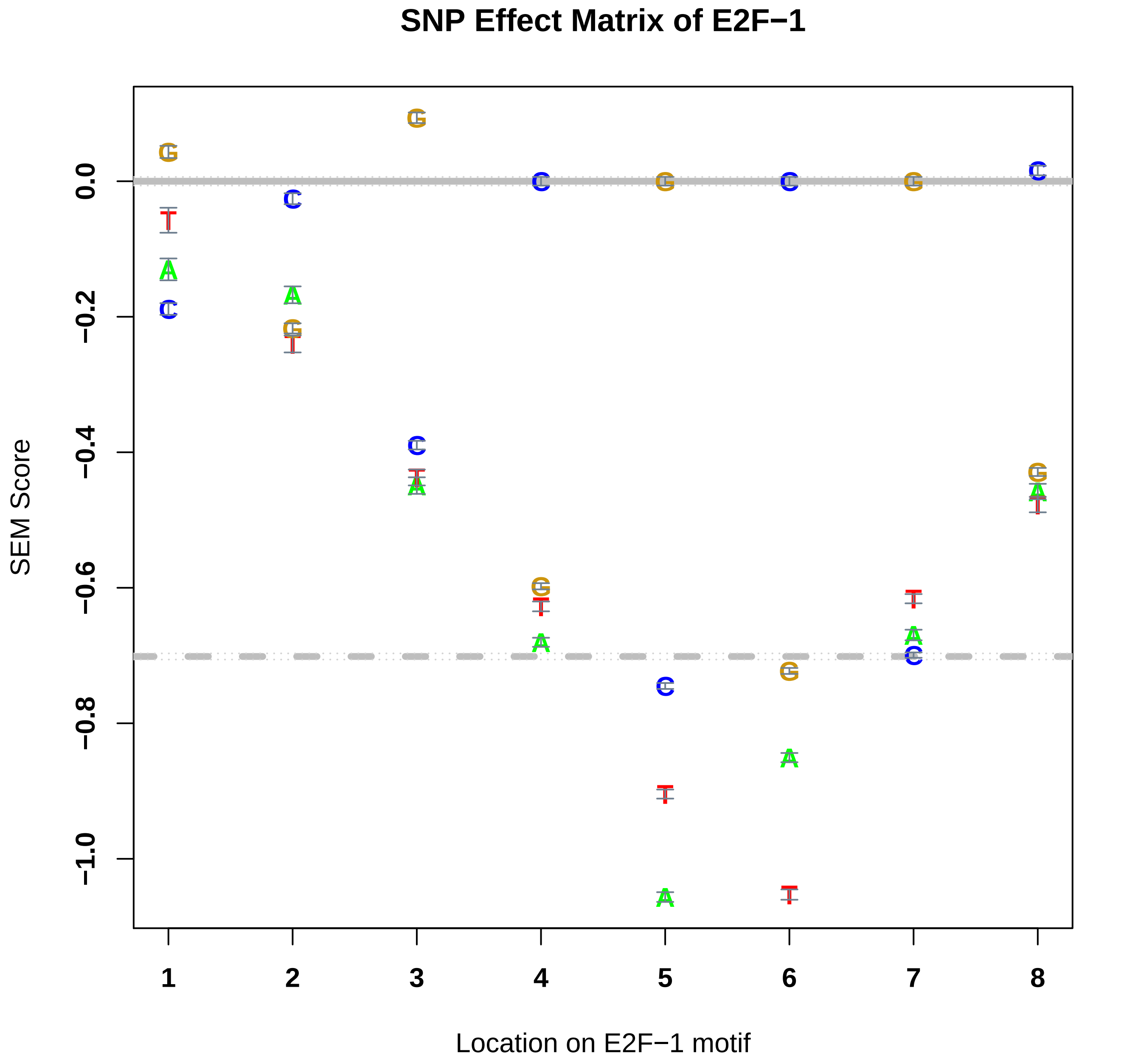

### M00430

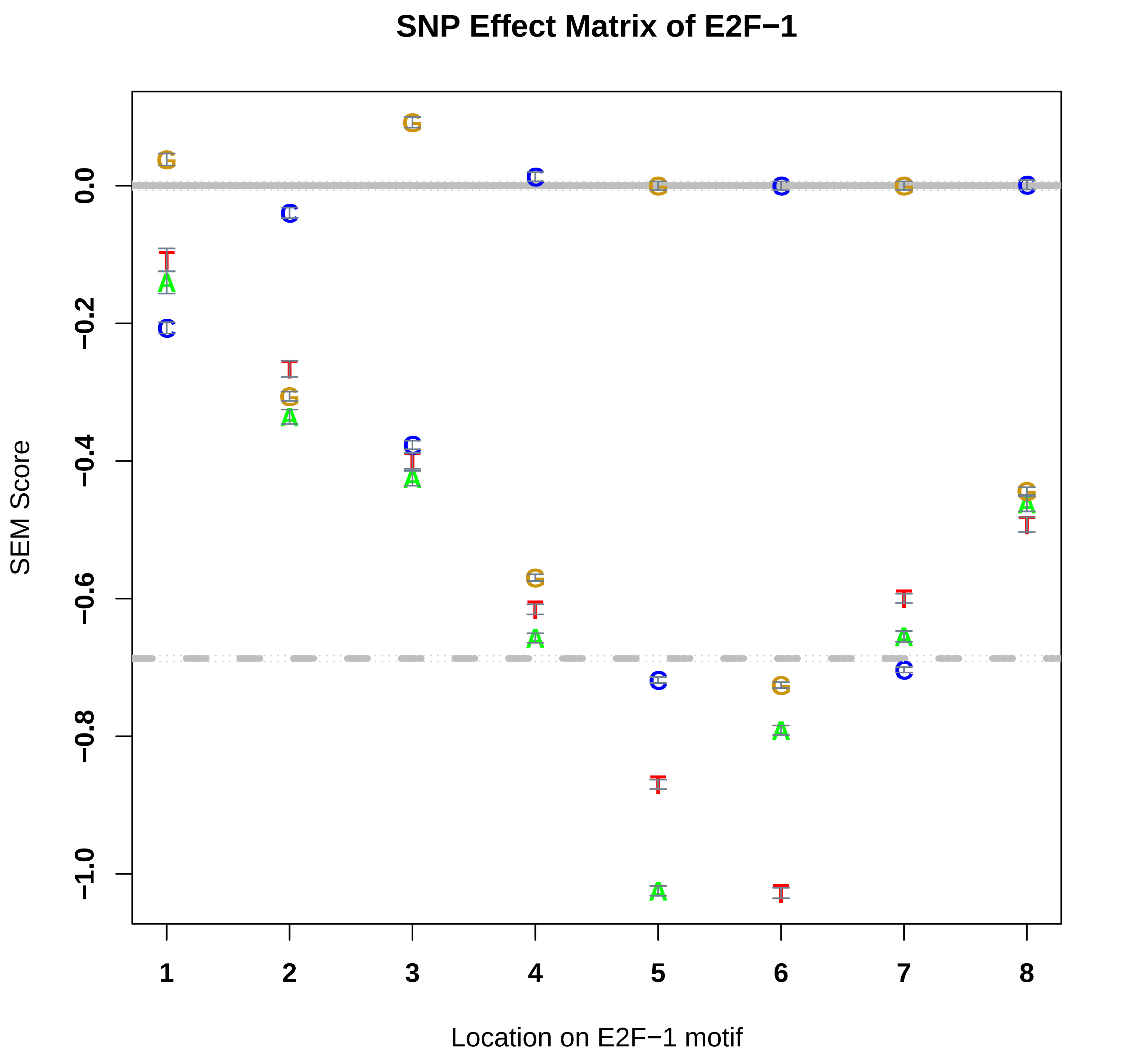

### M00431

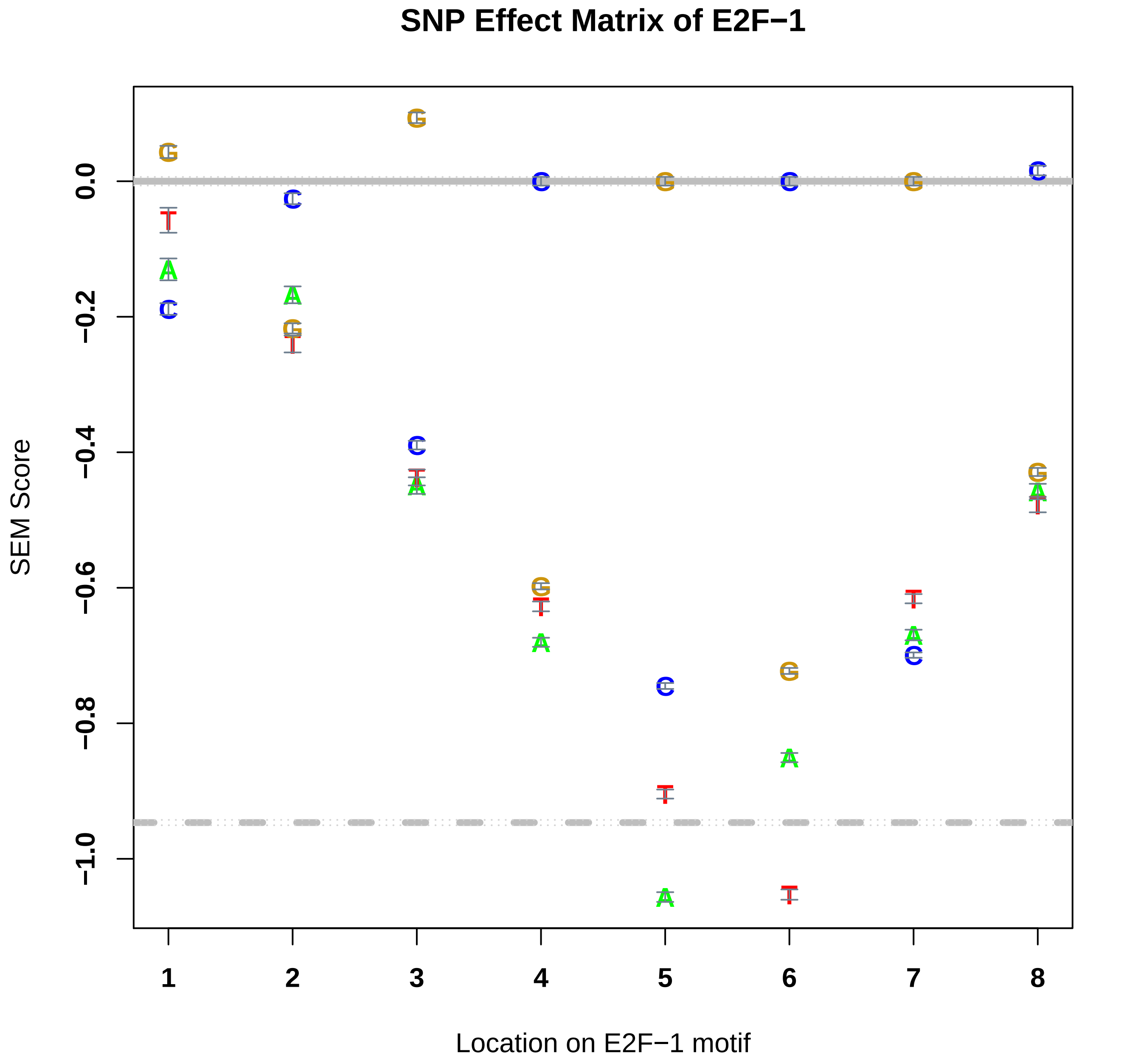

### M00471

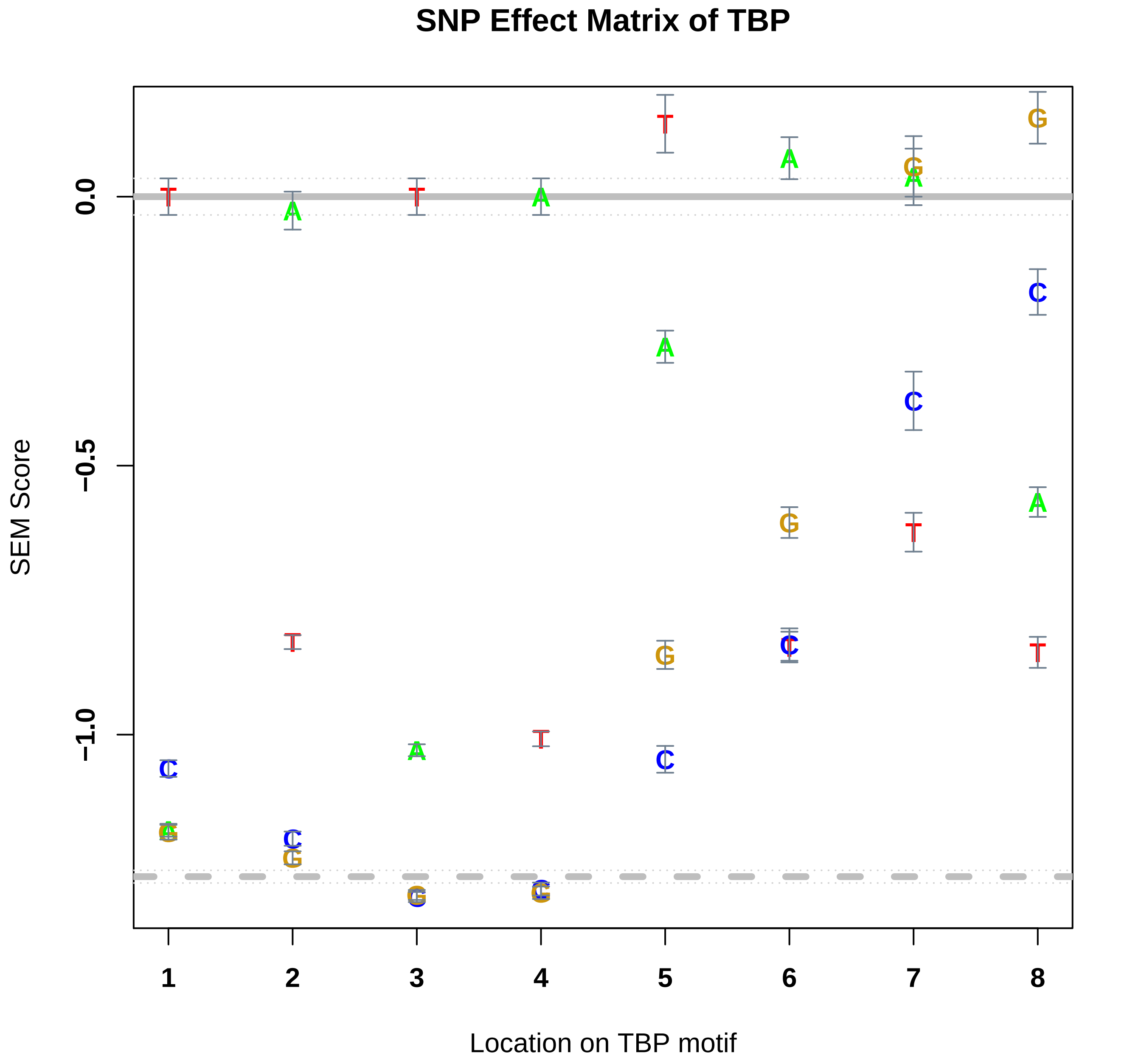

### M00491

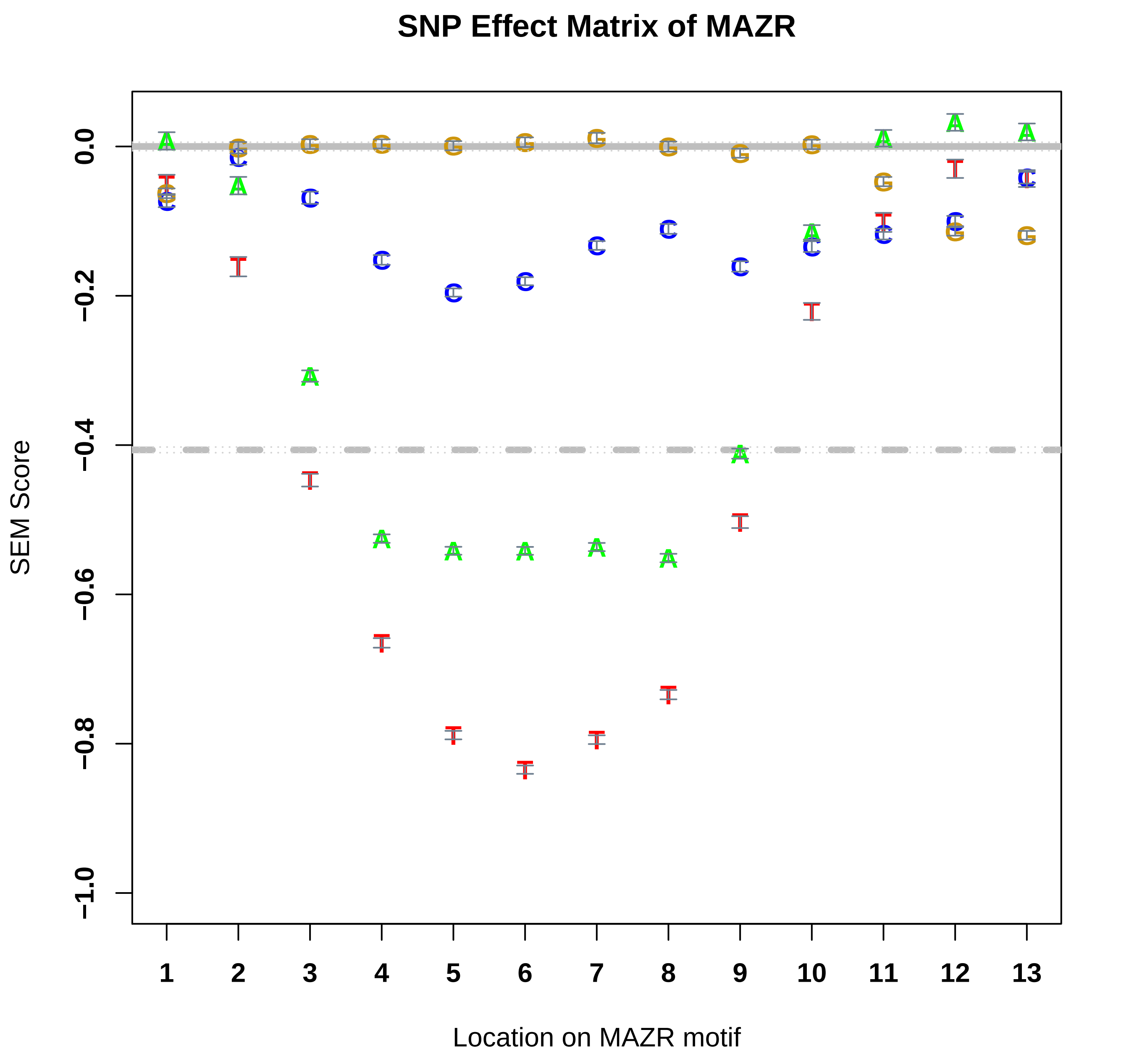

### M00492

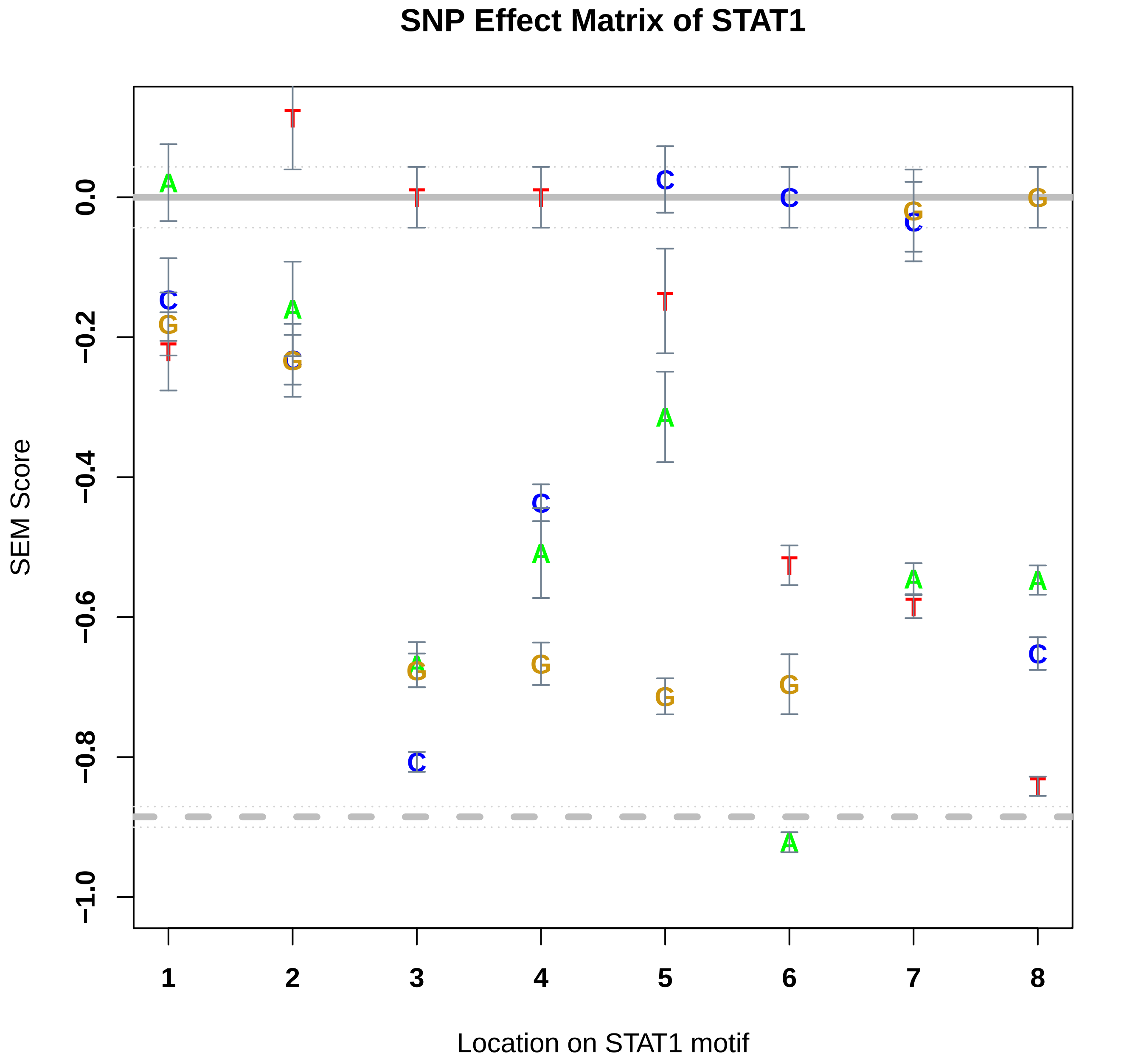

### M00493

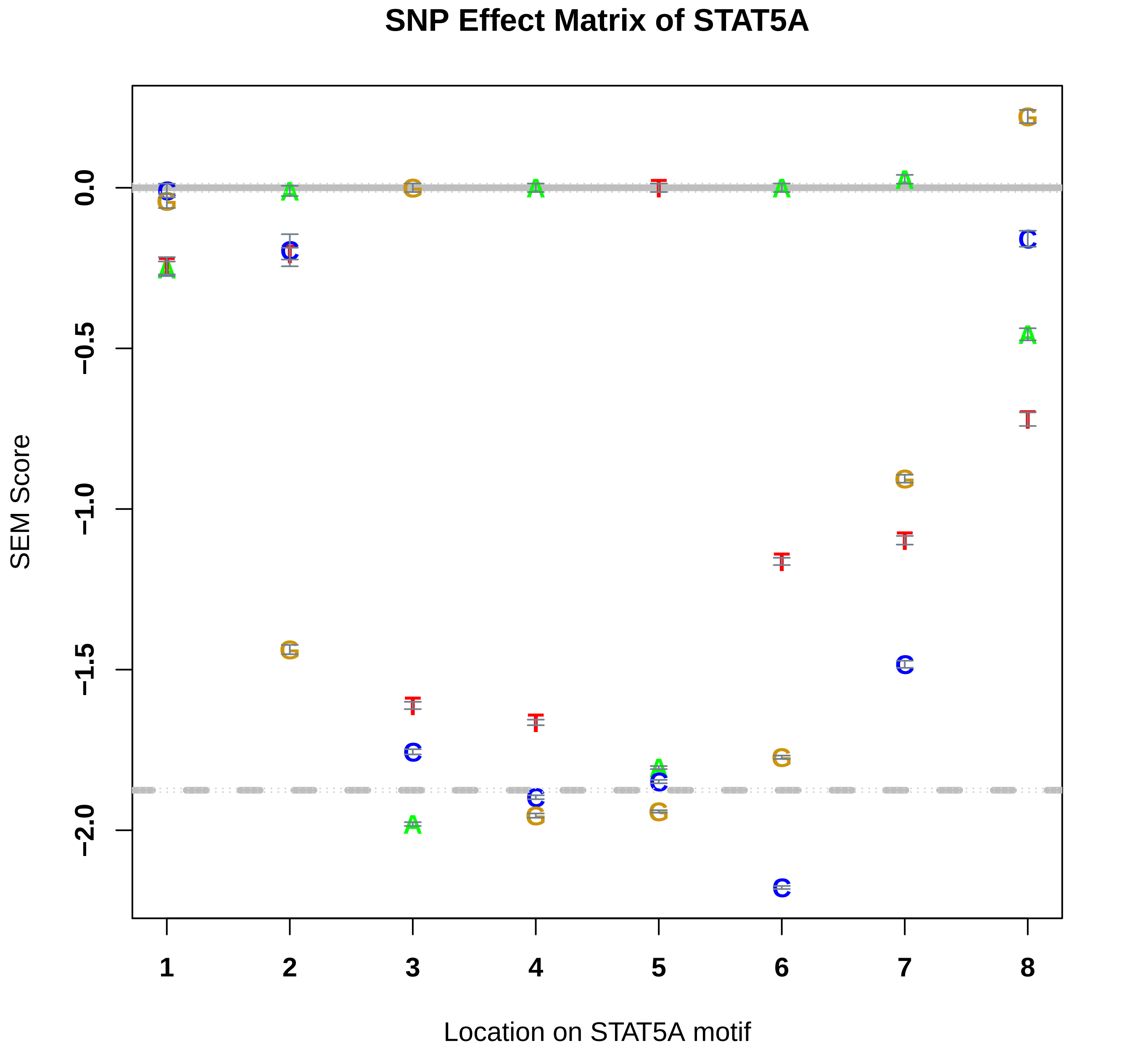

### M00496

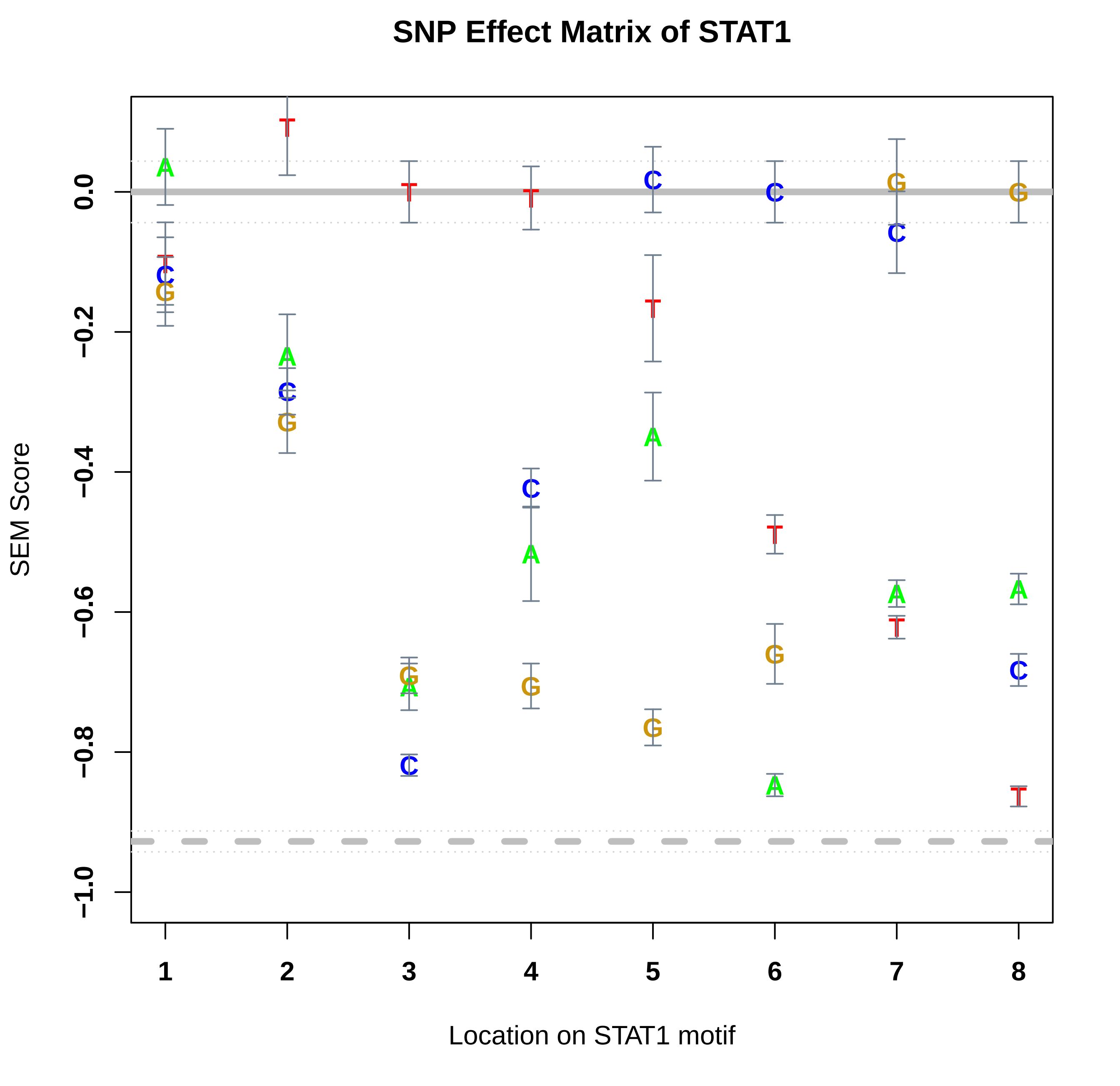

### M00497

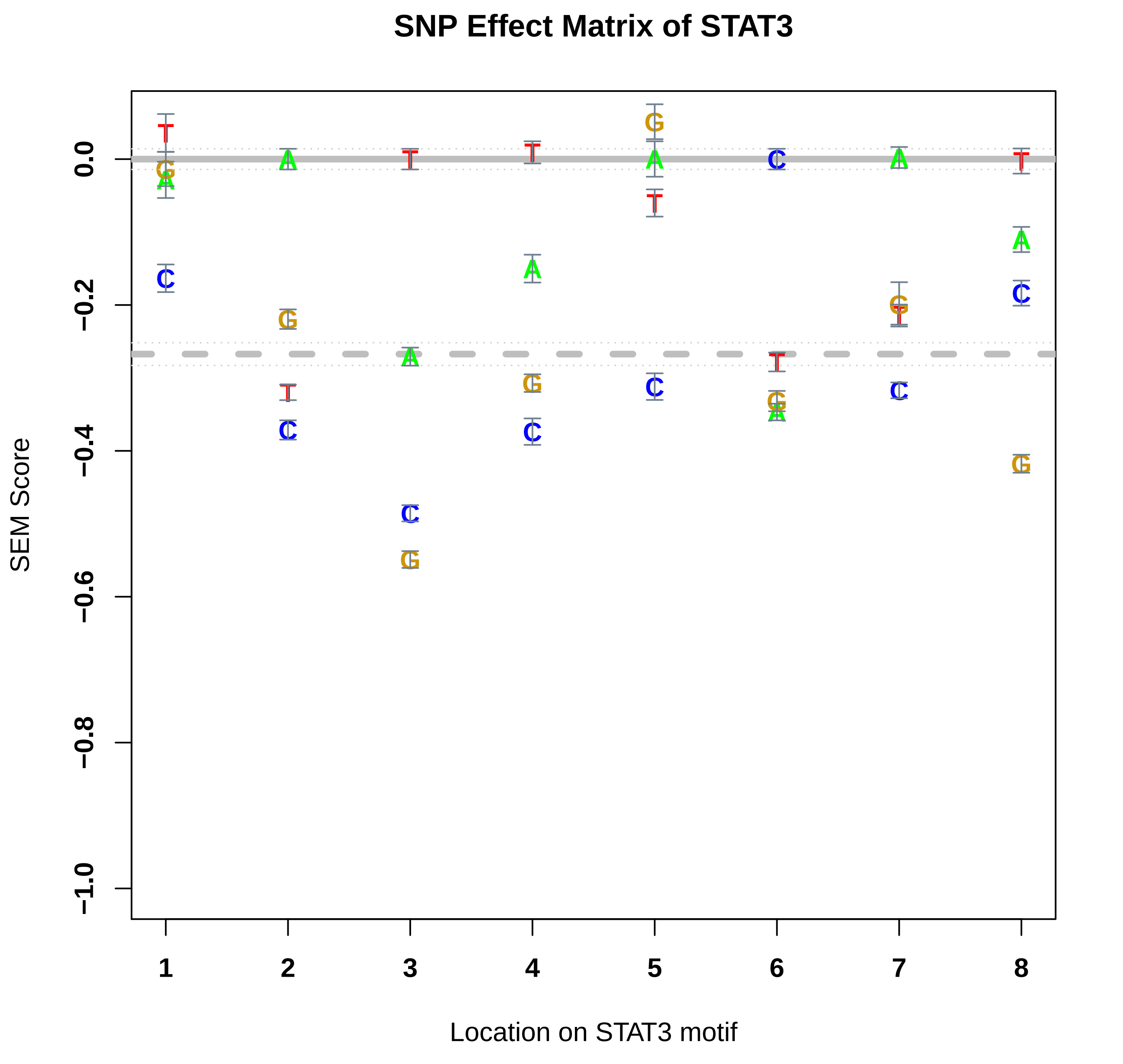

### M00499

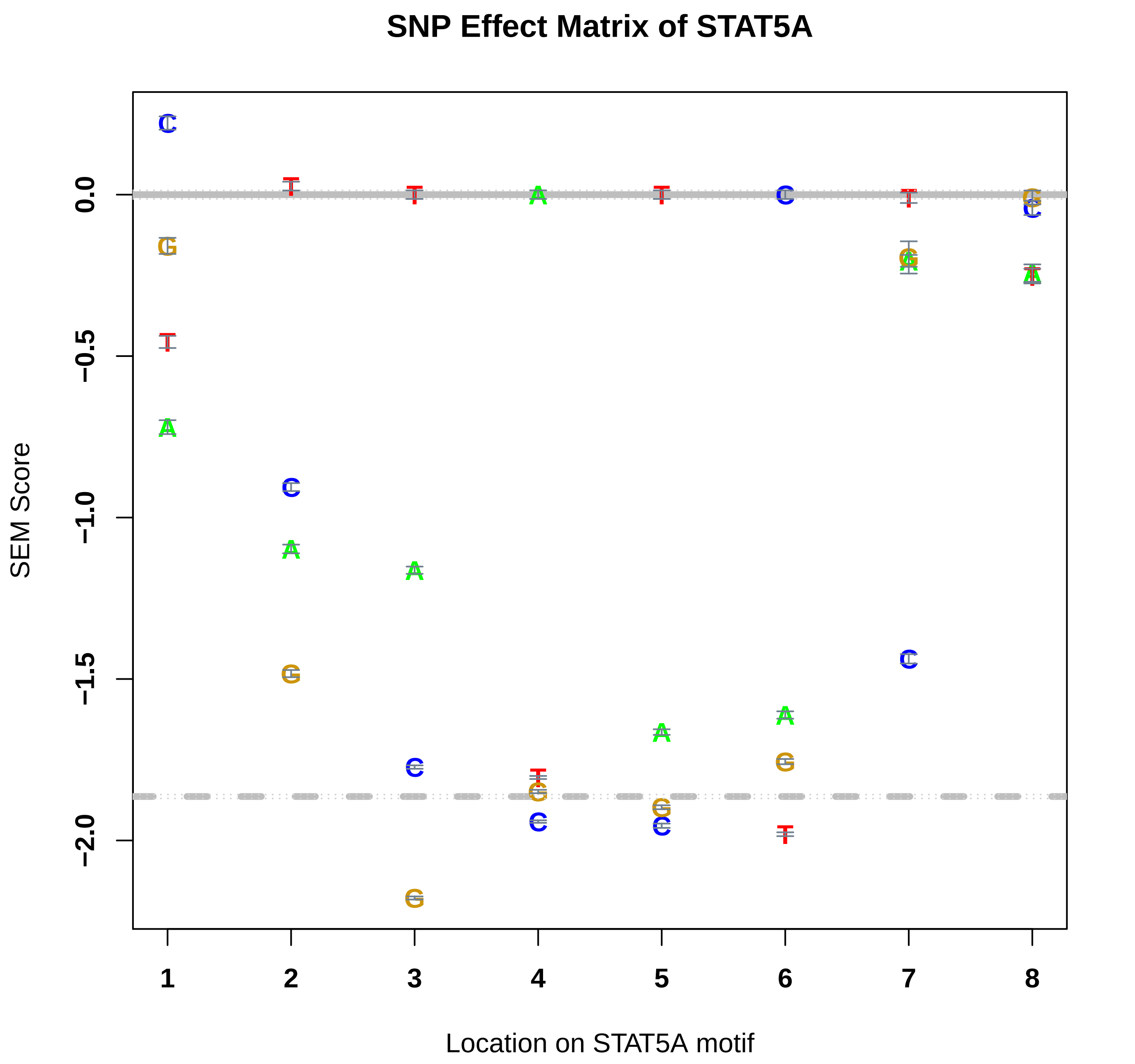

### M00499_GM12878

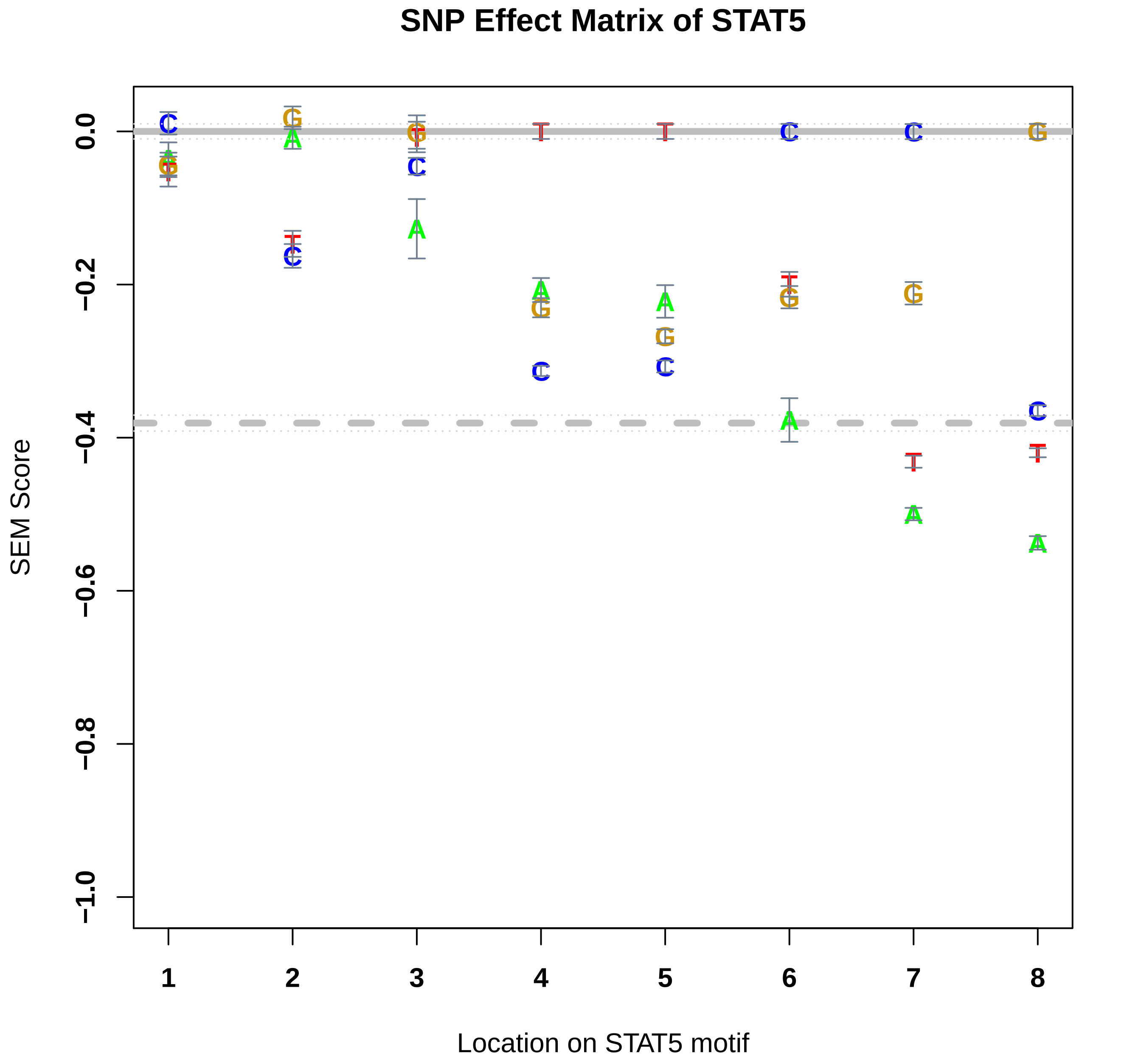

### M00499_K562

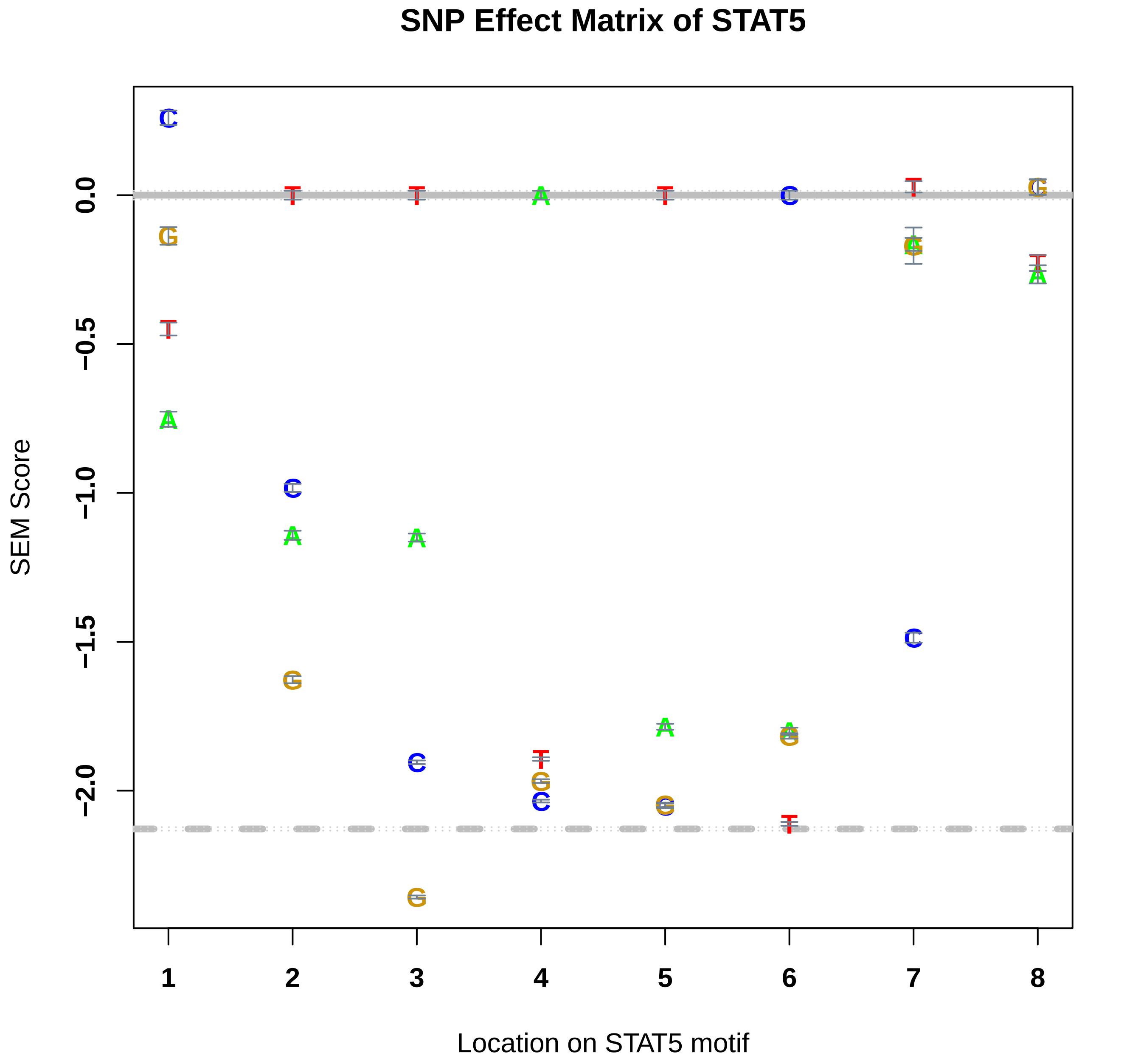

### M00652

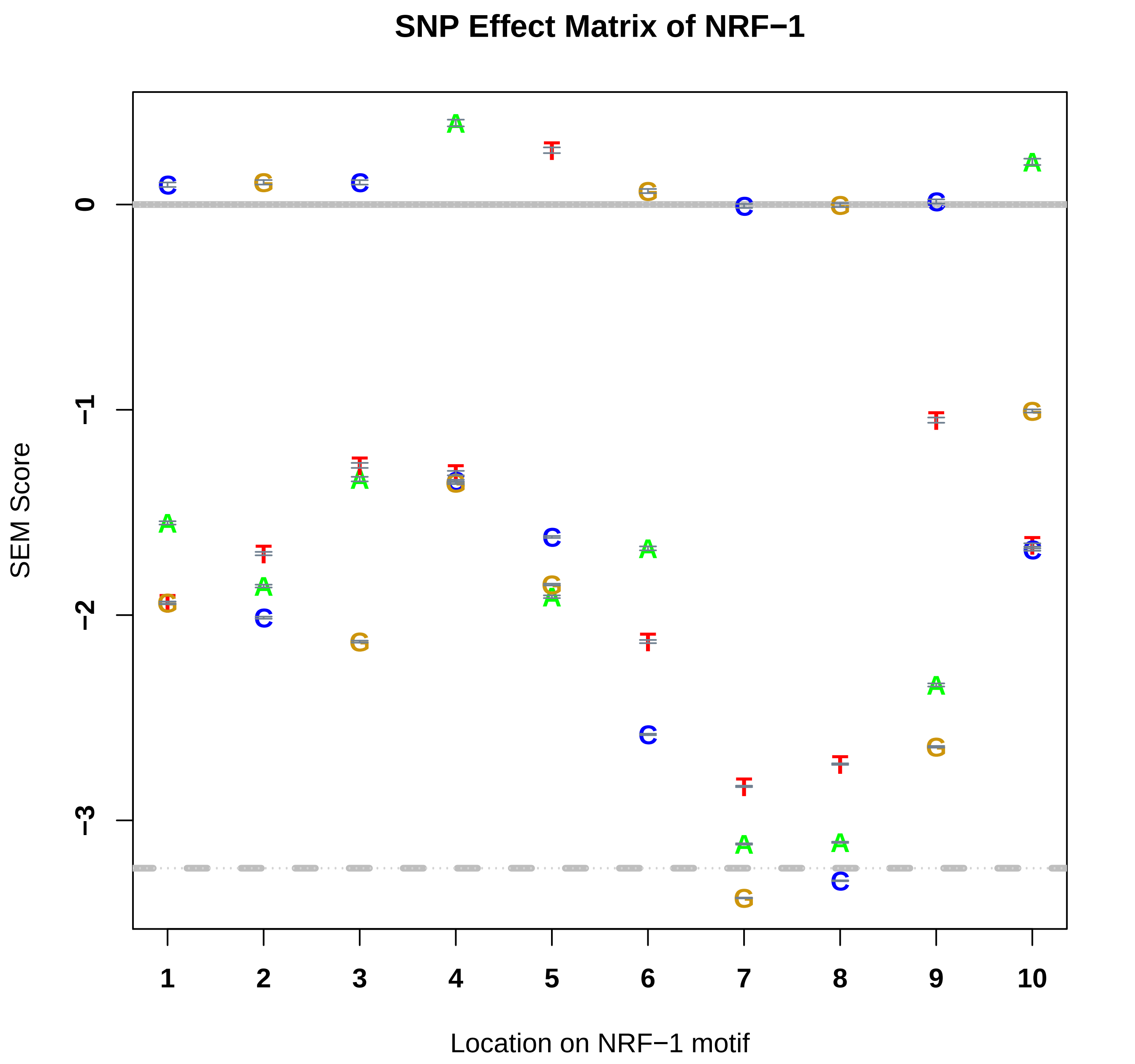

### M00698

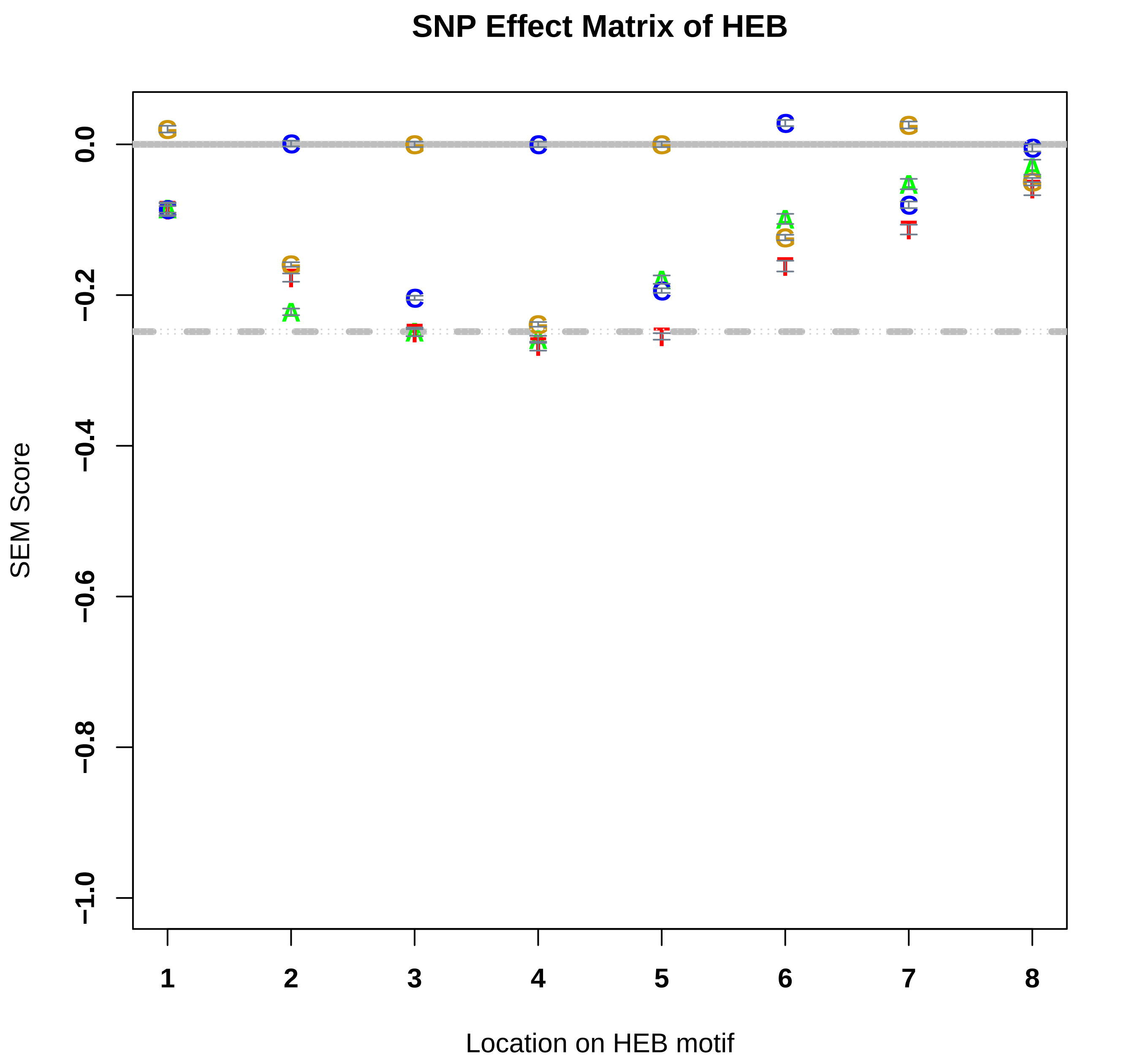

### M00724

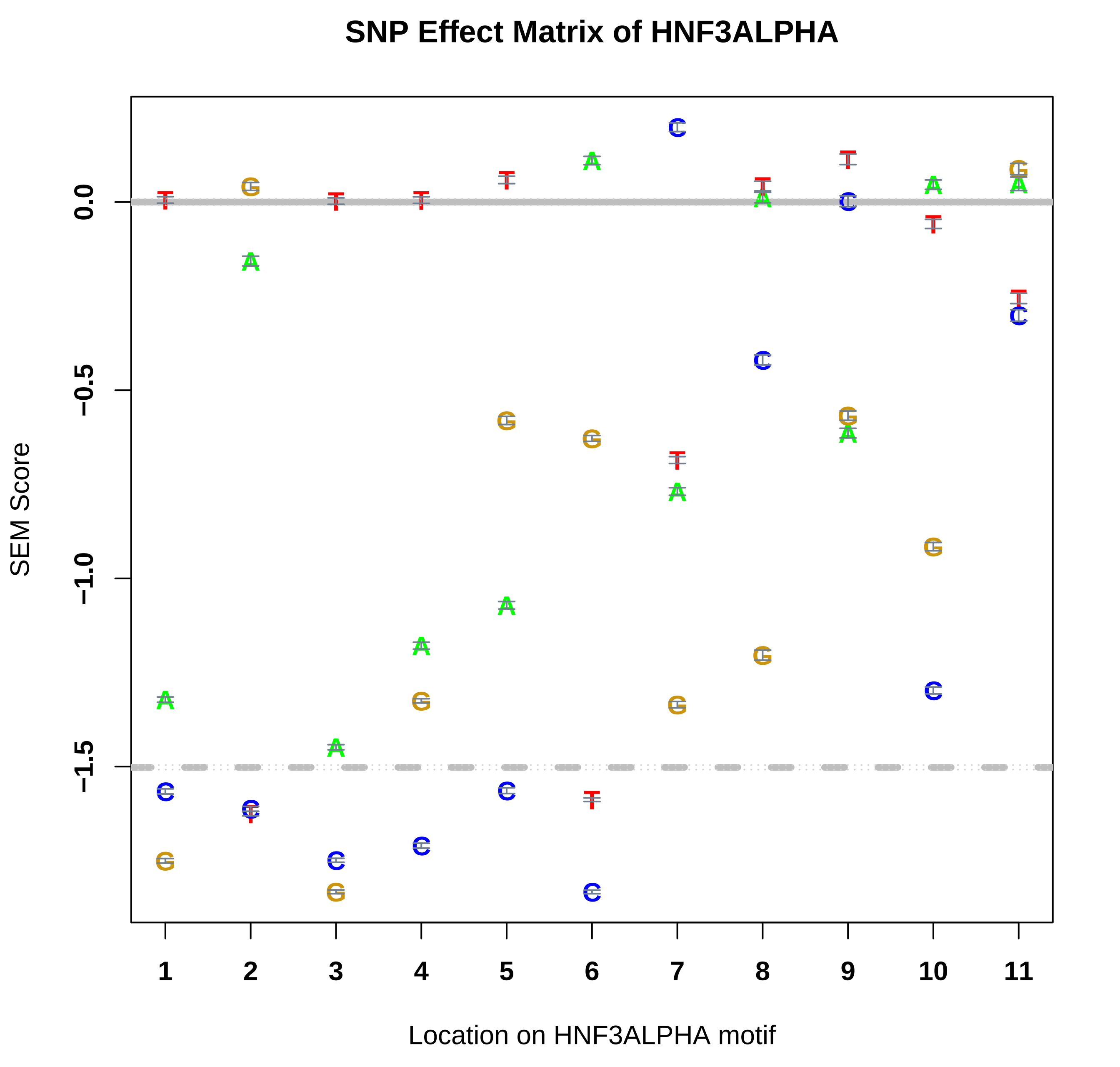

### M00727

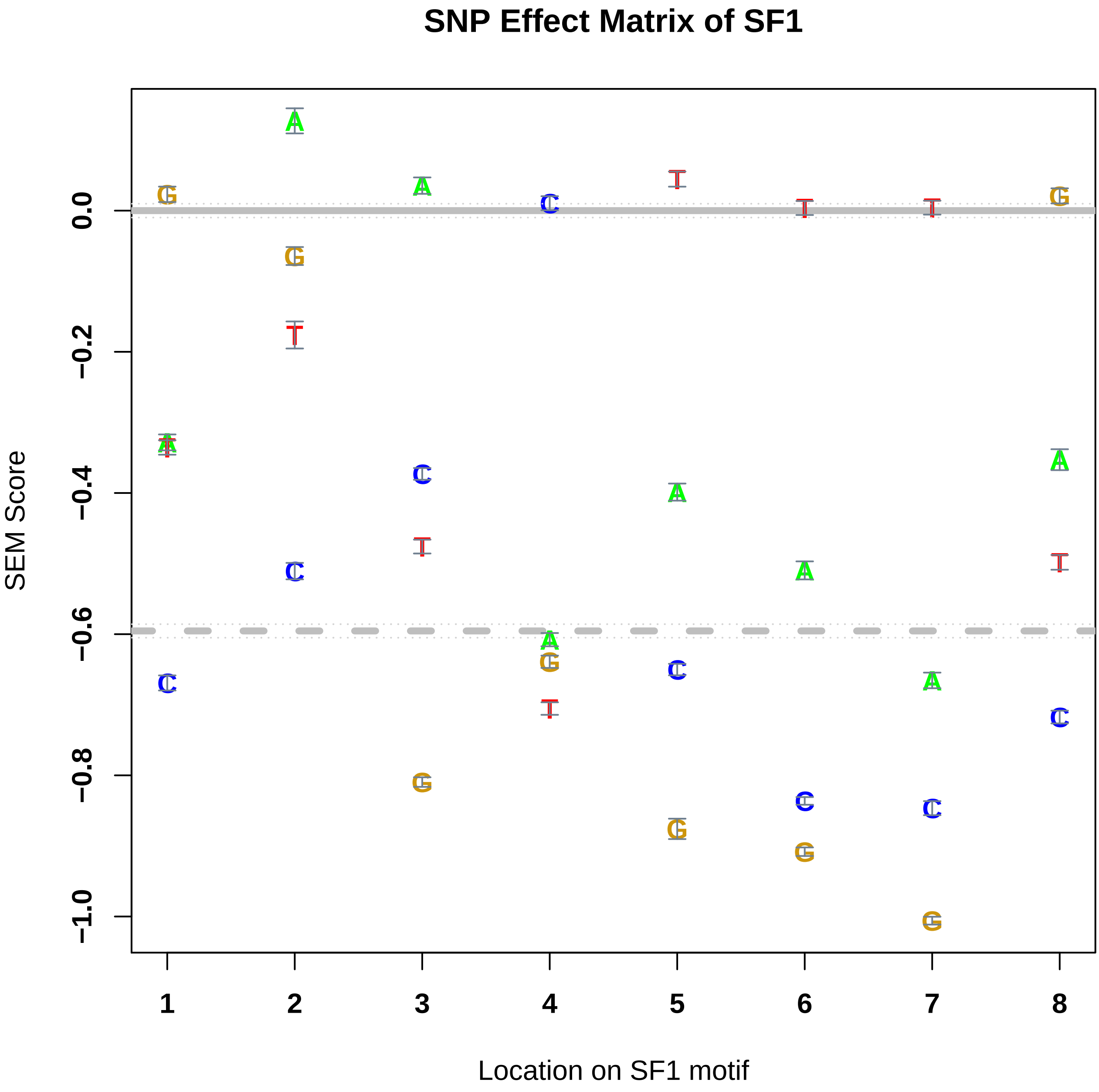

### M00736

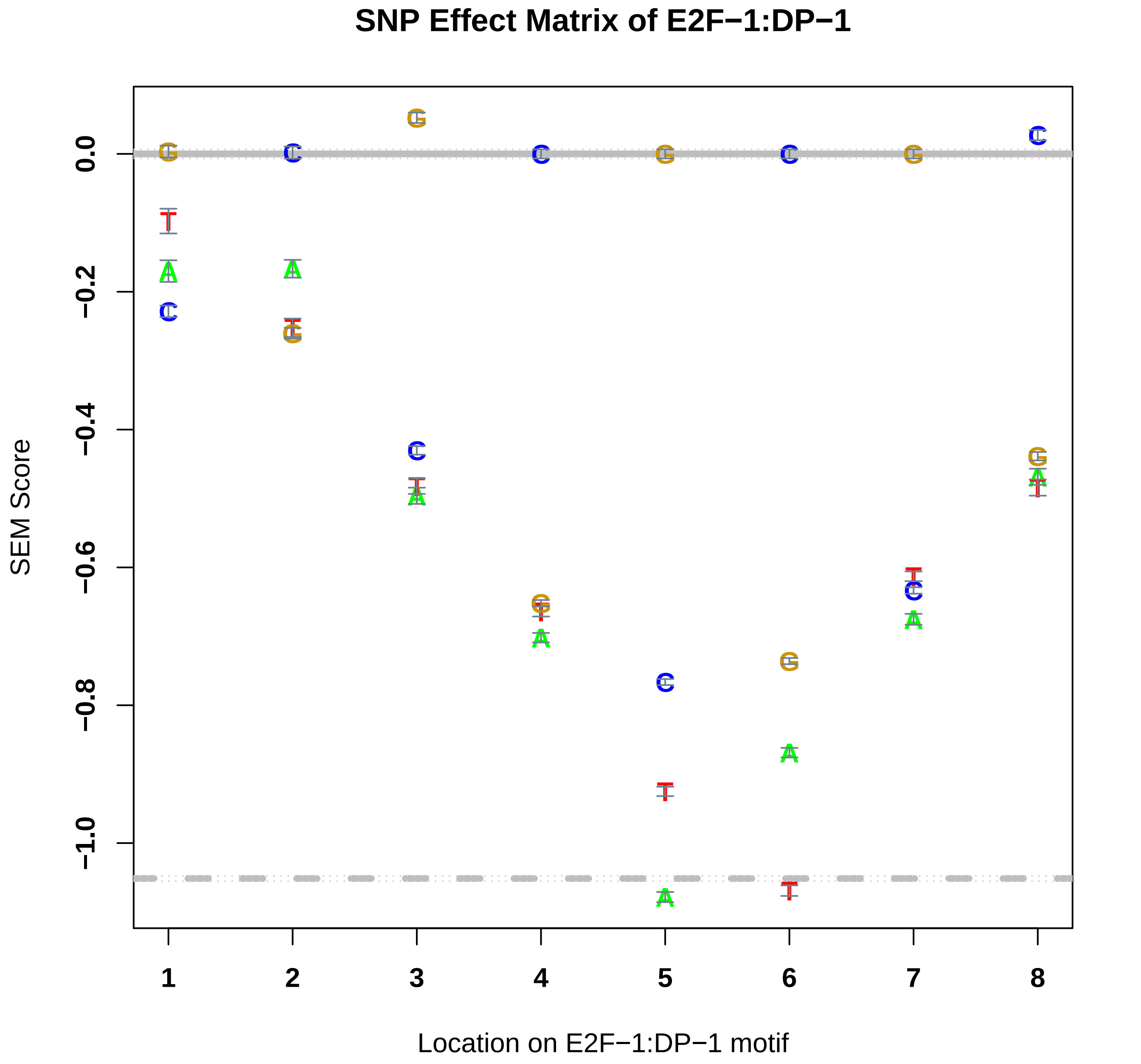

### M00737

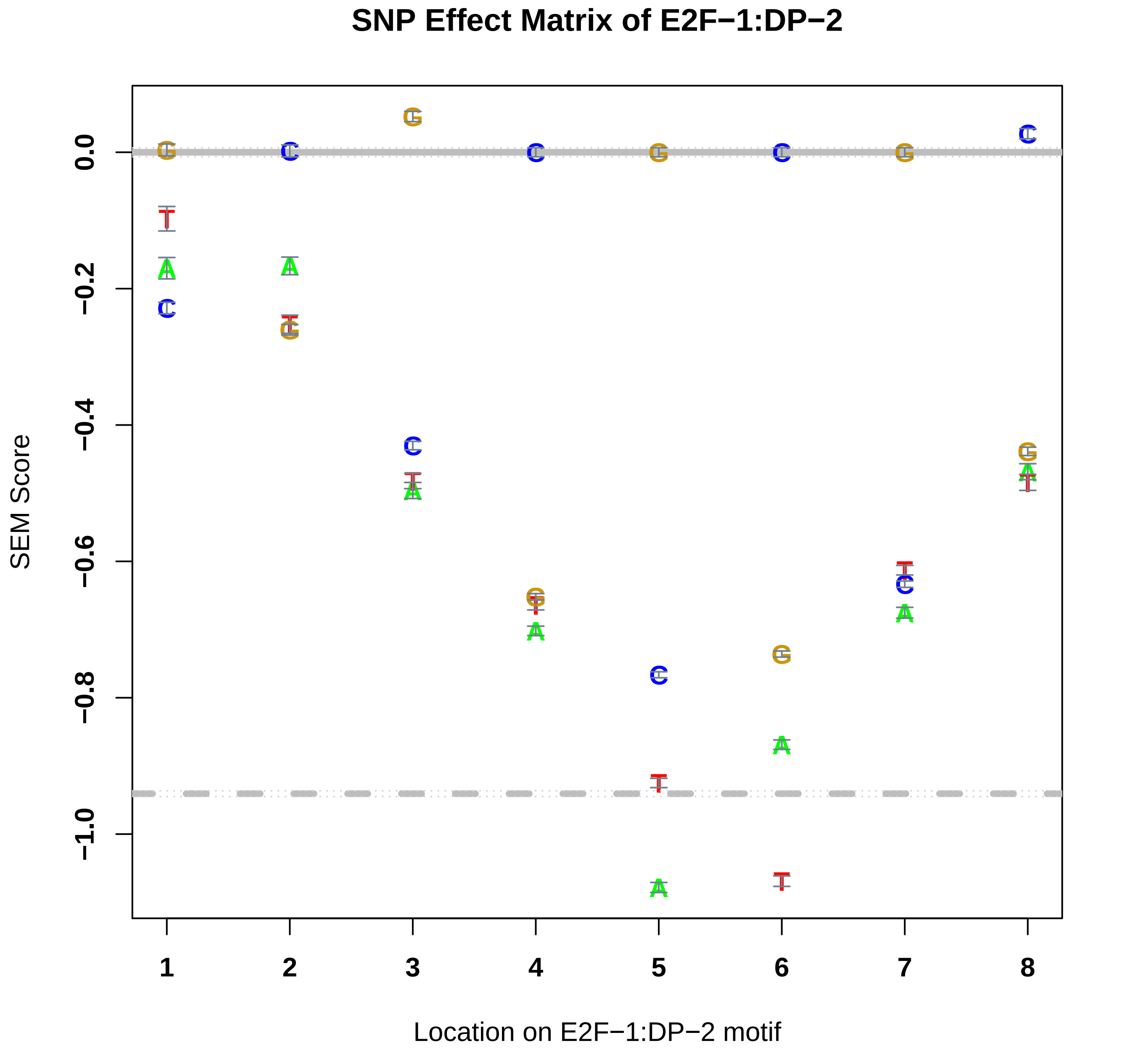

### M00738

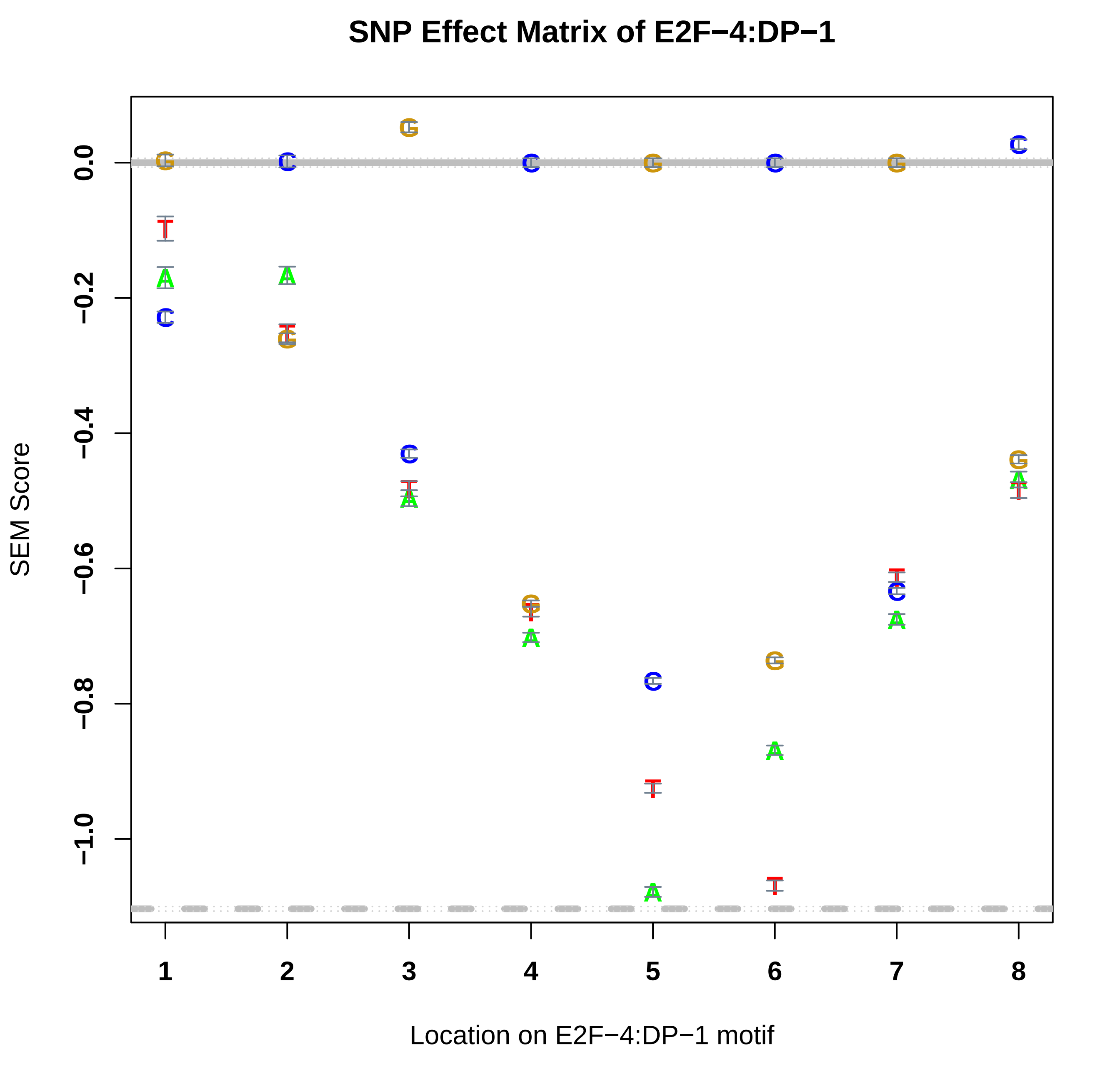

### M00739

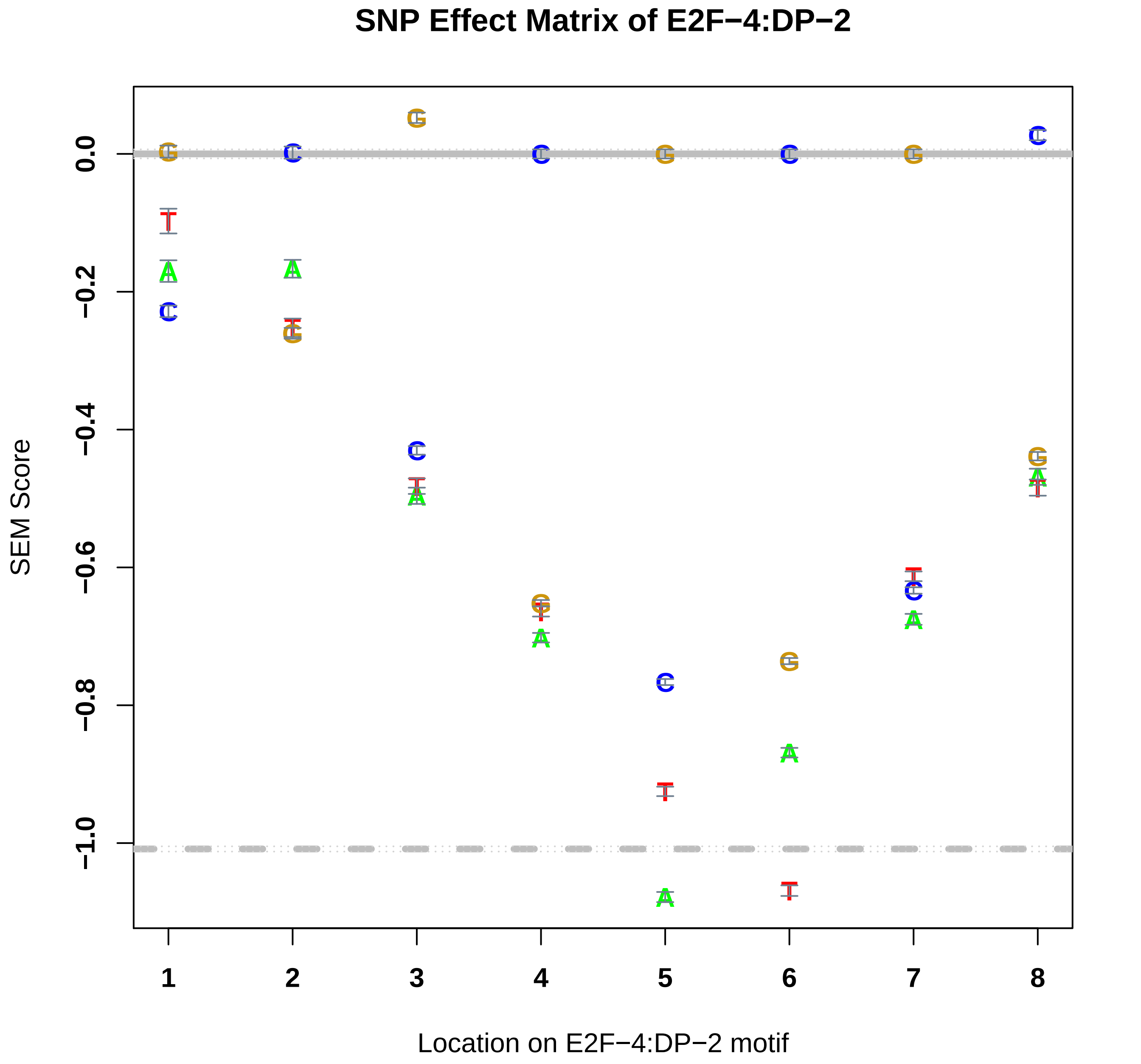

### M00740

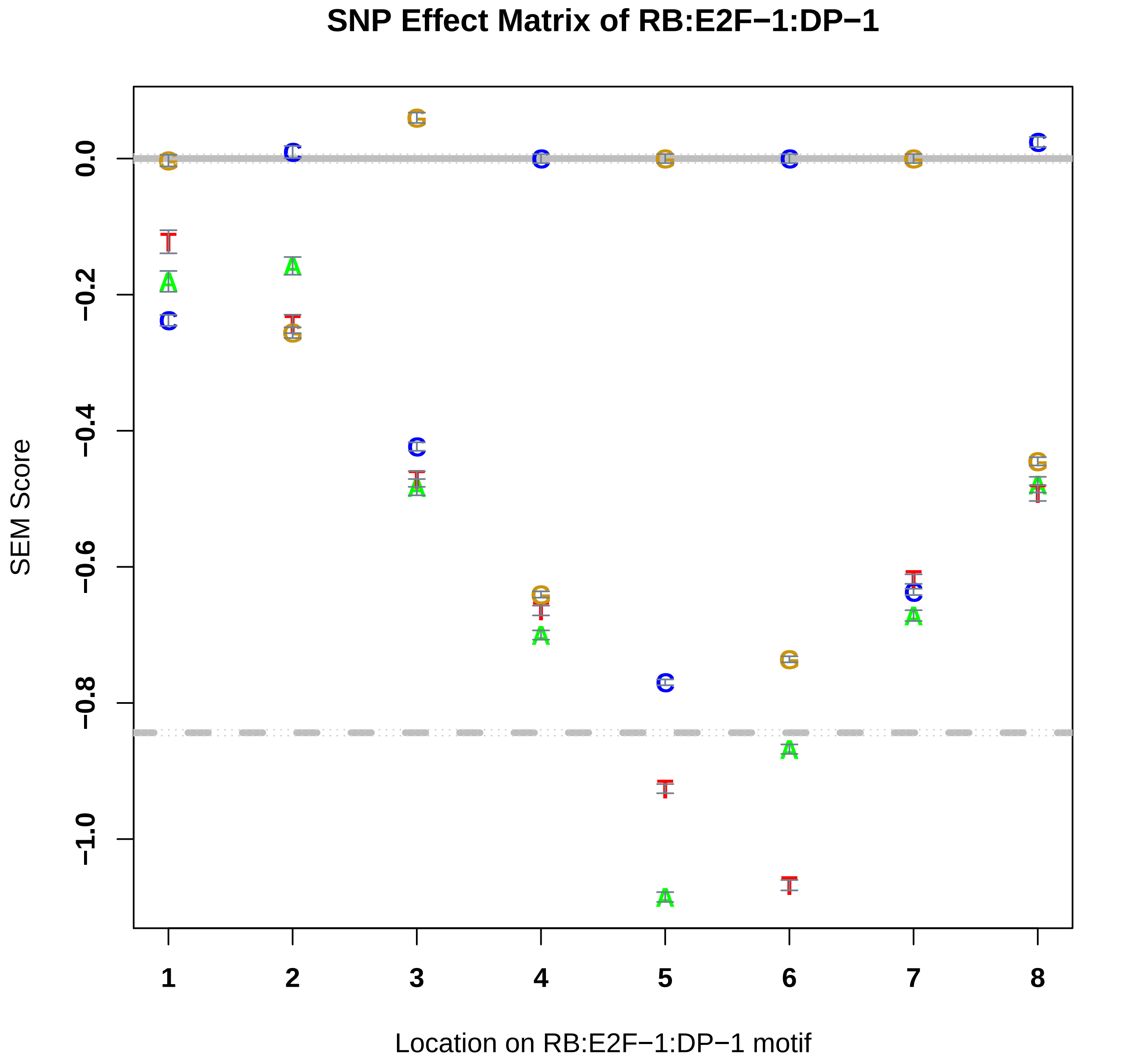

### M00746

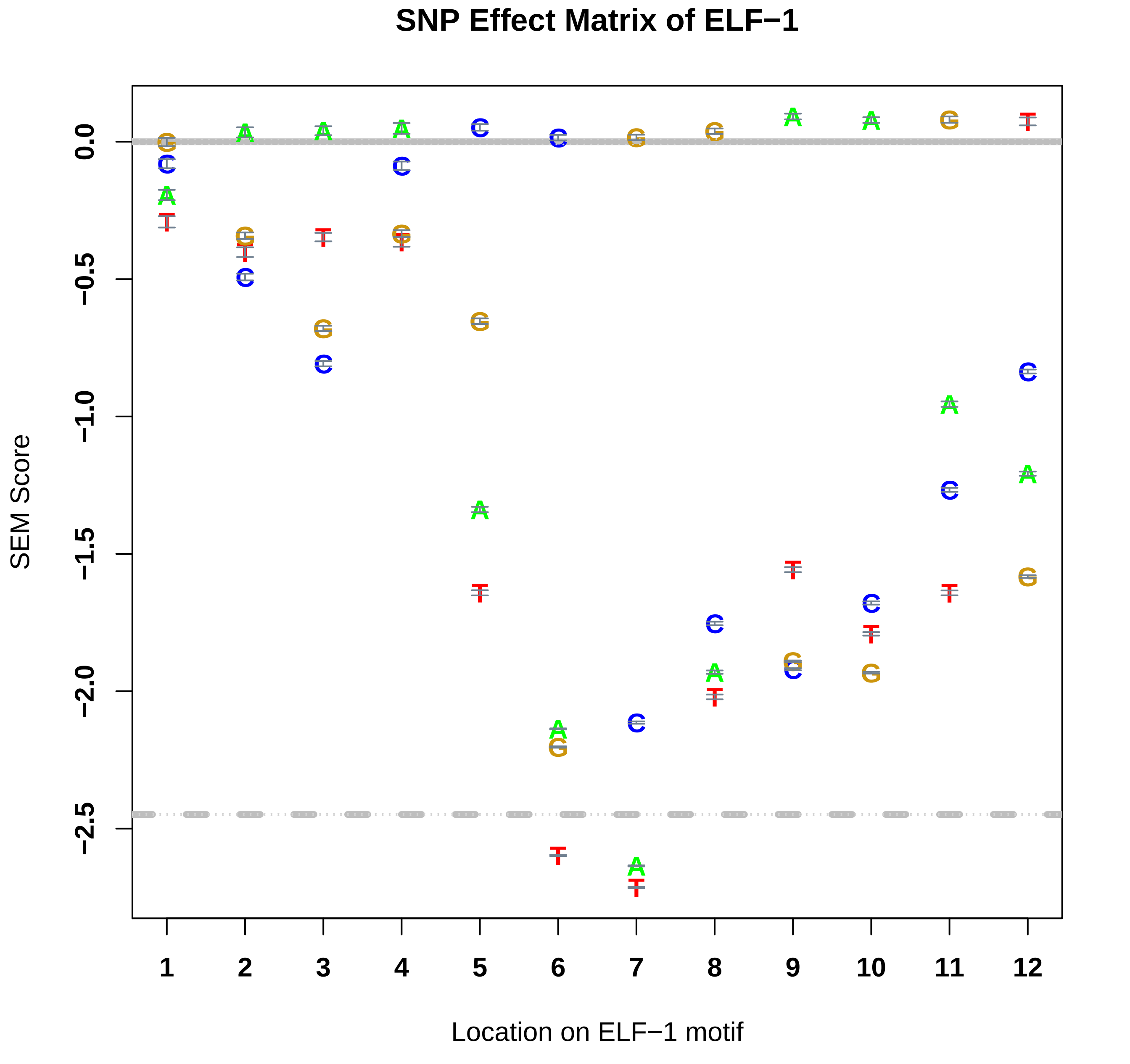

### M00749

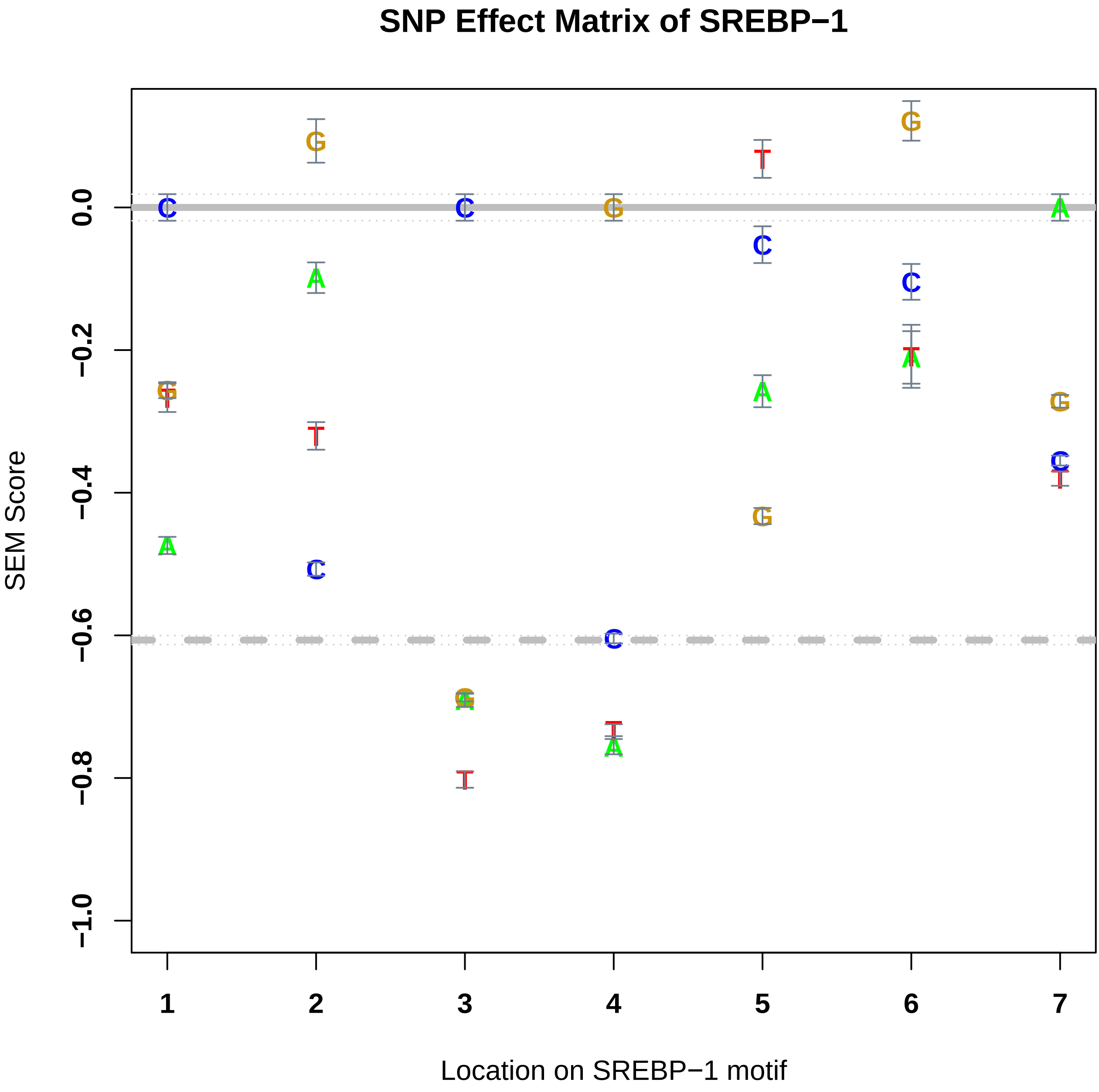

### M00751

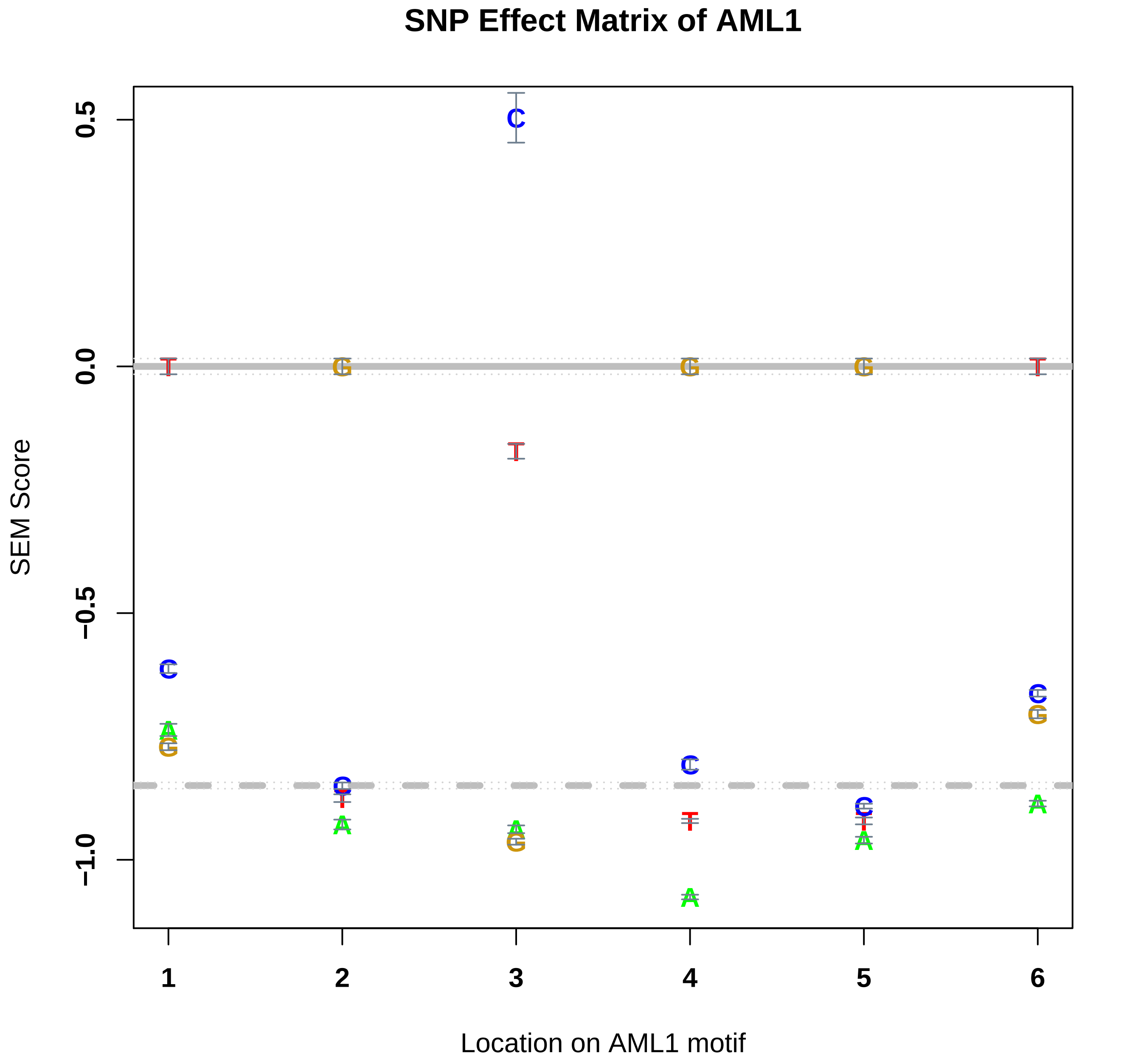

### M00773

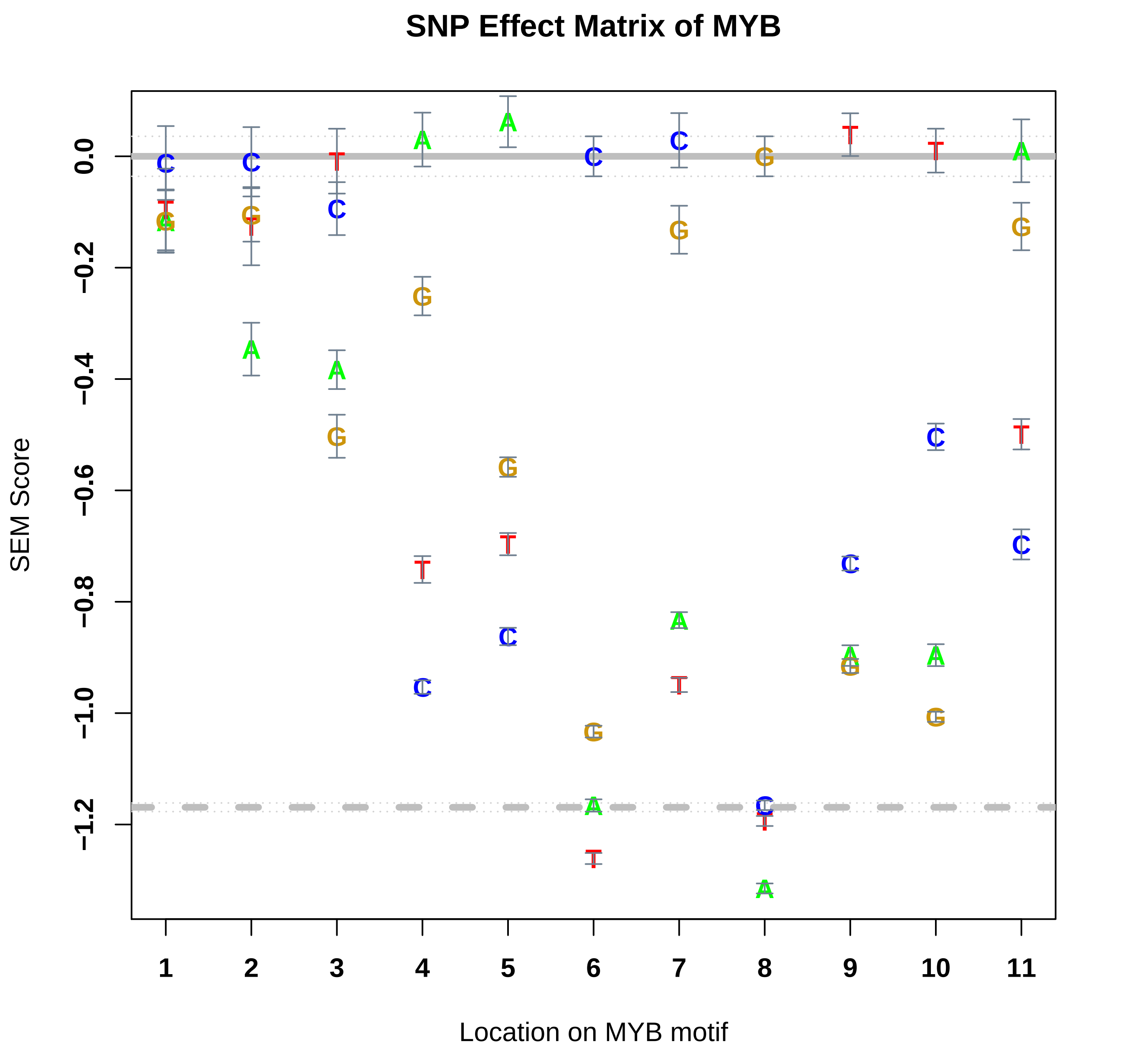

### M00776

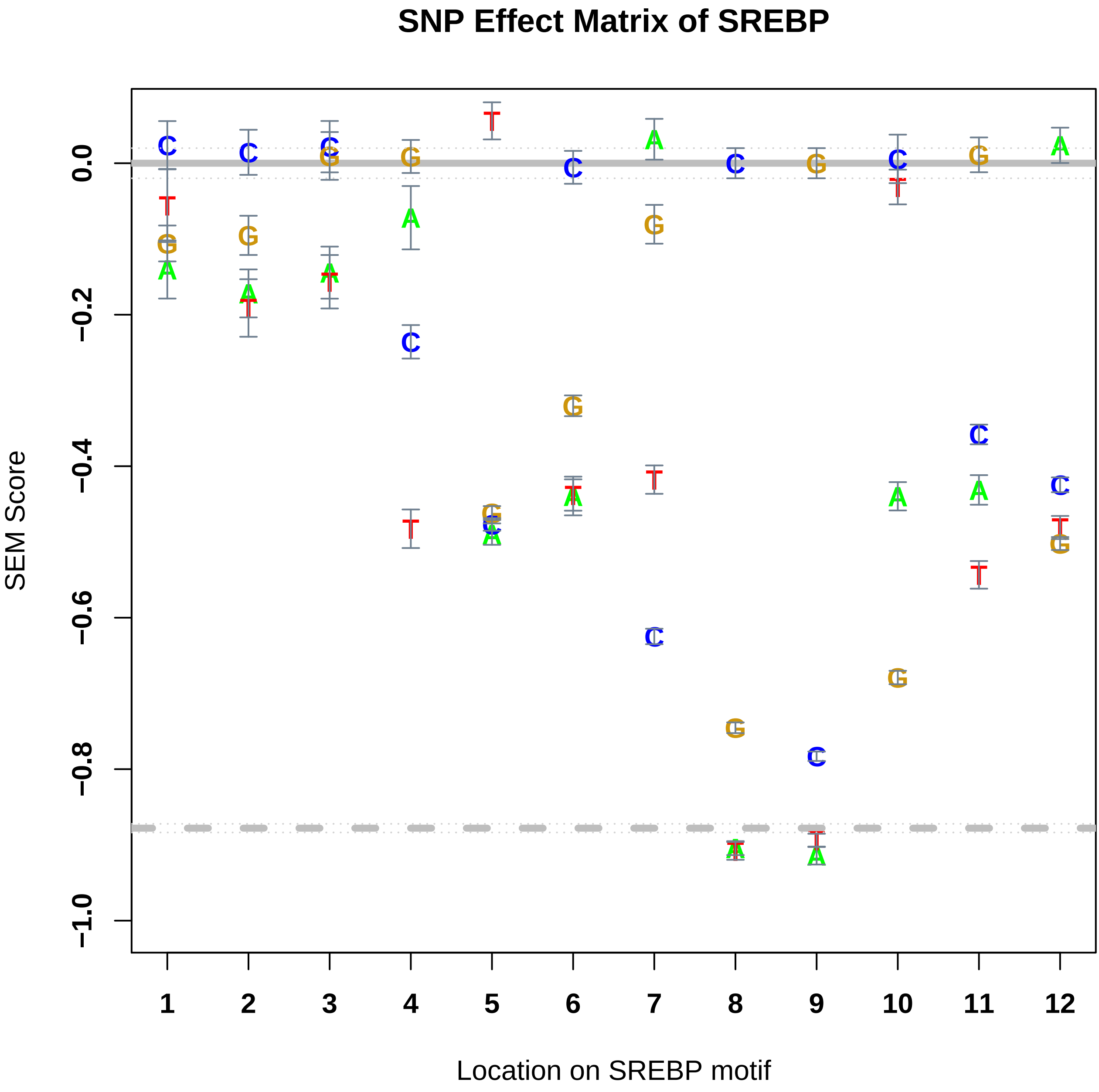

### M00778

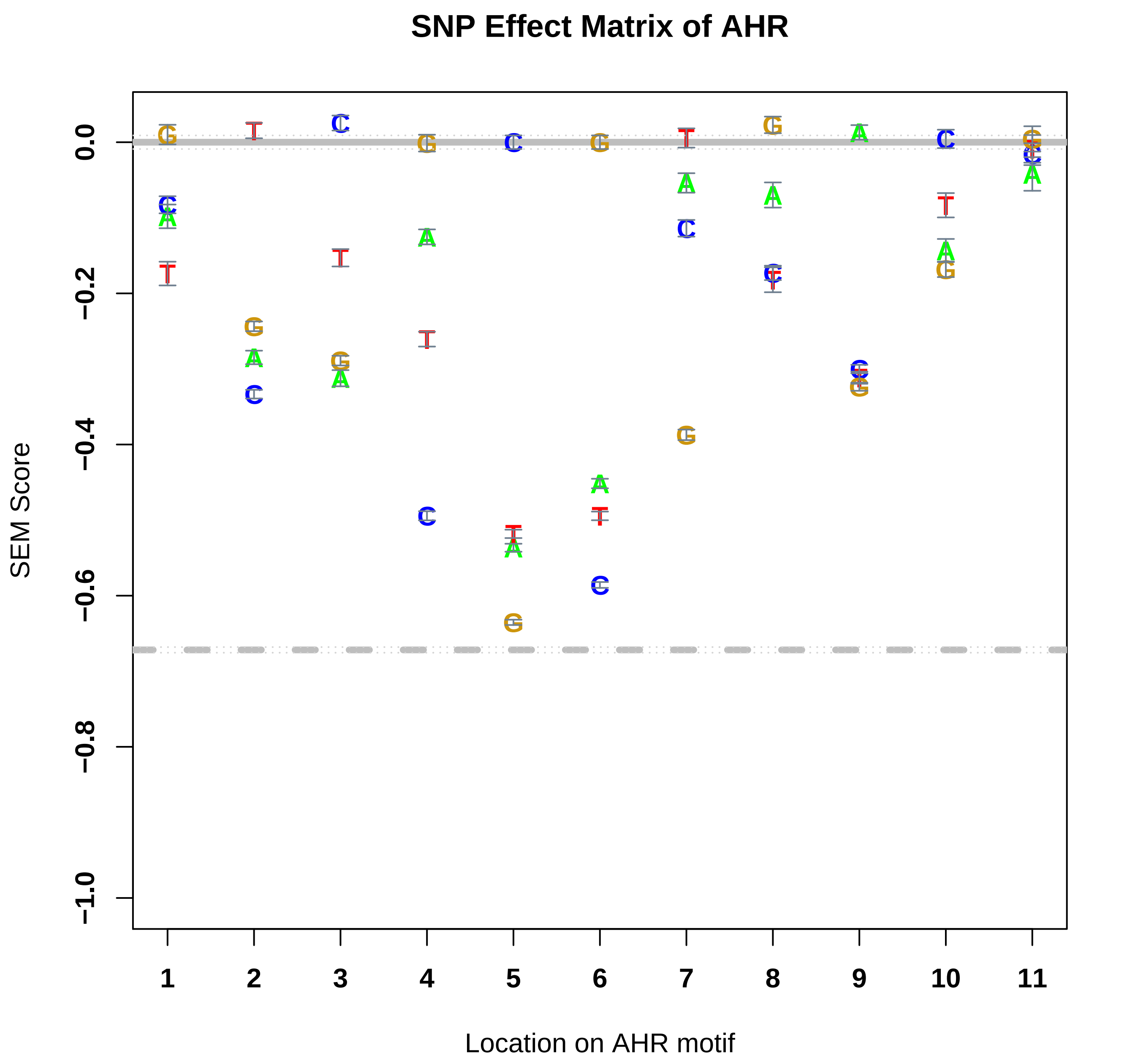

### M00789

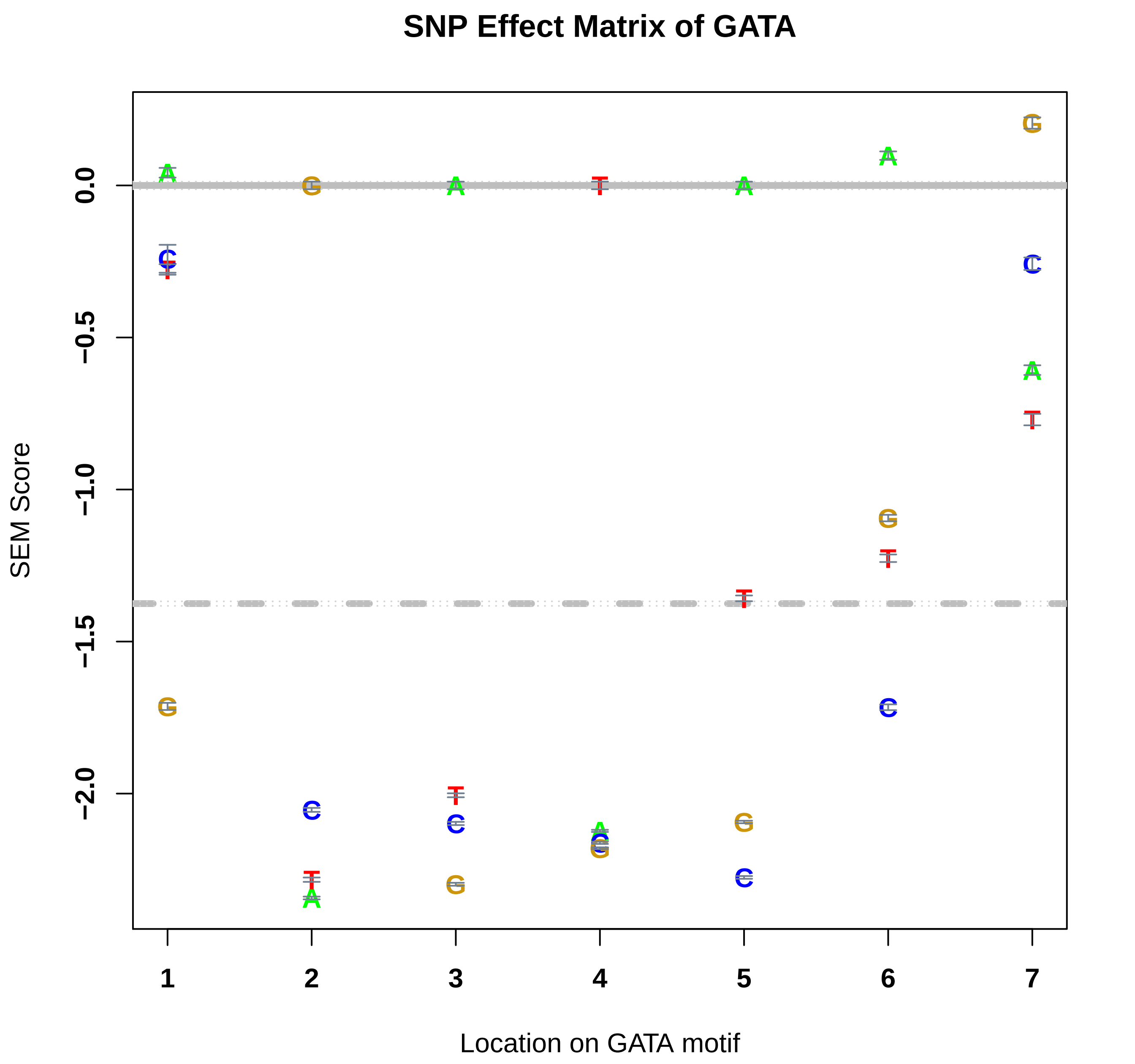

### M00792

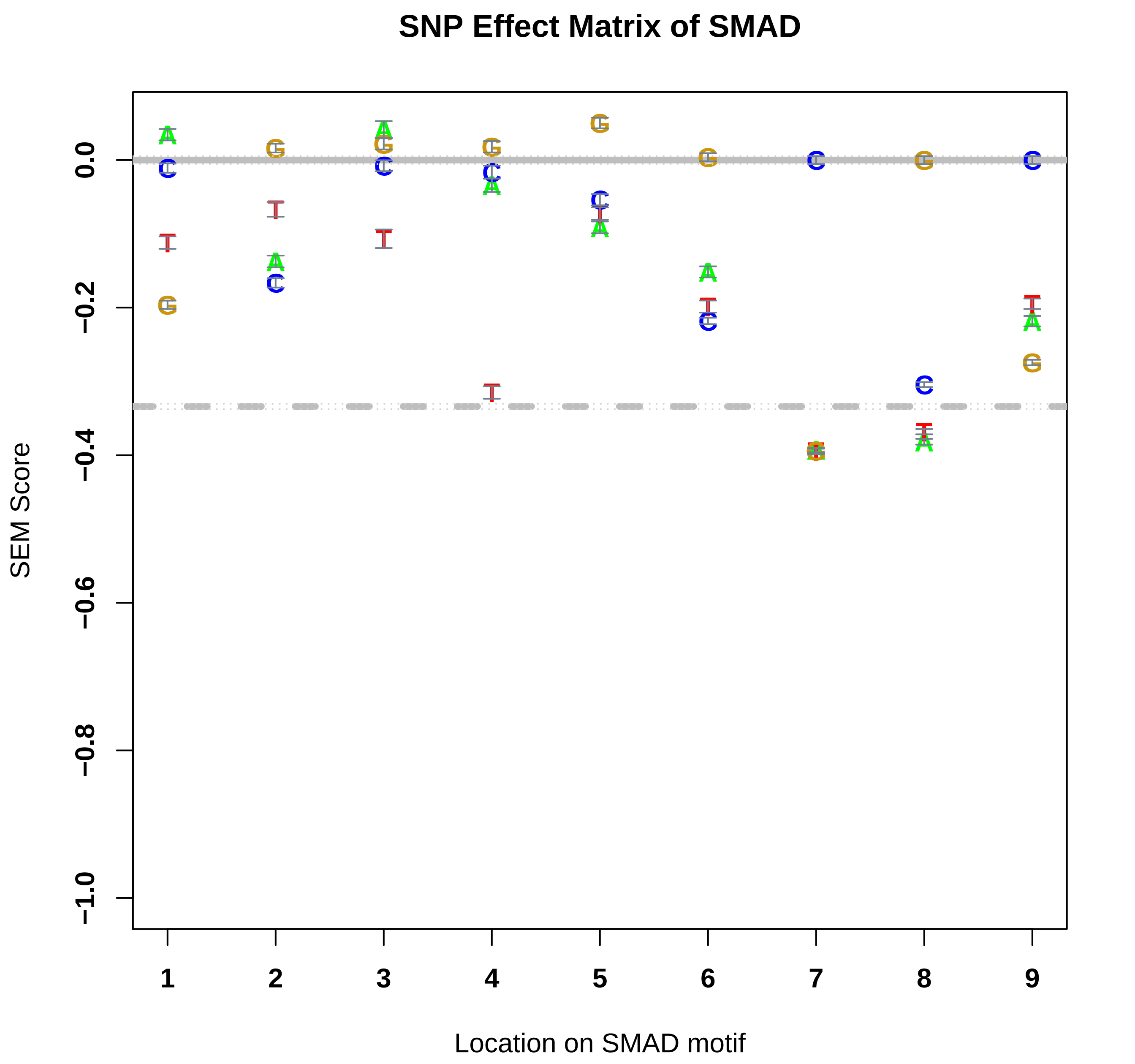

### M00793

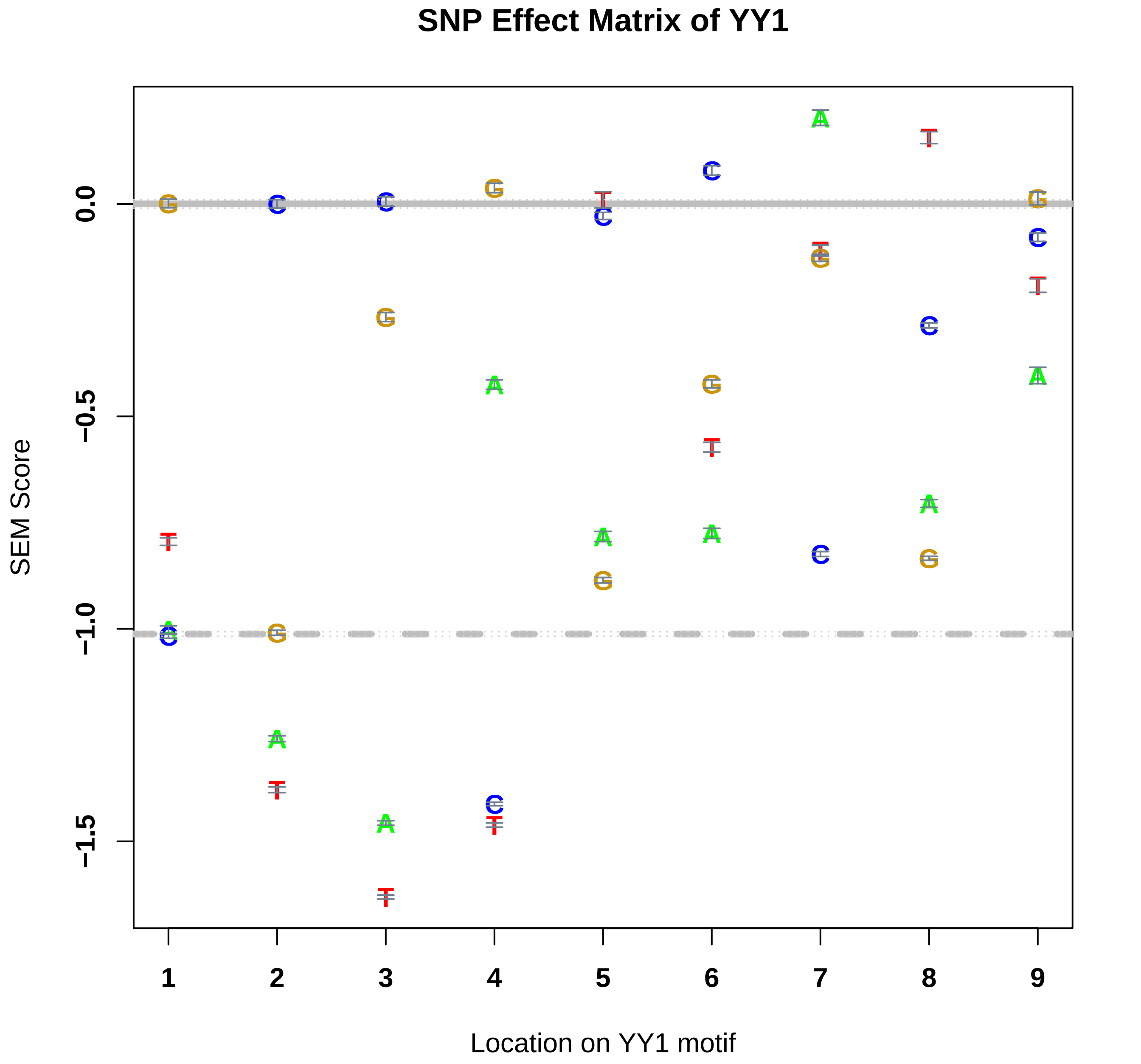

### M00799

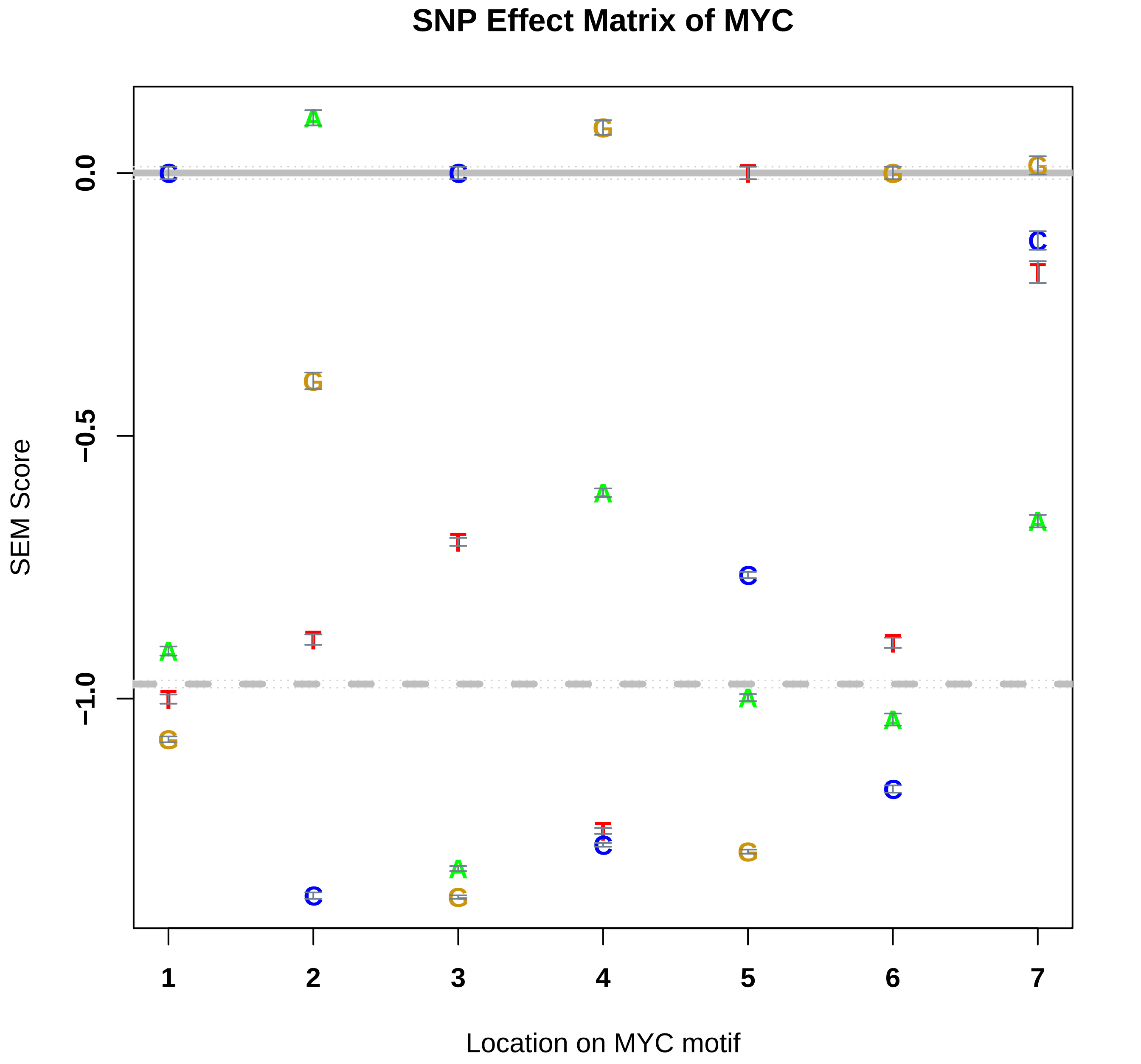

### M00801

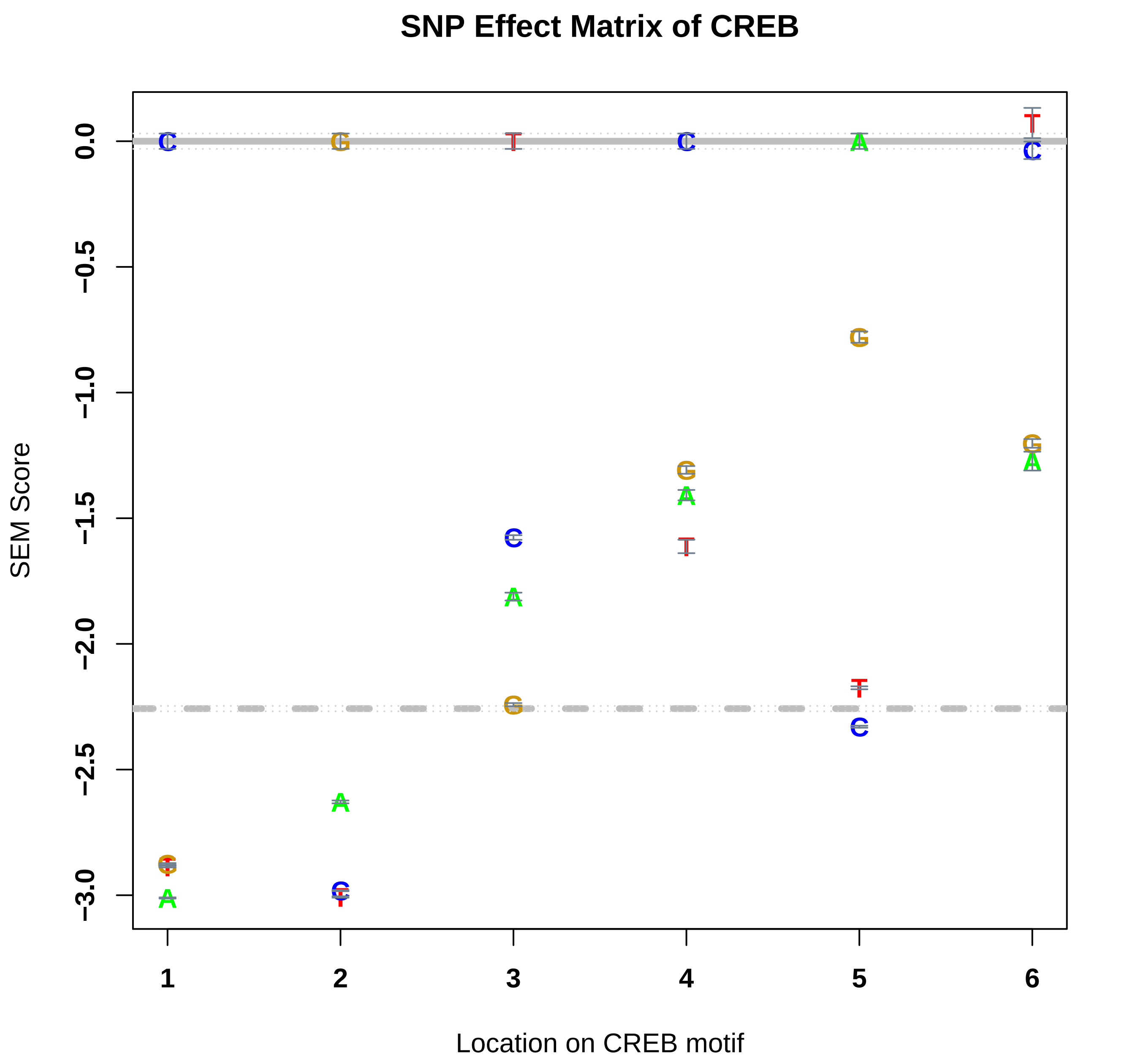

### M00803

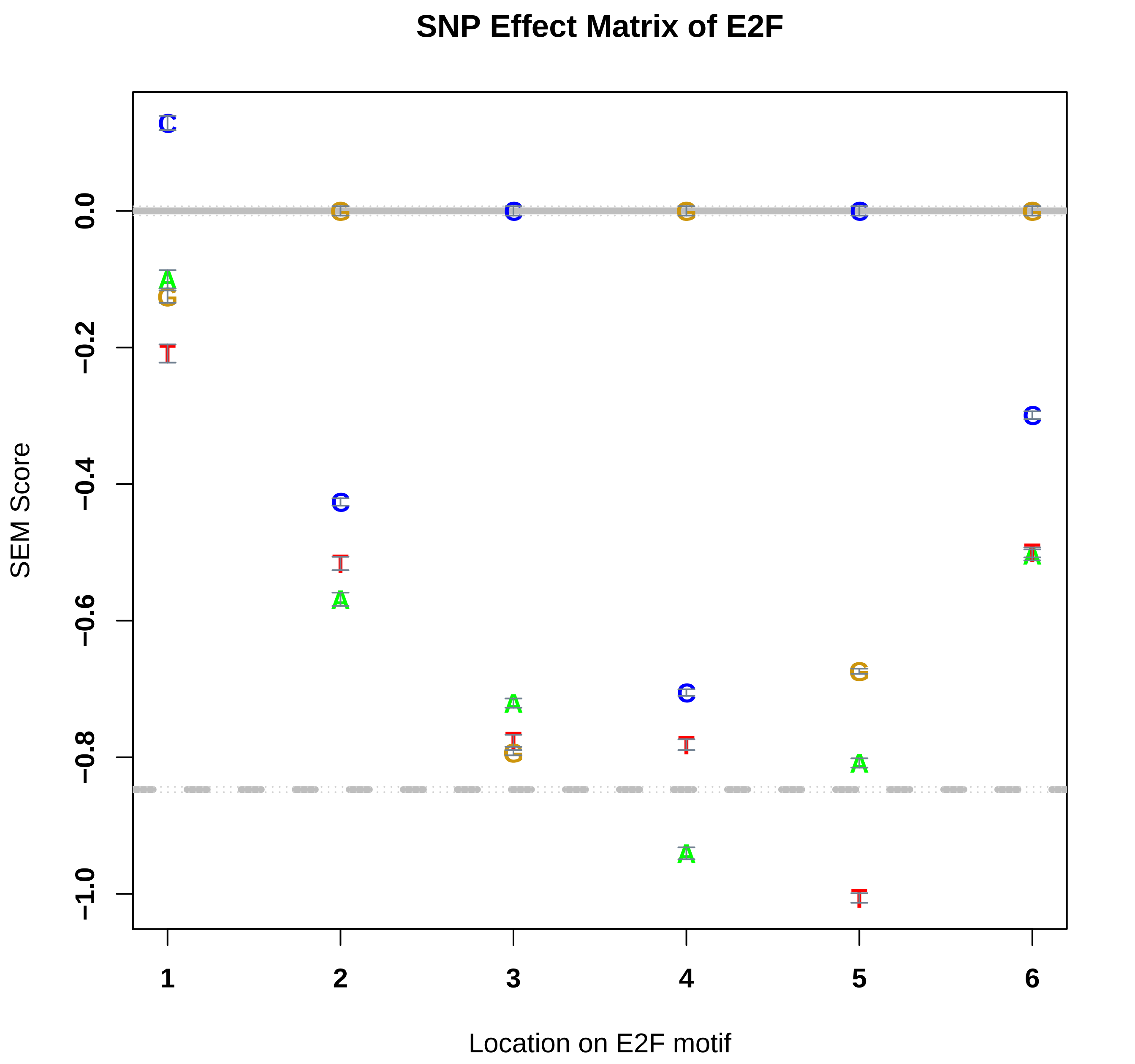

### M00913

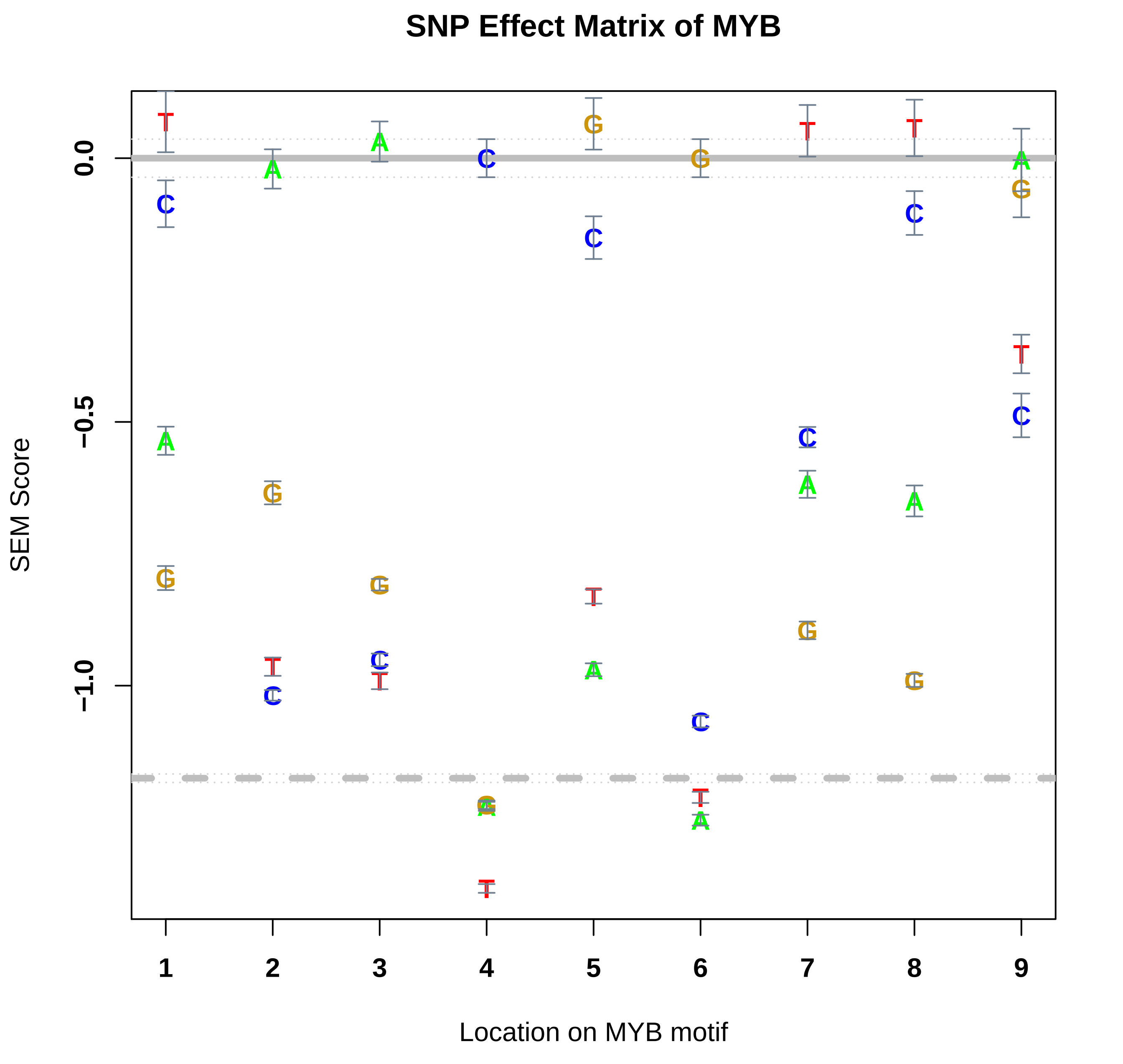

### M00915

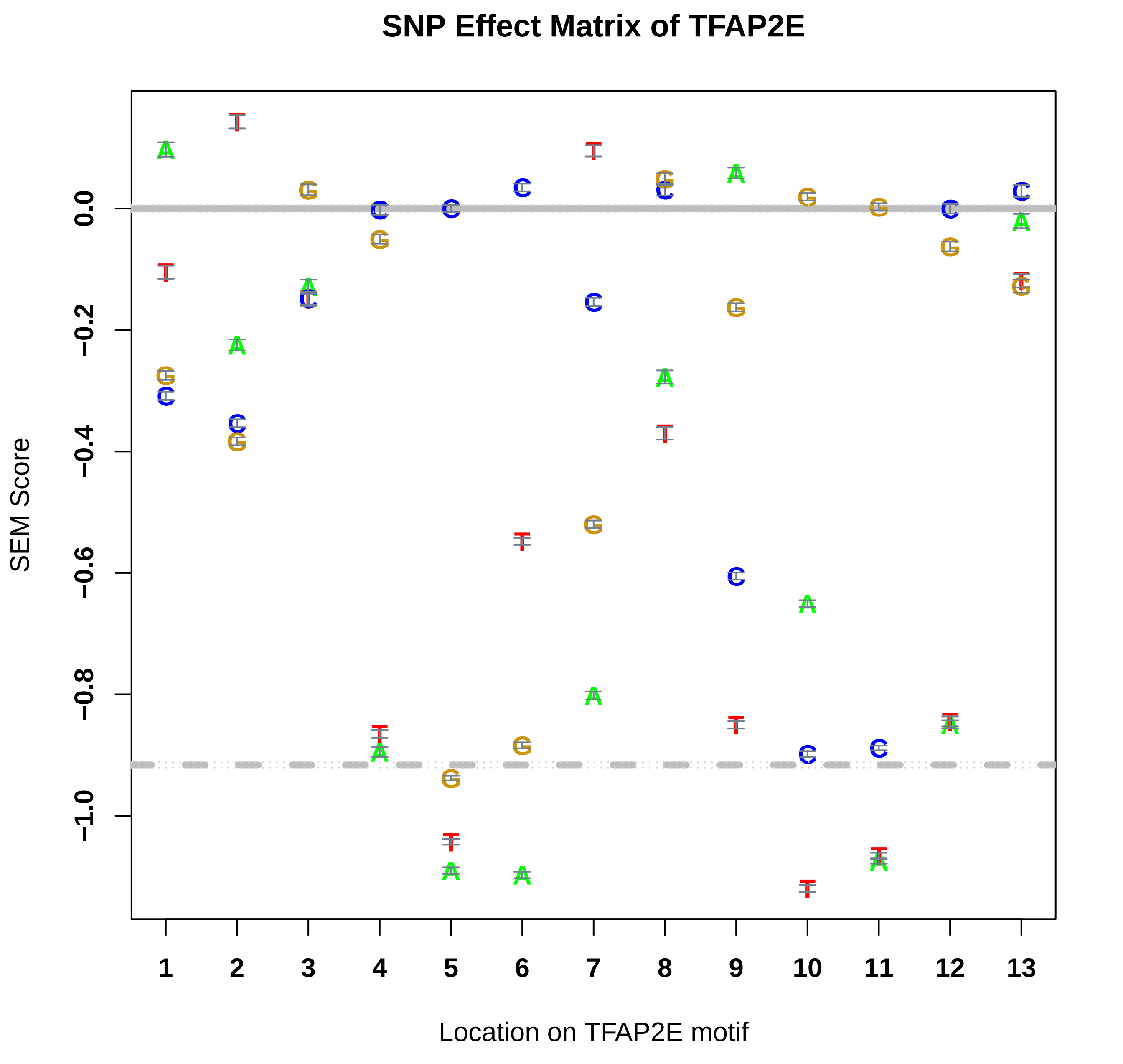

### M00917

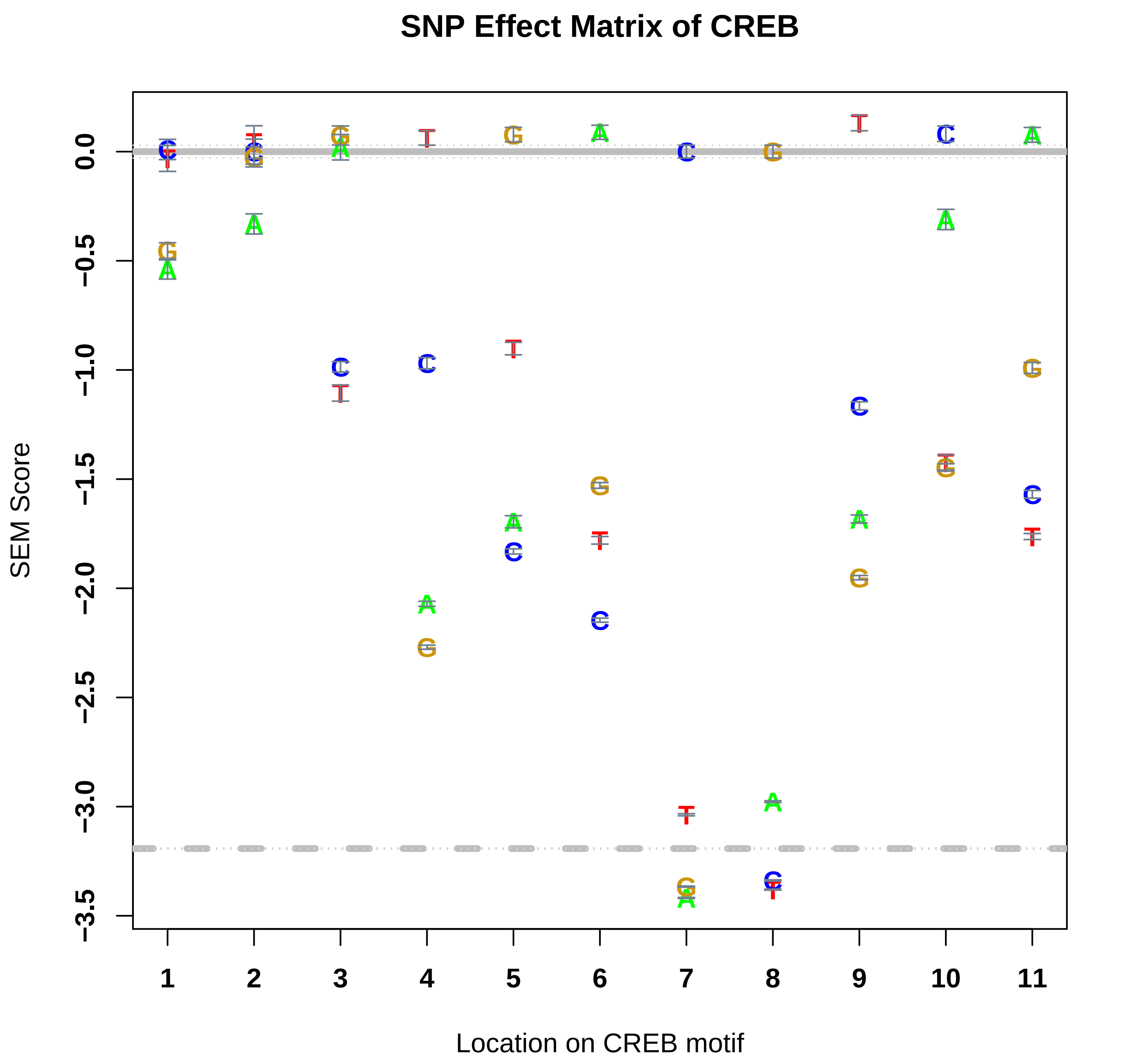

### M00918

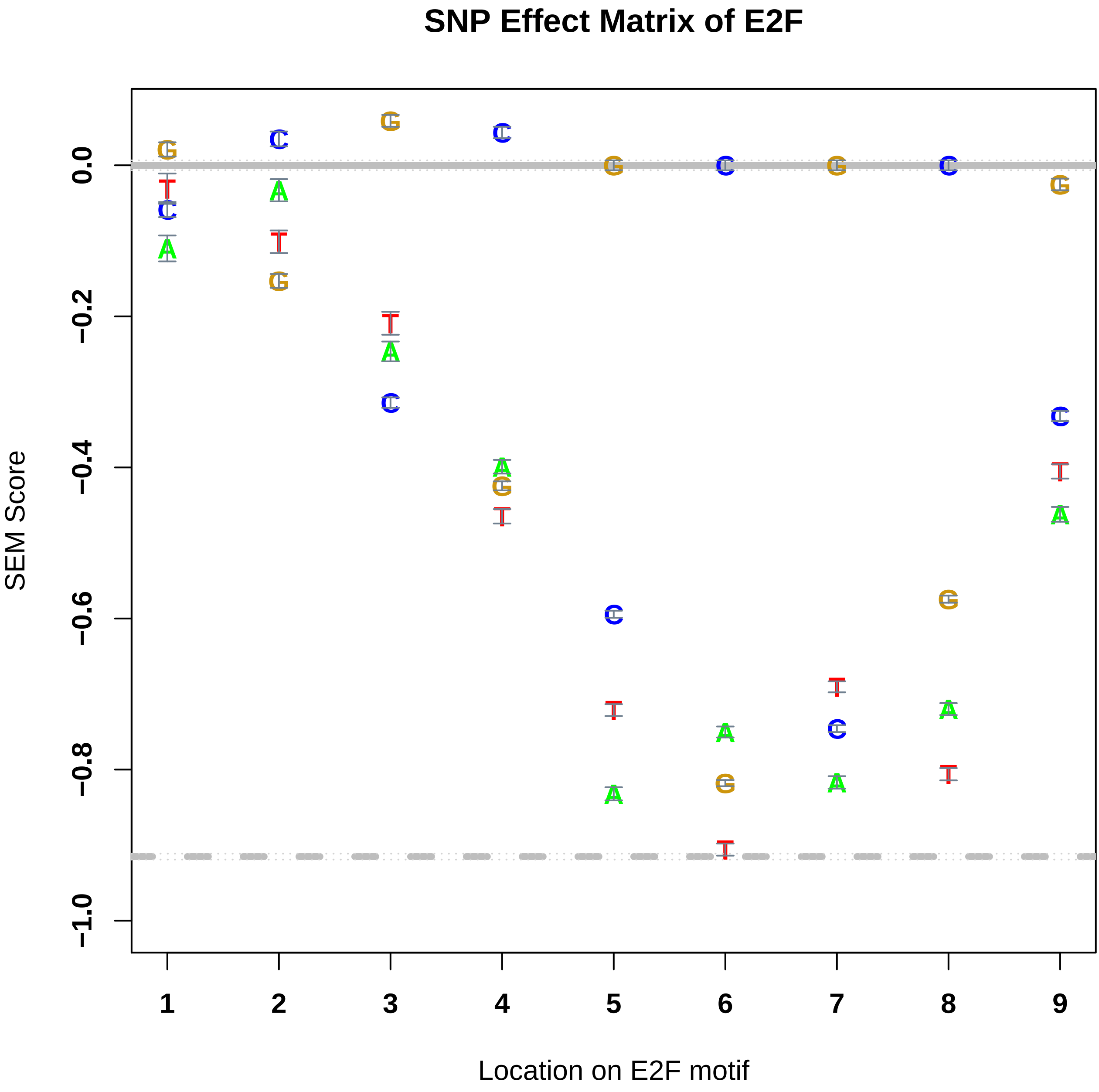

### M00919

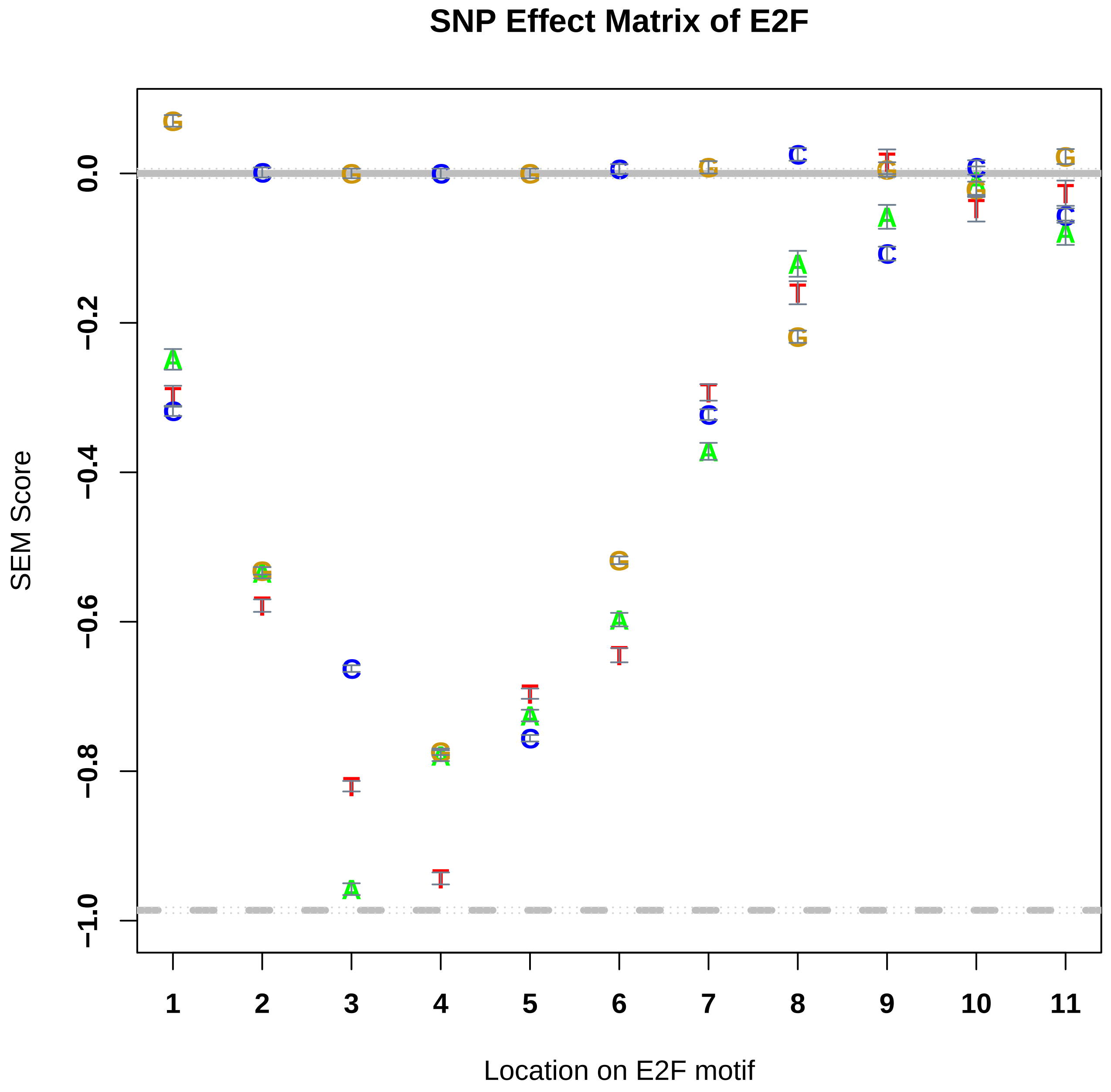

### M00921

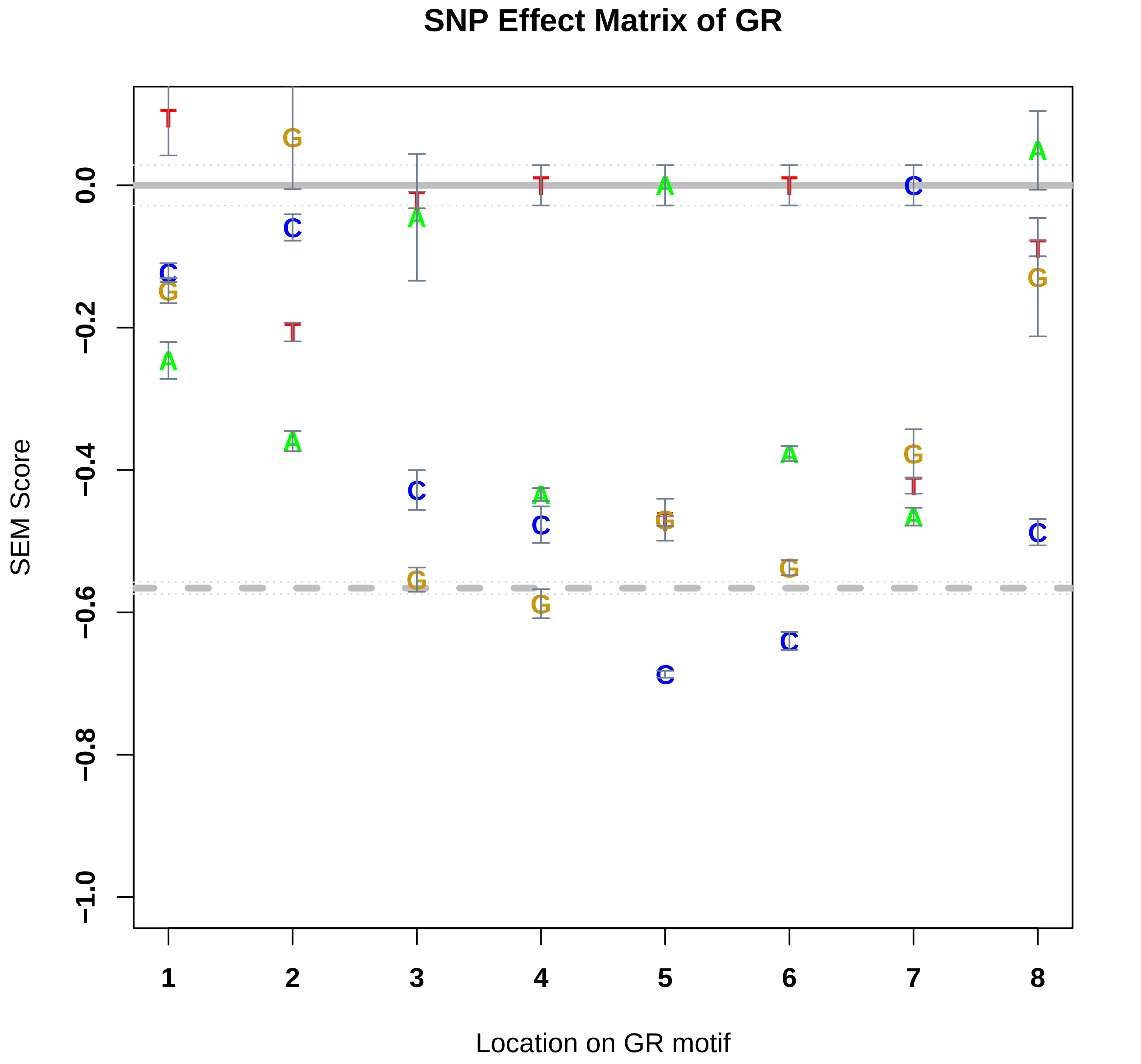

### M00925

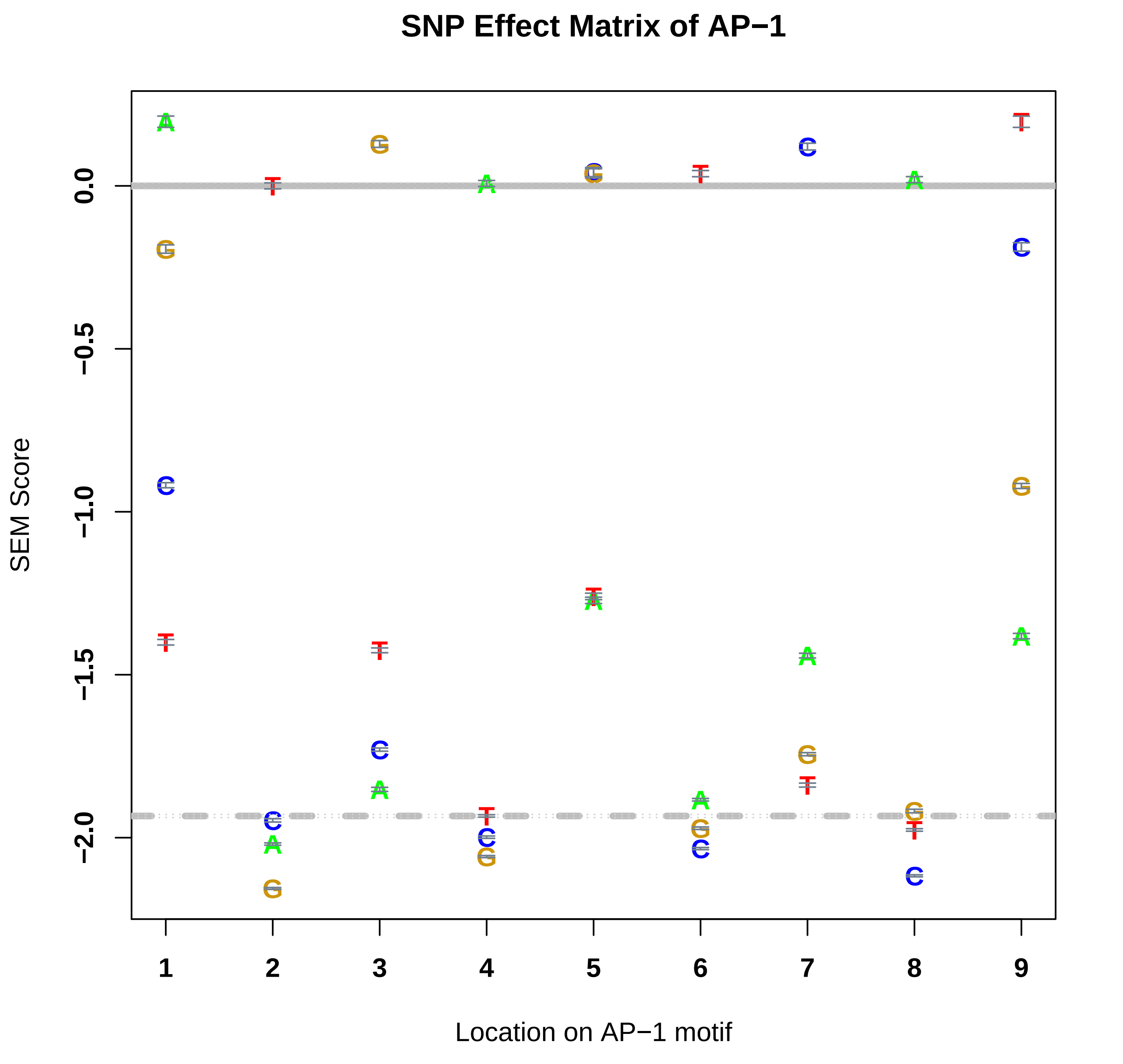

### M00926

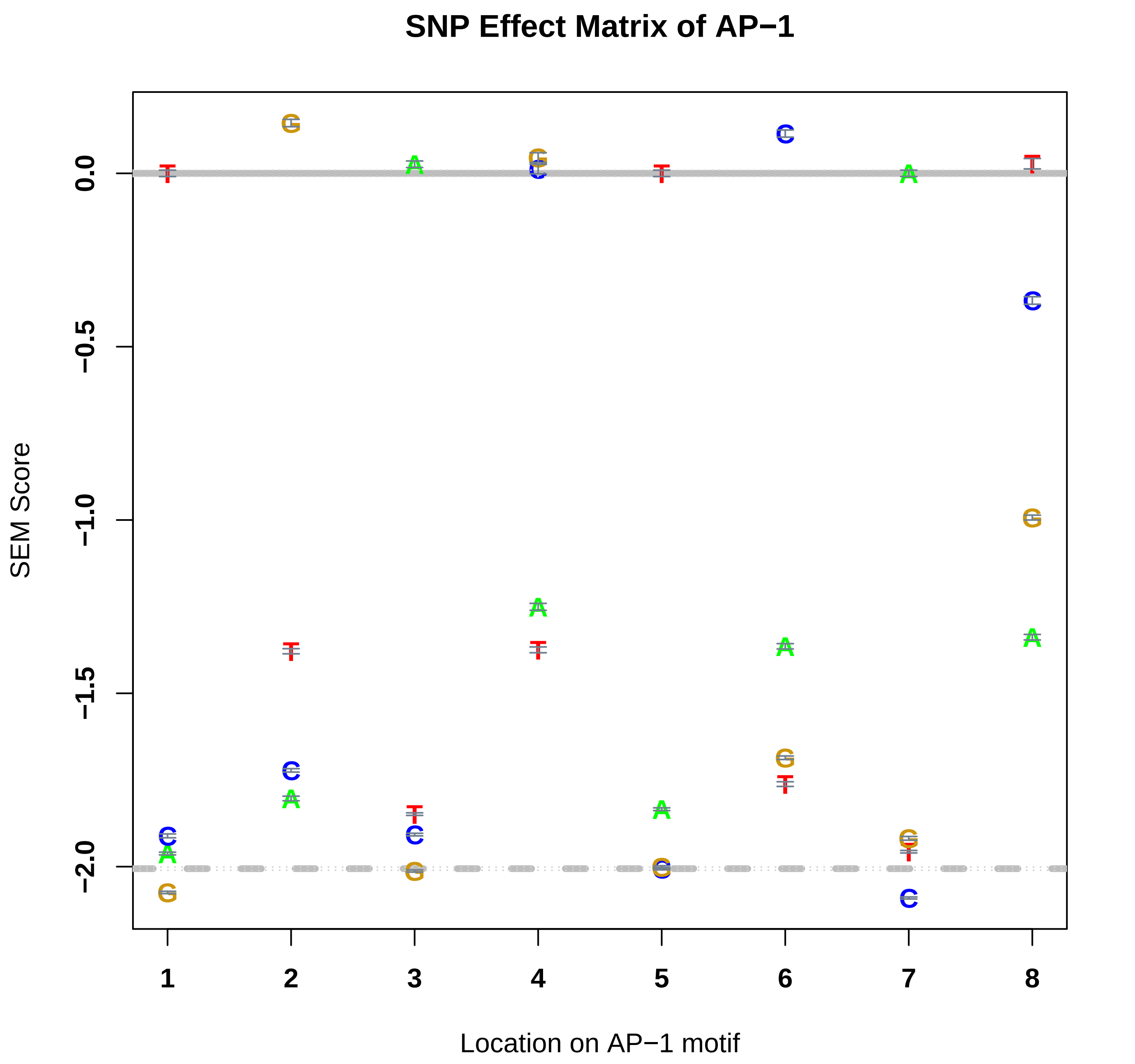

### M00931

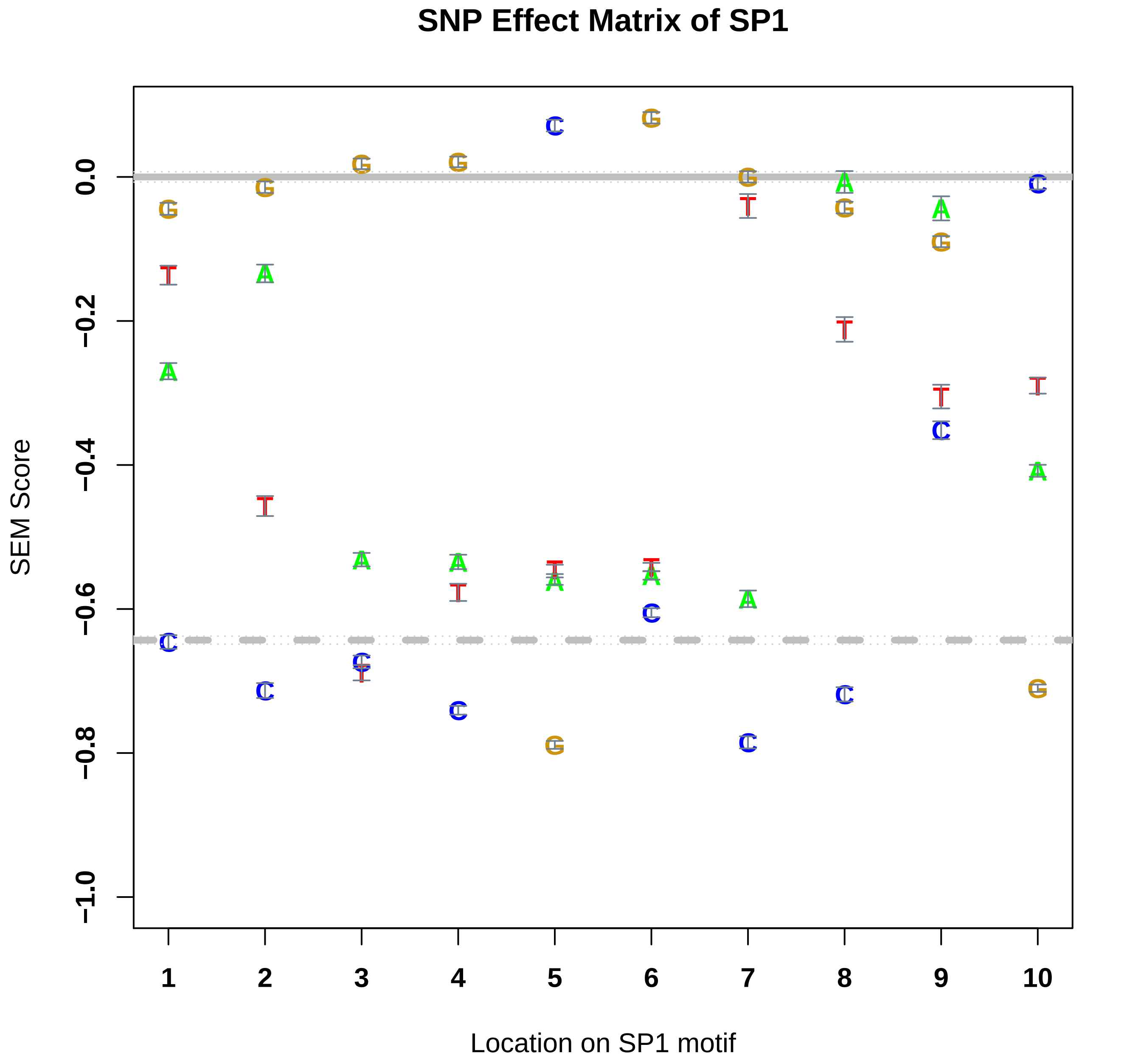

### M00932

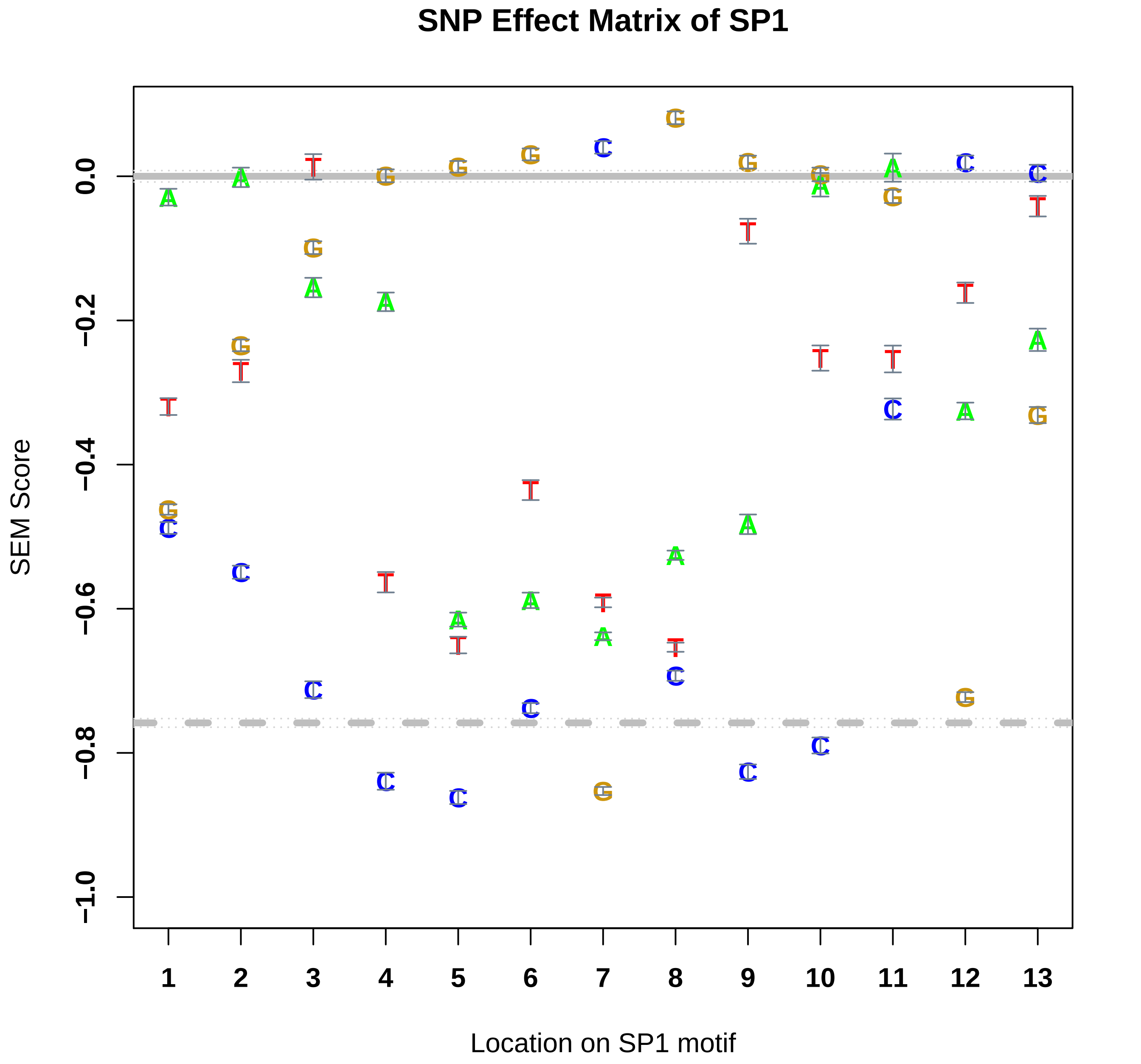

### M00933

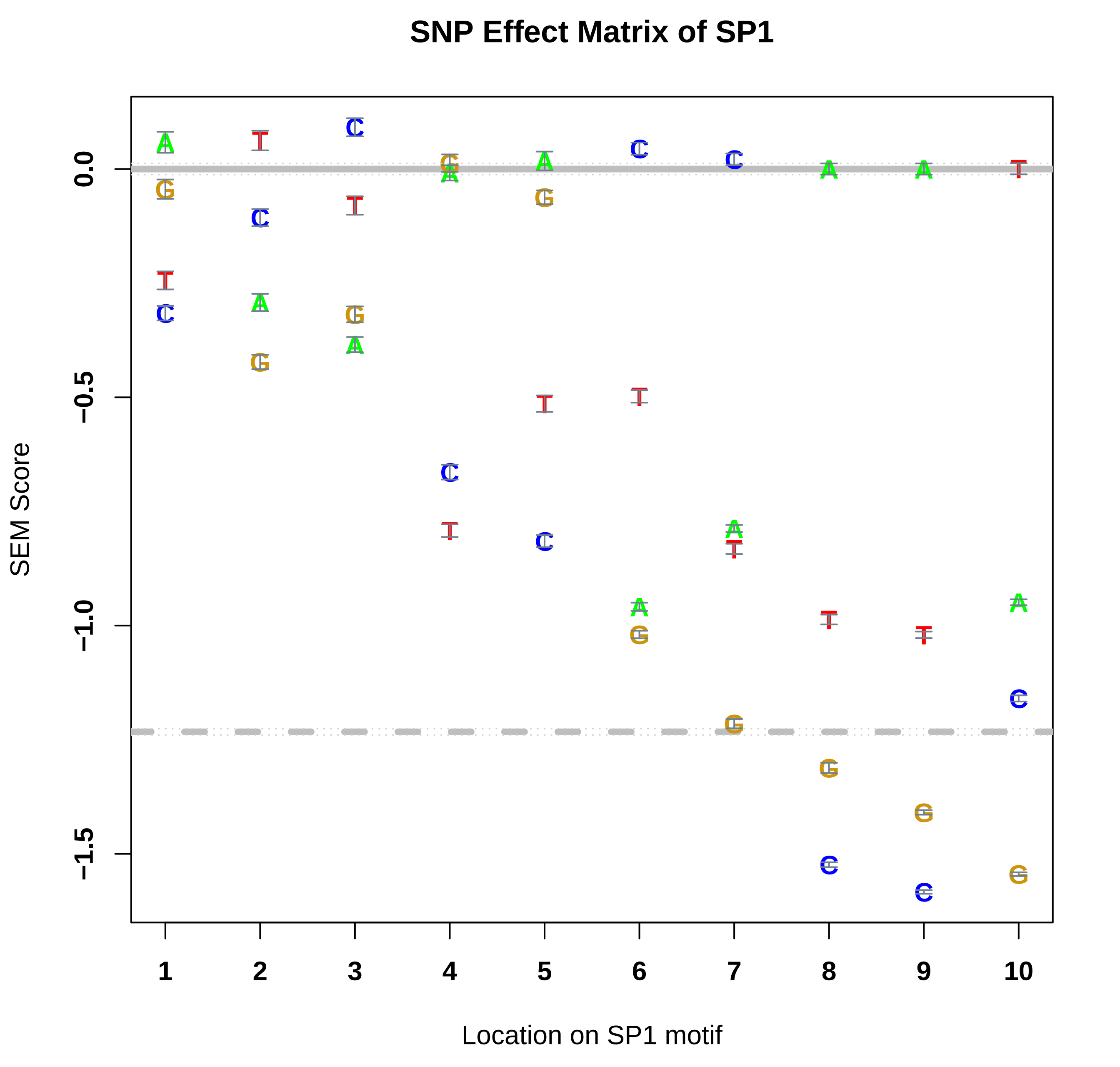

### M00939

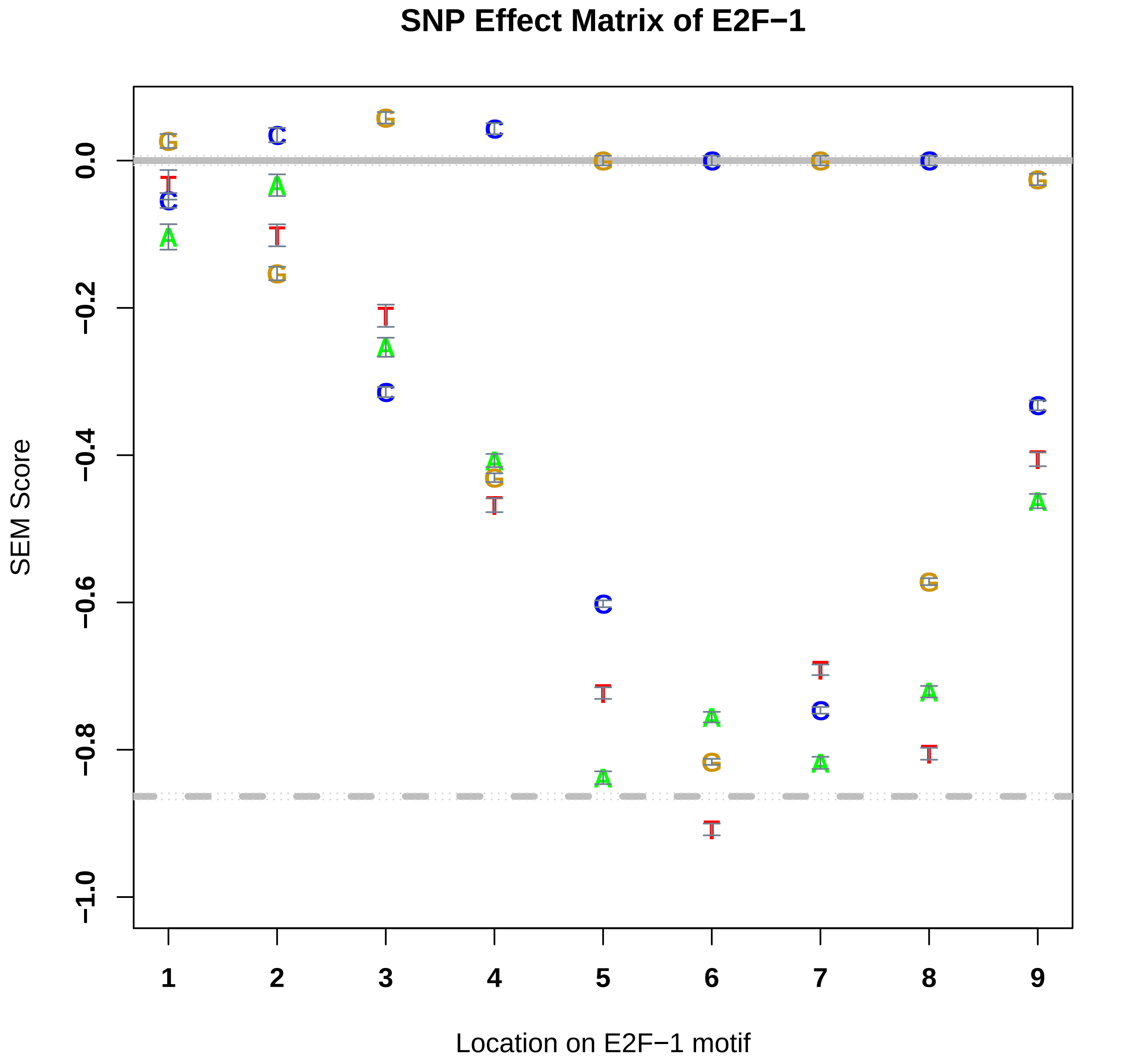

### M00940

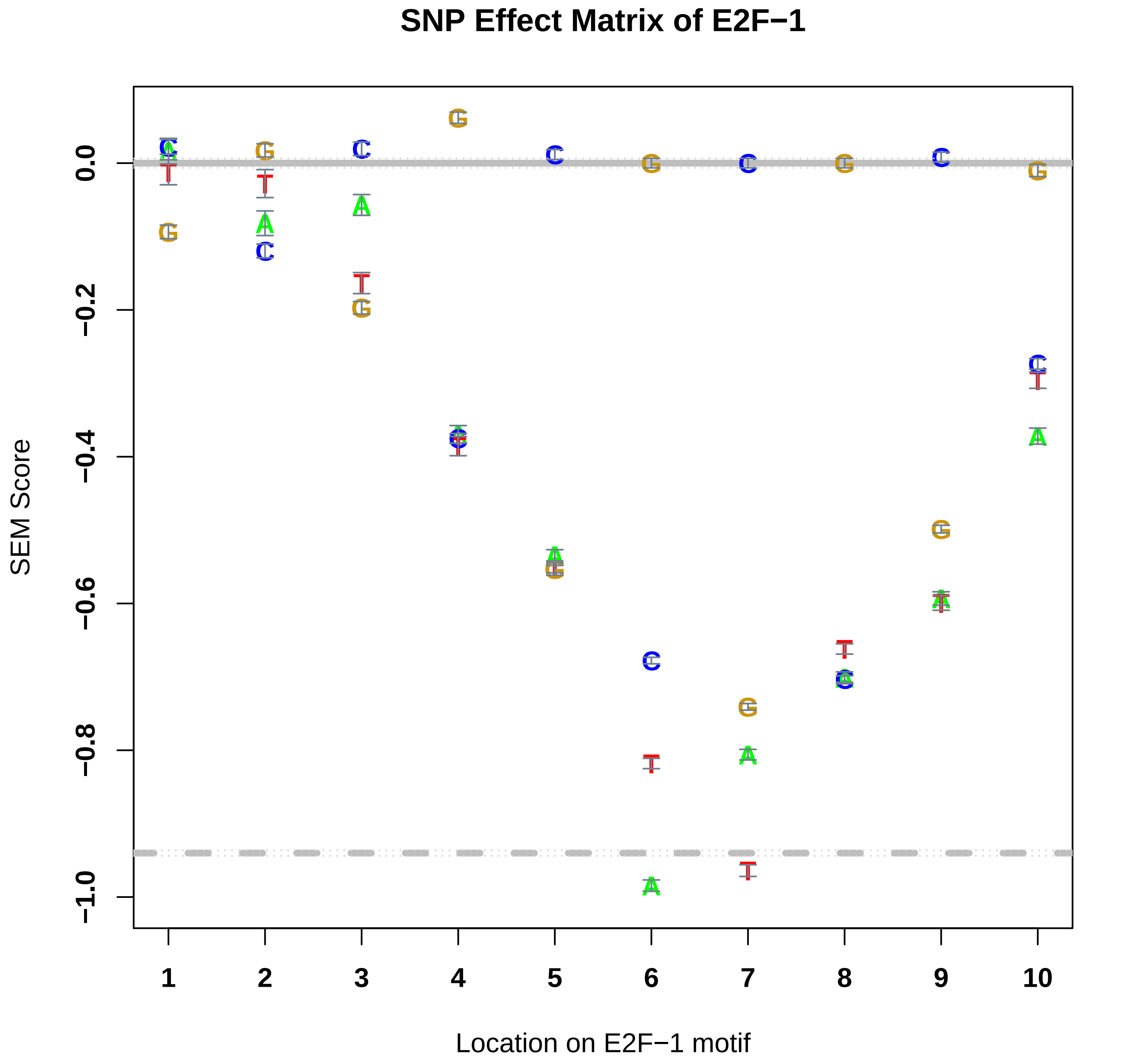

### M00967

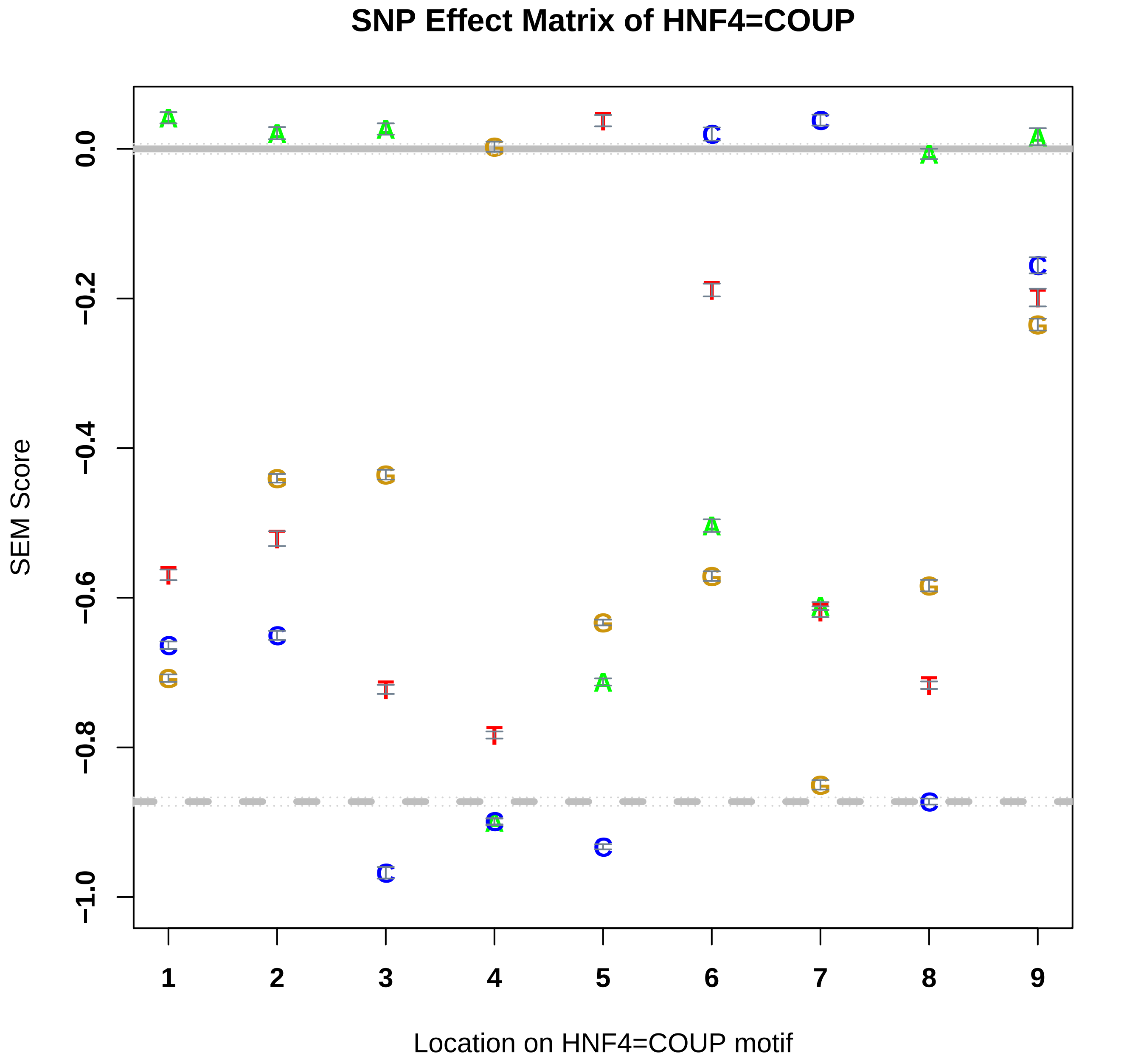

### M00974

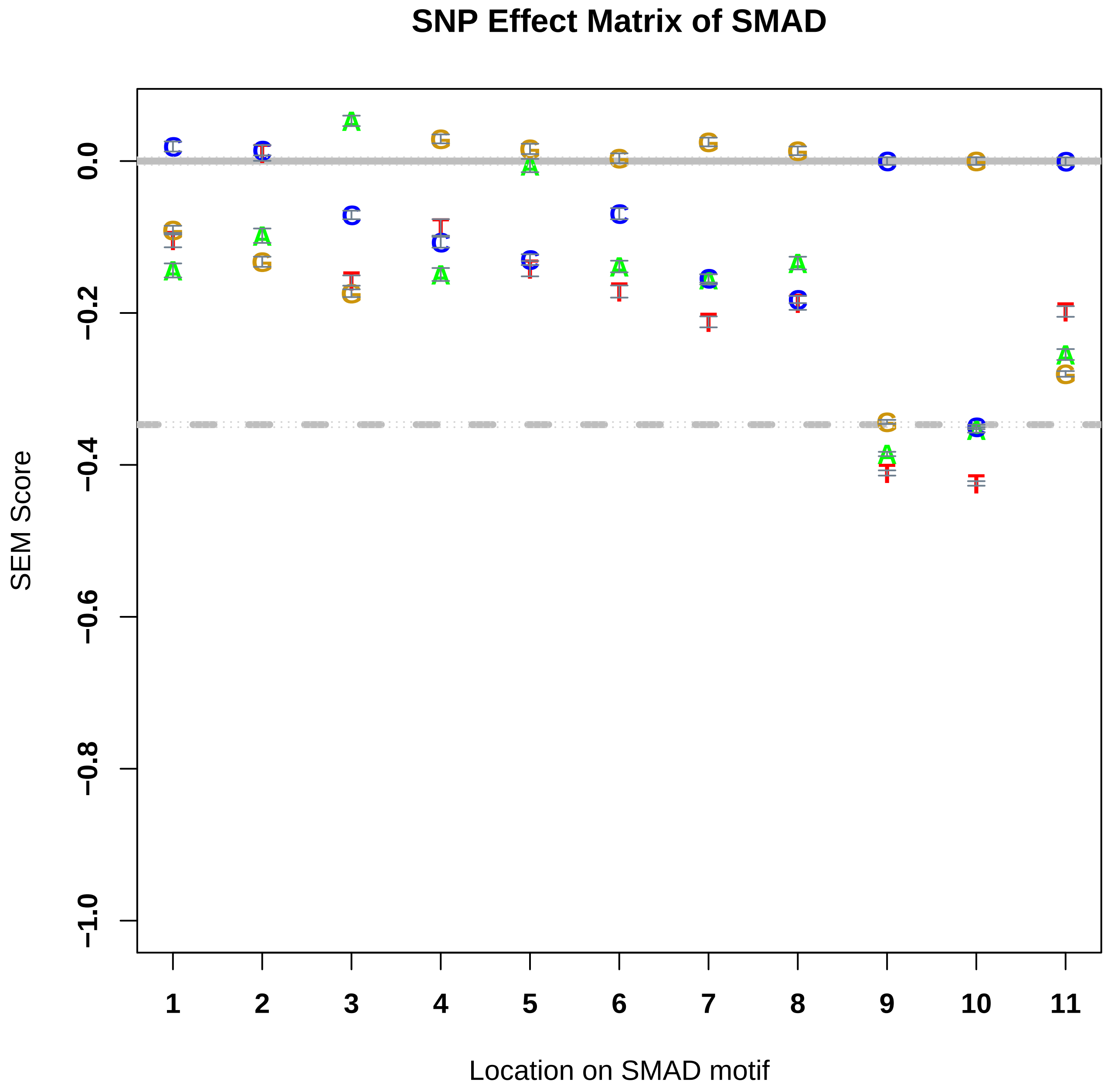

### M00976

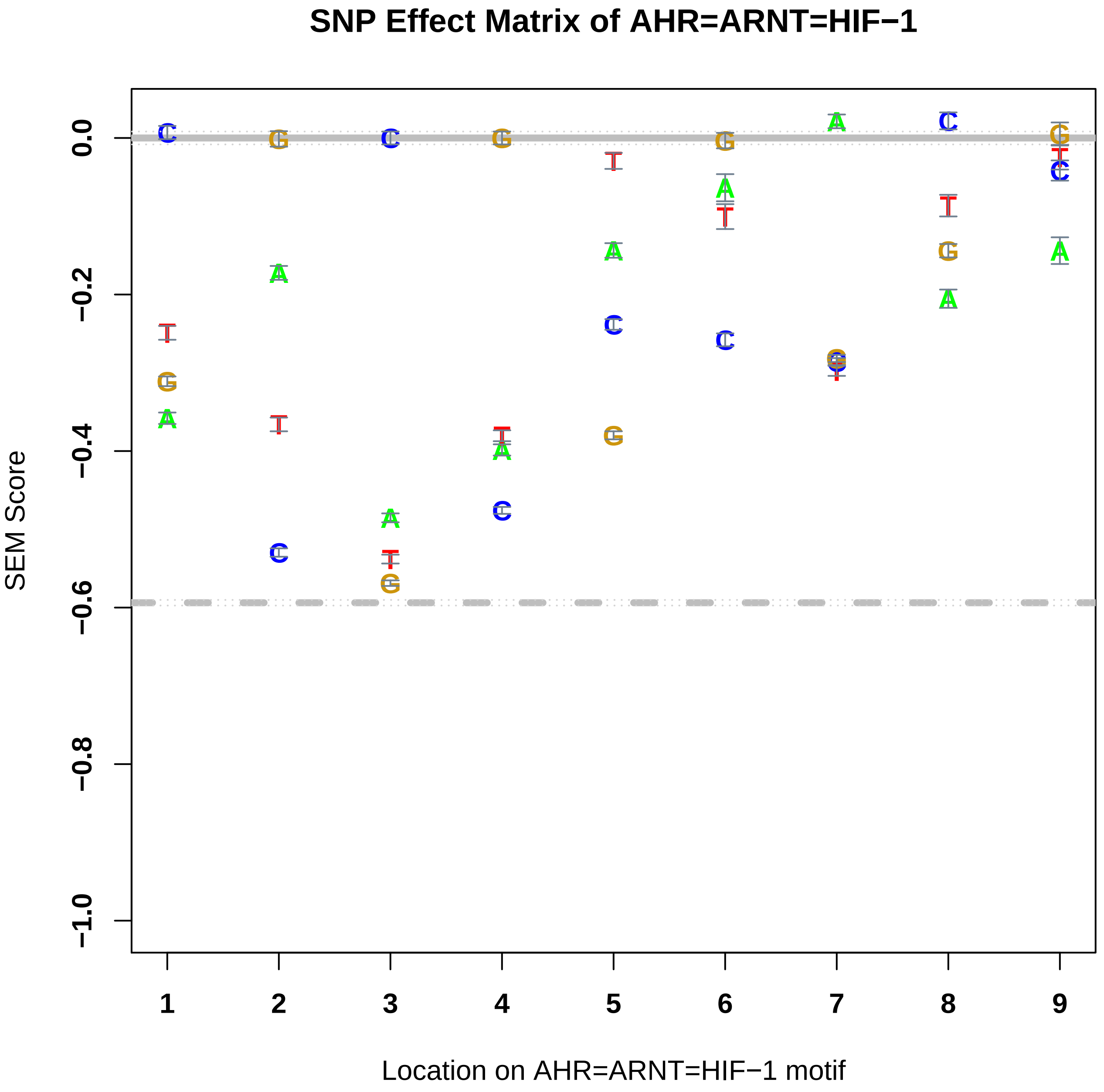

### M00977

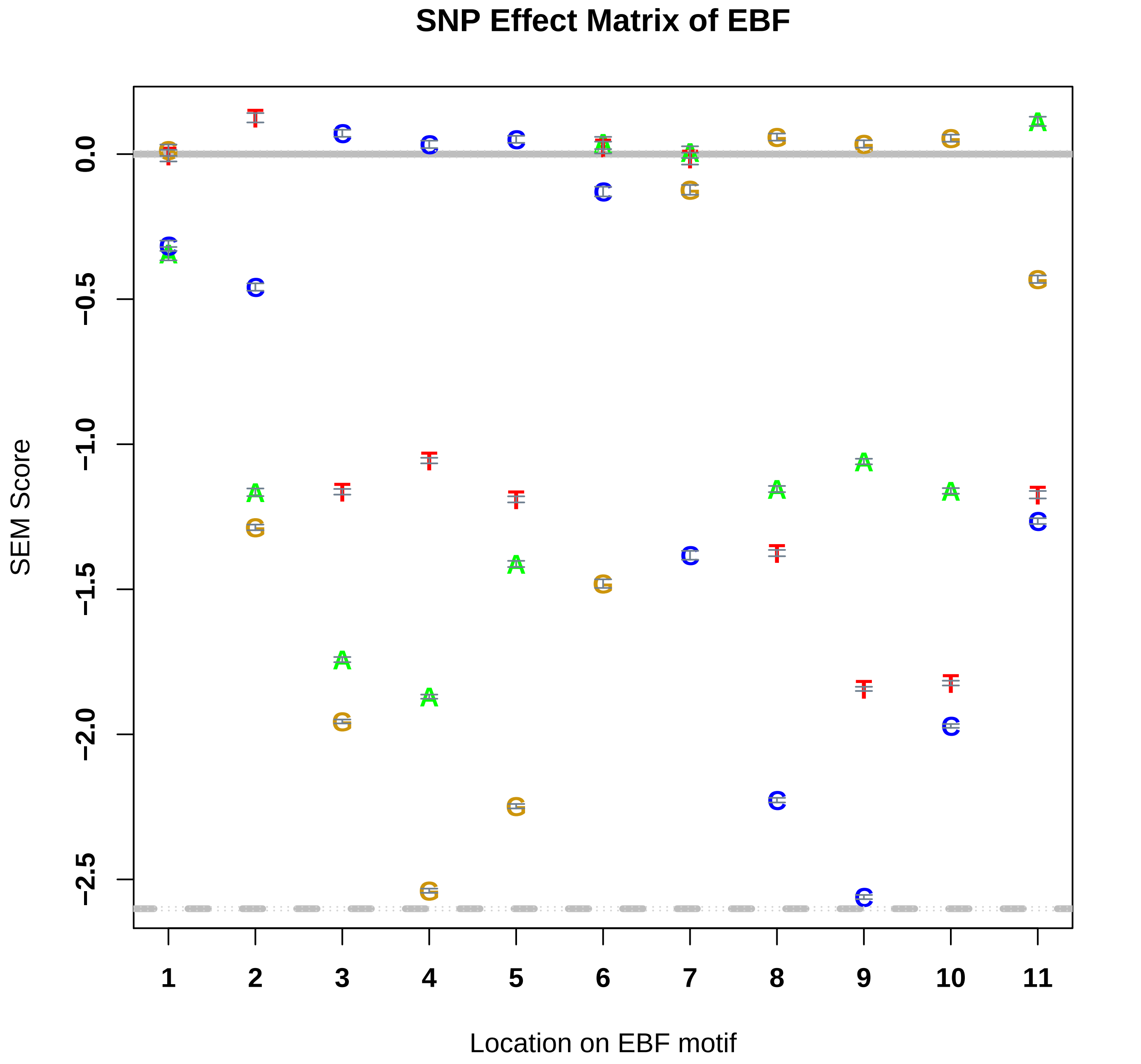

### M00981

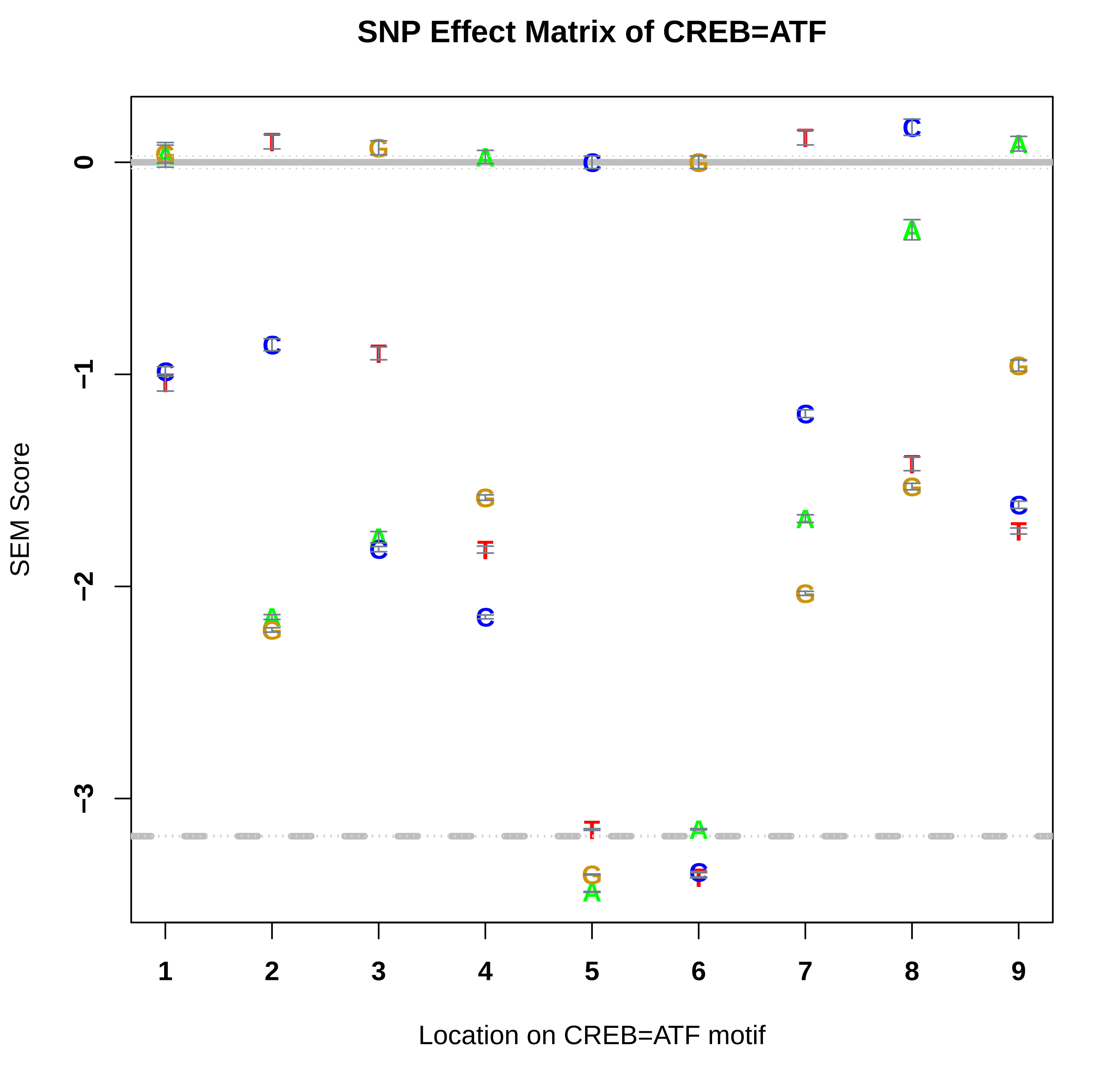
